# Brain-wide processing of gustatory information for gradient navigation in *Drosophila* larvae

**DOI:** 10.64898/2026.07.03.736052

**Authors:** Akhila Mudunuri, Katrin Vogt

## Abstract

Foraging in naturalistic environments is often challenging, as animals must evaluate varying sensory cues to locate optimal food sources. To successfully navigate a taste gradient, where perceived concentrations change over space and time, animals need to compare previously encountered taste qualities with current information. How such short-term taste memories are implemented in the brain and used to guide navigation remains poorly understood. Due to their powerful genetic toolkit and whole-brain connectome, *Drosophila* larvae are an excellent model organism for investigating the neural basis underlying taste gradient navigation. Using linear fructose and salt gradients, we show that larvae are attracted to high fructose concentrations, whereas they avoid high salt concentrations. To identify the neural basis underlying taste gradient navigation, we tested the role of cell types from the peripheral chemosensory system to higher brain regions. Contrary to conclusions from simple two-choice preference assays, we show that gradient navigation depends on different cell types across multiple layers of chemosensory processing and associative learning circuits, including mushroom body neurons. Attractive and aversive taste signals are conveyed through parallel, partially overlapping pathways that converge onto mushroom body output neurons, where they are integrated to shape motor behaviors during chemotaxis.

## Introduction

Animals can navigate complex, changing environments by integrating multiple sensory inputs to locate conditions that support growth and survival. Rather than relying on random exploration, many animals rely on directional information from environmental gradients to guide their movement towards beneficial resources. Successful gradient navigation can be computationally demanding, as it requires animals to compare sensory information across space and time while continuously updating behavioral decisions (Gomez-Marin & Louis, 2012; Schulze et al., 2015). Although the neural and behavioral mechanisms underlying olfactory (Louis et al., 2008; Kathman et al., 2024; Kramer et al., 2026), temperature (Luo et al., 2010; Klein et al., 2015; Balakrishnan & Haesemeyer, 2025; Eschbach et al., 2025) and light (Kane et al., 2013; Chen & Engert, 2014; Wolf et al., 2017) gradient navigation have been studied previously, far less is known about how animals navigate using gustatory cues (Luo et al., 2014; Dancausse et al., 2026).

During development, *Drosophila melanogaster* larvae need to feed continuously in order to reach the critical weight required for pupation (Bakker, 1959). In natural environments, larvae develop on decomposing plant and fungal substrates where nutrients are limited and unevenly distributed (Markow & O’Grady, 2008), requiring them to use sensory information to identify and navigate toward favorable food sources. The available extensive genetic toolkit, together with the whole-brain connectome (Winding et al., 2023), makes *Drosophila melanogaster* larvae a powerful system for investigating the neural circuits underlying taste gradient navigation.

Larvae respond to different gustatory cues, such as sugar or salt, with either attraction or avoidance, respectively. Previous studies of gustatory behavior have relied on a two-choice preference assay in which the tastant is present against pure agar. They showed that fructose is attractive in a concentration-dependent manner (Miyakawa et al., 1980; Schipanski et al., 2008; Rohwedder et al., 2012; Mishra et al., 2013), whereas salt is aversive at high concentrations but attractive at low concentrations (Liu et al., 2003; Niewalda et al., 2008; Zhang et al., 2013; Alves et al., 2014). Gustatory stimuli have also been widely used as reinforcers in learning and memory paradigms; however, innate fructose attraction and salt avoidance tested in the two-choice assay are preserved even when mushroom body circuits involved in learning were silenced (Pauls et al., 2010; Rohwedder et al., 2016; Widmann et al., 2016; Saumweber et al., 2018; Mancini et al., 2023; Weber et al., 2025).

Taste gradient navigation can require more than a simple stimulus response. To navigate effectively, larvae must retain information about previously encountered taste conditions, compare it with current sensory input, and adapt their movements accordingly. *Drosophila* larvae exhibit stereotyped movement behaviors consisting of forward peristaltic movement (runs) and stops, during which they perform head sweeps followed by a turning event (Riedl & Louis, 2012). During successful gradient navigation, larvae increase run length and reduce turning frequency and angle as they move toward favorable regions (Luo et al., 2010; Gomez-Marin et al., 2011; Eschbach et al., 2025). Inspired by salt gradient navigation assays in *C. elegans* (Luo et al., 2014) and nicotine gradient navigation assay in *D. melanogaster* larvae (Dancausse et al., 2026), we established a linear taste gradient assay for *Drosophila* larvae. We tested larvae individually to remove any social modulation via conspecifics (Mudunuri et al., 2026). Using this assay, we show that larvae navigate fructose gradients toward high concentrations, while in salt gradients, they avoid high salt regions and prefer to move toward lower concentrations.

Although the whole brain connectome is available, how gustatory information is processed and transformed into navigational behavior remains poorly understood. Gustatory cues are detected by chemosensory organs present on the larval head and the gustatory sensory neurons project to the subesophageal zone (SEZ), the primary taste processing center in the larvae (Vosshall & Stocker, 2007; Apostolopoulou et al., 2015; Berck et al., 2016; Rist & Thum, 2017; Miroschnikow et al., 2018; Maier et al., 2021; Komarov et al., 2025; Richter et al., 2025). However, how sensory neurons convey taste information into the SEZ and how projection pathways relay SEZ output to higher-order brain regions remains functionally uncharacterized. Connectomic analysis of the olfactory processing pathway has shown that these neurons additionally receive non-olfactory sensory input (Berck et al., 2016; Miroschnikow et al., 2018), yet how gustatory signals are incorporated into these circuits to guide behavior remains unclear. Moreover, gustatory valence signals are represented in mushroom body input neurons, mainly the dopaminergic neurons (Thum & Gerber, 2019; Weber et al., 2025), which connect to Kenyon Cells (KCs) and converge onto mushroom body output neurons, such as MBON-m1, that bias larval navigational decisions (Eschbach et al., 2025).

To investigate the neural basis of taste gradient navigation, we examined the functional role of multiple cell types identified through connectomic analysis, from the chemosensory periphery to higher order brain regions to mushroom body output neurons. Our results show that fructose and salt are detected by different peripheral receptors, and their information is conveyed to the higher brain by distinct but partially overlapping parallel pathways that ultimately drive fructose attraction and salt avoidance. We also find requirements for MB circuit cell types, expanding the role of this neuropil for gustatory innate behavior.

## Results

### *Drosophila* larvae navigate fructose and salt gradients using Gr43a and Ir76b

To investigate larval navigation on taste gradients, we created linear gradients of fructose and salt (0 - 1M) mixed with agar in a 25 x 25 cm square arena (Fig. 1A). Individual early third-instar larvae were placed at the center, and their behavior was recorded for 15 minutes at 1 fps under infrared illumination. Larvae moved randomly on pure agar plates, but successfully navigated both 1M gradients, accumulating at high, attractive fructose concentrations and avoiding high, aversive salt concentrations (Fig. 1B, C, Movie S1-S3).

**Fig 1.**
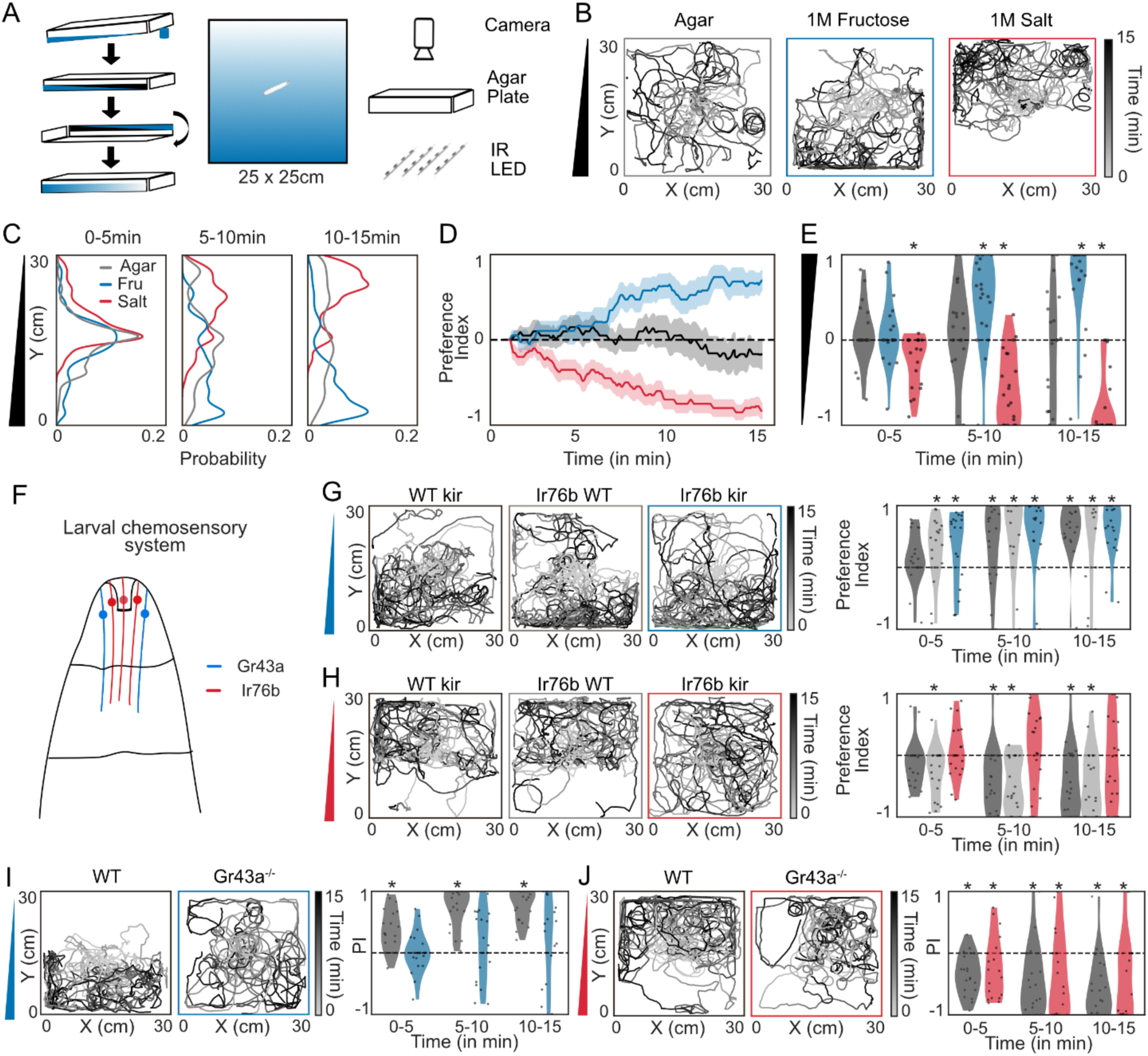
Drosophila larvae require different sensors to navigate fructose or salt taste gradients: (**A**) Schematic illustrating the preparation of a taste gradient. Individual early third-instar larvae were placed in the center; behavior was recorded for 15 min under infrared illumination. (**B**) Overlayed larval trajectories on agar (gray), fructose (blue), and salt (red) 0-1M gradients (N = 20 larvae/condition). (**C**) Larval distribution along the Y-axis for different time bins. (**D**) Preference index over time (mean: solid line; SEM: shaded). (**E**) Binned preference index (5-min intervals; violin plots with each dot representing an individual larva; bootstrapped confidence interval (CI) test against zero = Cohen’s d > 0.35; see Data S1). (**F**) Diagram of the larval sensory system. (**G**) Trajectories of WT/kir (black), Ir76b/WT (gray), and Ir76b/kir (blue) larvae on fructose gradients (N = 20). Binned preference index. (**H**) Trajectories of WT/kir (black), Ir76b/WT (gray) and Ir76b/kir (red) larvae on salt gradients (N = 20). Binned preference index. (**I**) Trajectories of WT (black) and Gr43a^-/-^ (blue) on fructose gradients (N = 20). Binned preference index. (**J**) Trajectories of WT (black) and Gr43a^-/-^ (red) on salt gradients (N = 20).

Quantification over time, using a preference index, showed that larvae progressively accumulated at high fructose and low salt concentrations (Fig. 1D, E). We further looked at larval distribution and locomotor behavior on the agar control and gradients (Fig. S1/S2). On fructose gradients, more larvae reached high fructose regions faster (= successful fructose navigation) and dwelled there longer than when they were tested only on agar (Fig. S1A). In contrast, on salt gradients, more larvae reached low-salt regions (= successful salt navigation) than on agar controls, with comparable travel and dwell times (Fig. S2A). We additionally analyzed larval movement patterns during navigation toward either the top or bottom of the arena or after arrival within the preferred region. During navigation, larvae showed overall fewer but faster and longer runs and smaller turn angles on fructose and salt compared to agar (Fig. S1B, C, S2B, C). We also detected behavioral modulations based on movement direction as larvae increased their number of runs and decreased the turn angle when moving towards high fructose concentrations (Fig. S1D, E) and when moving away from high salt concentrations (Fig. S2D, E). Upon successfully reaching preferred gradient regions, high concentrations of fructose and low concentrations of salt, larvae continued to have increased speed compared to agar controls, but had similar behavioral modulations to agar controls (Fig. 1F-I, Fig. S2F-I).

To test whether larvae rely on continuous gradient information to locate high fructose concentrations, we replaced the linear fructose gradient with a discrete 5 cm-wide strip of 1M fructose at one end of the arena, with plain agar elsewhere (Fig. S3). In the gradient condition, larvae accumulated in high fructose regions (Fig. S3A-D). In the strip conditions, however, accumulation was limited to larvae that happened to encounter the strip. Approximately 50% of larvae found the strip, but reached it faster than in the gradient condition (Fig. S3E). Larvae exposed to the strip arena had increased speed and showed no signs of slowing down during or after navigation (Fig. S3F-G).

To examine how gradient steepness and different taste concentrations affect navigation, we tested larval responses across fructose and salt gradients with maximum concentrations of either 0.5M, 1M, or 2M (Fig. S4A-L). On fructose, larvae navigated 1M gradients, showed slowed navigation on 0.5M gradients, and exhibited no directional bias on 2M gradients (Fig. S4A-F). Success rates were highest on 1M; larvae reached high fructose regions fastest and dwelled longest compared to weaker or stronger gradients. On salt, larvae displayed directional navigation in 1M and 2M gradients, but not in 0.5M gradients (Fig. S4G-L). More larvae reached low salt regions as gradient strength increased, and although travel times were similar across concentrations, dwell time was shortest in the 2M gradient. During navigation, behavioral modulation - higher run frequency, longer runs, smaller turn angles, and higher speed when moving towards high fructose and away from low salt occurred only in 1M fructose and salt gradients, respectively (Fig. S4E/F, K/L). Based on these results, 1M fructose and 1M salt gradients were used for all subsequent experiments, as these conditions produced the most robust gradient navigation.

Taste gradient navigation requires larvae to detect fructose and salt via their chemosensory system (Fig. 1F). The ionotropic receptor 76b (Ir76b) acts as a coreceptor for other ionotropic receptors detecting low concentrations of salt (Zhang et al., 2013), while the gustatory receptor 43a (GR43a) functions as the primary fructose receptor in larvae (Mishra et al., 2013). Silencing Ir76b expressing neurons using *UAS-Kir2.0* did not affect fructose gradient navigation, but abolished salt gradient navigation (Fig. 1G/H, Fig. S5). Larvae with silenced Ir76b expressing neurons had much lower success rates in reaching low salt regions (Fig. S5F), exhibited lower stop frequency and failed to modulate locomotor parameters during navigation (Fig. S5G). Wild-type (WT) larvae navigated fructose gradients effectively, accumulating in high fructose regions, whereas Gr43a^-/-^ mutants failed to do so (Fig. 1I, Fig. S6). More WT larvae reached these regions than Gr43a^-/-^ mutants, with no difference in travel time; however, Gr43a^-/-^ larvae that arrived dwelled shorter (Fig. S6B). During navigation, WT larvae had higher speed during runs and more stopping events than Gr43a^-/-^ larvae (Fig. S6C). WT larvae increased their run frequency in the direction of high fructose concentrations during navigation but showed no behavioral modulation upon arrival (Fig. S6D). Both genotypes navigated salt gradients similarly (Fig. 1J, Fig. S6E).

To assess whether olfactory sensing contributes to taste gradient navigation, we tested Orco^-/-^ mutant larvae, which have non-functioning olfactory receptors. Fructose and salt gradient navigation was similar between WT and Orco^-/-^ mutants (Fig. S7), indicating that olfaction does not contribute to larval navigation in taste gradients.

### Projection neurons are required for taste gradient navigation

Fructose and salt are detected at the sensory level by Gr43a and Ir76b, respectively. To determine how taste information reaches higher brain regions, we quantified olfactory and non-olfactory synaptic input onto various local interneurons and projection neurons using the EM connectome dataset (Winding et al., 2023). Olfactory input is based on the number of direct synapses from olfactory sensory neurons and non-olfactory input is based on the number of direct synapses from other sensory neurons entering the brain through the antennal nerve or maxillary nerve.

We focused on two octopaminergic neurons (OAN-a1, OAN-a2), which originate in the SEZ and project to the mushroom body (MB) calyx (Eichler et al., 2017; Saumweber et al., 2018; Wong et al., 2021; Franke et al., 2026), local inhibitory interneurons in the larval antennal lobe (Keystone and Picky LNs) and two classes of olfactory projection neurons - uniglomerular PNs (uPNs) and the cobra multiglomerular PN (mPN) - which connect the larval antennal lobe to higher brain regions (Berck et al., 2016; Vogt et al., 2021).

OAN-a1 and OAN-a2 receive substantially greater non-olfactory than olfactory input (Fig. 2A), indicating that they might relay gustatory and other non-olfactory information to the MB. To determine the involvement of OAN-a1/a2 neurons in taste gradient navigation, we silenced them by crossing *SS24765-Split GAL4* (Eschbach et al., 2020; Franke et al., 2026) with *UAS-Kir2.0* and tested manipulated larvae on fructose and salt gradients. Silencing OAN-a1/a2 neurons impaired fructose gradient navigation (Fig. 2B). Success rate in reaching high fructose regions was comparable to controls but larvae exhibited increased speed (Fig. S8A-C). Once in high fructose regions, larvae tend to spend less time there than controls. They increased their speed upon reaching these regions and failed to modulate their behavior across the gradient to stay in the high fructose region (Fig. S8D). In contrast, silencing OAN-a1/a2 had no effect on salt gradient navigation (Fig. 2C, Fig. S8E-H).

**Fig 2.**
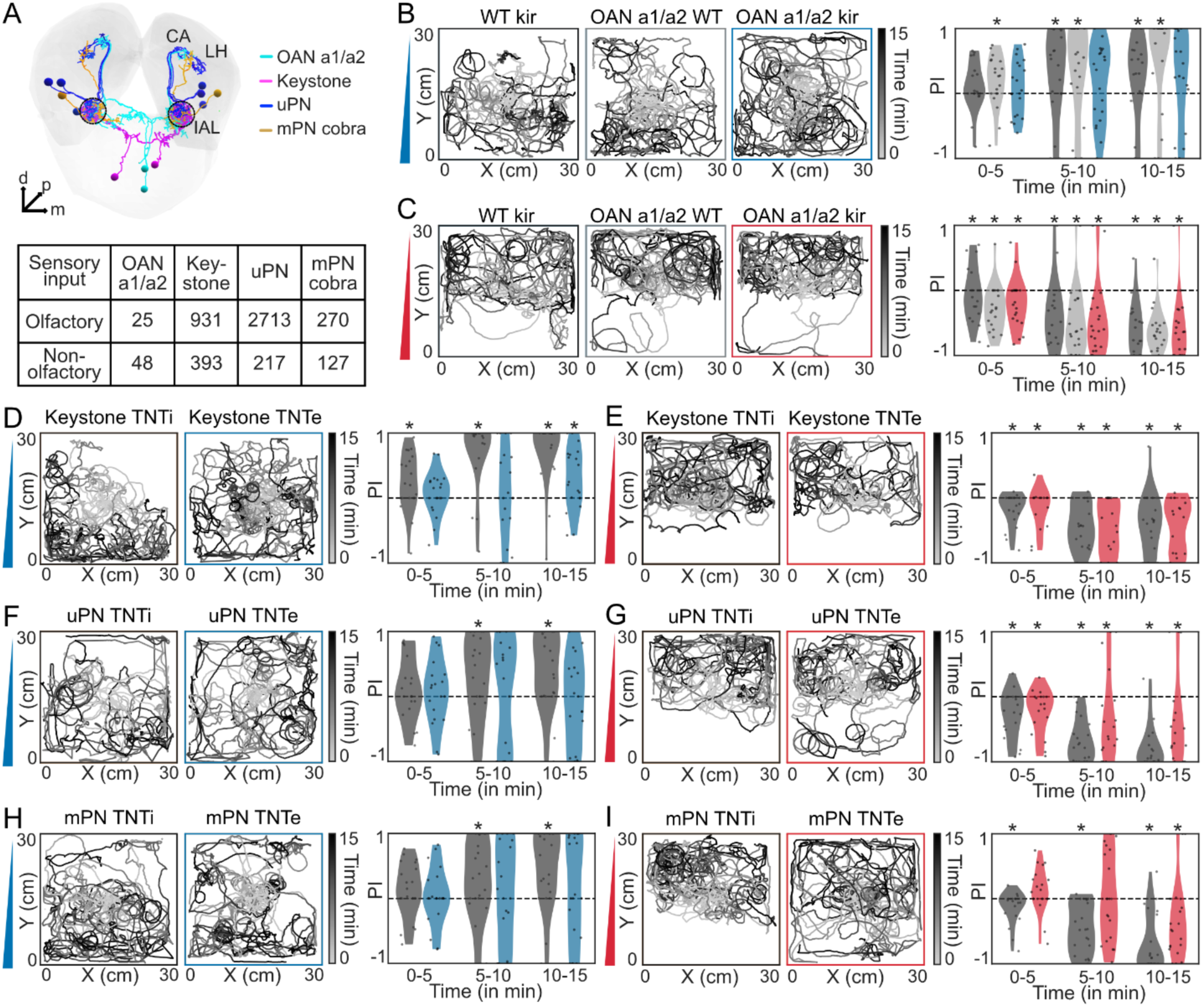
Projection neurons are required for taste gradient navigation: (**A**) Electron microscopy (EM) reconstructions of OAN a1/a2, Keystone, uPN and cobra mPN neurons (larval antennal lobe (IAL), mushroom body calyx (CA), lateral horn (LH)). Olfactory and non-olfactory input to these neurons was calculated from EM data. (**B**) Trajectories of WT/kir (black), OAN a1/a2/WT (gray) and OAN a1/a2/kir (blue) larvae on fructose gradients (N = 20). Binned preference index (5 min intervals; violin plots with each dot representing an individual larva; bootstrapped confidence interval (CI) test against zero = Cohen’s d > 0.35, see Data S1) (**C**) Trajectories of WT/kir (black), OAN a1/a2/WT (gray) and OAN a1/a2/kir (red) larvae on salt gradients (N = 20). Binned preference index. (**D**) Trajectories of Keystone/TNTi (black) and Keystone/TNTe (blue) larvae on fructose gradients (N = 20). Binned preference index. (**E**) Trajectories of Keystone/TNTi (black) and Keystone/TNTe (red) larvae on salt gradients (N = 20). Binned preference index. (**F**) Trajectories of uPN/TNTi (black) and uPN/TNTe (blue) larvae on fructose gradients (N = 20). Binned preference index. (**G**) Trajectories of uPN/TNTi (black) and uPN/TNTe (red) larvae on salt gradients (N = 20). Binned preference index. (**H**) Trajectories of mPN/TNTi (black) and mPN/TNTe (blue) larvae on fructose gradients (N = 20). Binned preference index. (**I**) Trajectories of mPN/TNTi (black) and mPN/TNTe (red) larvae on salt gradients (N = 20). Binned preference index.

Keystone, uPNs, and cobra mPNs receive substantial olfactory input but also considerable non-olfactory input, suggesting a role for these neurons in multisensory processing (Berck et al., 2016). As crosses using *UAS-Kir2.0* were not viable, we silenced these neurons by expressing tetanus toxin (*UAS-TNTe*). As a control we crossed the GAL4 lines to an inactive version of TNT (*UAS-TNTi*); both TNT lines also expressed *tsh-GAL80*, restricting GAL4 expression to the central brain.

Keystone is a prominent GABAergic local interneuron spanning the SEZ and the larval antennal lobe and connecting to many cell types in the olfactory pathway (Berck et al., 2016). Keystone-TNTe (*GMR27F08-GAL4* (Odell et al., 2022)) larvae failed to navigate fructose gradients (Fig. 2D). They had lower success rates, took longer to arrive at high fructose zones, and spent less time there than controls (Fig. S9A, B). During navigation they increased their speed and made lesser stops, which continued upon reaching high fructose (Fig. S9C, D). Keystone-silenced larvae navigated salt gradients similarly to the control (Fig. 2E, Fig. S9E-H).

We also examined picky local interneurons, which innervate the larval antennal lobe. Although dominated by olfactory input, picky neurons receive a small amount of non-olfactory input and synaptic input from OAN-a1/a2, Keystone and uPNs (Fig. S10A). Silencing picky 1, picky 3 and picky 4 LNs did not affect navigation on fructose or salt gradients (Fig. S10-12).

Silencing uPNs (*GH146-GAL4*) using TNTe disrupted fructose gradient navigation (Fig. 2F, Fig. S13A). uPN-TNTe larvae only showed a modest reduction in success rate, however, many more of them ended up in the lowest fructose region (Fig. S13B). During navigation and after reaching high fructose, they increased run frequency, run length, and speed (Fig. S13C-D). On salt gradients, uPN-TNTe larvae navigated salt but with lower success rate and reduced dwell time upon reaching low salt (Fig. 2G, Fig. S13E-F). Their navigation strategies were similar to controls, but upon reaching low salt they increased speed and reduced stop frequency towards high salt concentrations (Fig. S10G, H). Silencing the cobra mPN (*GMR32E03-GAL4*) with TNTe impaired navigation on both taste gradients (Fig. 2H, I). mPN-TNTe larvae showed much lower success rates for both fructose and salt (Fig. S14) and exhibited increased run frequency and turn angle during navigation. On fructose gradients they decreased run length towards high fructose during navigation but were similar to controls post navigation. In contrast, on salt gradients, they increased run frequency, run length and speed toward high salt post navigation.

Overall, functional requirement experiments indicate that OAN-a1/a2, Keystone, uPNs, and cobra mPN are involved in fructose gradient navigation, while uPNs and cobra mPN are involved in efficient salt gradient navigation.

### Kenyon cells and the APL neuron are required for taste gradient navigation

Intrinsic neurons of the MB are KCs, which receive multisensory input in their dendrites, the MB calyx, and monoaminergic modulation in their axons, the MB lobes and the GABAergic anterior paired lateral (APL) inhibitory neuron. The APL neuron receives extensive synaptic KC input in the MB lobes and provides inhibitory feedback to the KCs in the calyx (Mancini et al., 2023; Masuda-Nakagawa et al., 2014; Saumweber et al., 2018). EM analysis indicates that projection neurons OAN a1/a2 and uPN relay taste information to the MB calyx and to the APL neuron (Fig. 3A).

**Fig 3.**
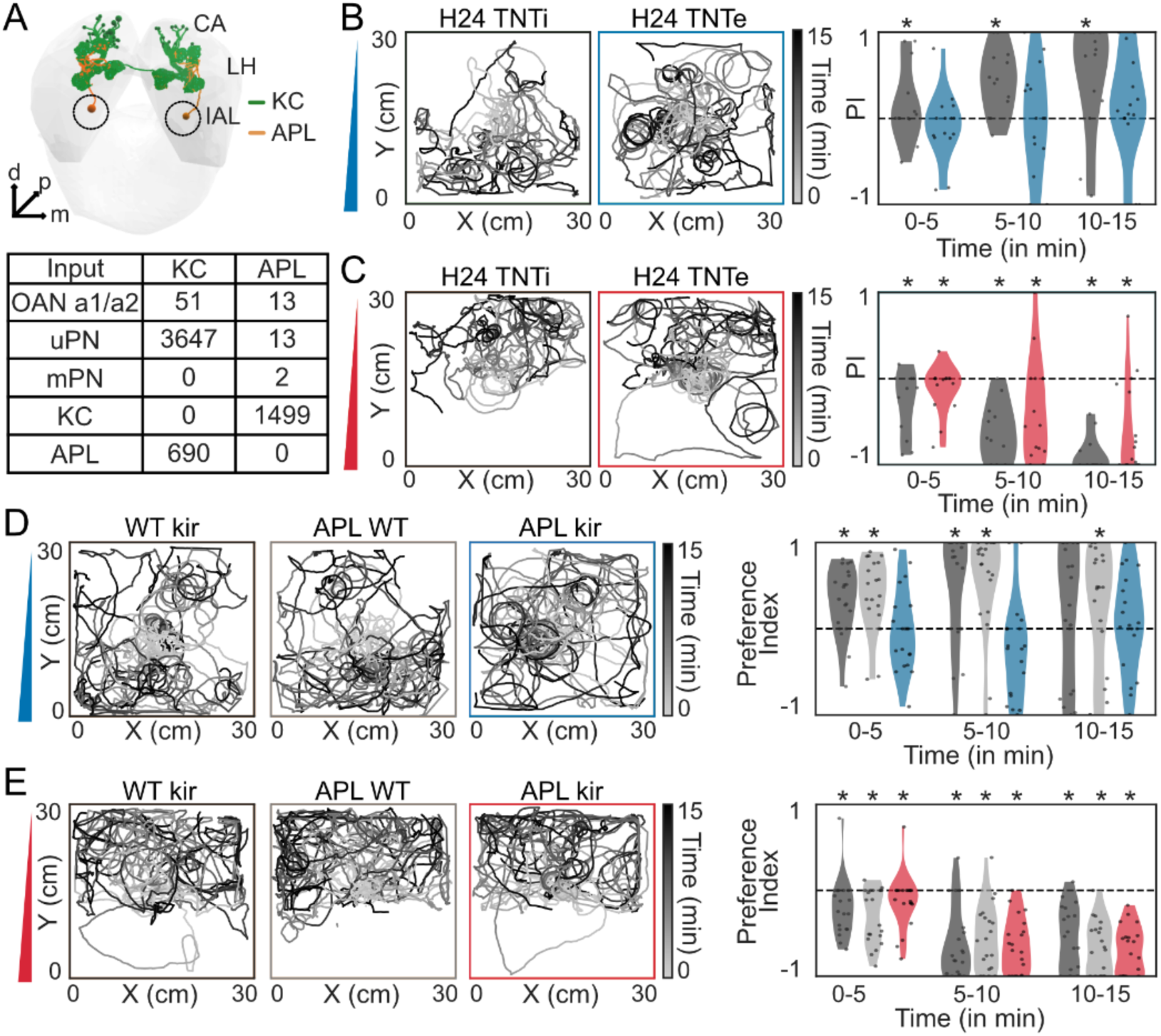
Kenyon Cells and APL neurons are required for taste gradient navigation: (**A**) Electron microscopy (EM) reconstructions of mushroom body (green) and APL (orange) neurons (larval antennal lobe (IAL), mushroom body calyx (CA), lateral horn (LH)). The number of synapses between cell types was calculated from EM data. (**B**) Trajectories of H24/TNTi (black) and H24/TNTe (blue) larvae on fructose gradients (N = 20). Binned preference index (5 min intervals; violin plots with each dot representing an individual larva; bootstrapped confidence interval (CI) test against zero = Cohen’s d > 0.35, see Data S1) (**C**) Trajectories of H24/TNTi (black) and H24/TNTe (red) larvae on salt gradients (N = 20). Binned preference index. (**D**) Trajectories of WT/kir (black), APL/WT (gray) and APL/kir (blue) larvae on fructose gradients (N = 20). Binned preference index. (**E**) Trajectories of WT/kir (black), APL/WT (gray) and APL/kir (red) larvae on salt gradients (N = 20). Binned preference index.

To test the MB’s role in taste gradient navigation, we first silenced KCs (*H24-GAL4*) with TNTe. These larvae failed to navigate fructose gradients (Fig. 3B, Fig. S15A) and showed a reduced success rate for reaching high fructose concentrations (Fig. S15B). During navigation they exhibited longer runs and smaller turn angles, and they maintained an elevated speed after reaching high fructose regions compared to controls (Fig. S15C, D). On salt gradients, larvae with silenced KCs were still capable of navigation but did so more slowly (Fig. 3C, Fig. S15E). They ended up more often in the high salt region and showed a slightly reduced success rate for reaching low salt concentrations, took longer to arrive, and spent less time in low salt concentrations (Fig. S15F). As with fructose, their navigation was characterized by increased run length both during navigation and after reaching low salt regions (Fig. S15G, H).

Silencing KCs with a different Gal4 line (*MB247-GAL4*) produced similar effects (Fig. S16). These larvae could not navigate fructose gradients, displayed increased run frequency during navigation, and showed prolonged run length after reaching high fructose regions (Fig. S16A-F). They retained the ability to navigate salt gradients but had a lower success rate for reaching low salt regions (Fig. S16G-J) and had an increased speed and run length during navigation and increased run frequency upon reaching low salt regions (Fig. S16K/L).

To probe APL’s (*SS01671-Split Gal4*) influence, we silenced the neuron using *UAS-Kir2.0*. APL silenced larvae were impaired in navigating fructose gradients (Fig. 3D, Fig. S17A), exhibiting a reduced success rate for reaching high fructose concentrations and longer times to arrive there (Fig. S17B). Similar to larvae with silenced KCs, they showed increased run length and speed during navigation (Fig. S17C). They had larger turn angles as they navigated toward high fructose concentrations, and even after approach (Fig. S17D). In contrast, APL-silenced larvae could navigate salt gradients, with success rates and behavioral strategies during/post navigation comparable to controls (Fig. 3E, Fig. S17E-H). Our data indicates that disrupting MB circuitry by silencing KCs impairs navigation of both fructose and salt gradients, whereas silencing the APL neuron impairs only fructose gradient navigation.

### Dopamine neurons are required for taste gradient navigation

KCs receive modulatory input from eight dopaminergic neurons (DANs) that cluster into two anatomically distinct clusters in fly larvae: the primary protocerebral anterior medial (pPAM) and the dorsolateral 1 (DL1) (Eichler et al. 2017, Fig 4A). Each DAN innervates a specific, non-overlapping MB compartment for valence coding (Eichler et al., 2017; Rohwedder et al., 2016; Saumweber et al., 2018; Schleyer et al., 2020). pPAM-associated DANs encode reward, while specific DL1 DANs encode punishment, allowing associative learning in the MB with positive and negative reinforcement (Eschbach et al., 2020; Weber et al., 2025).

**Fig 4.**
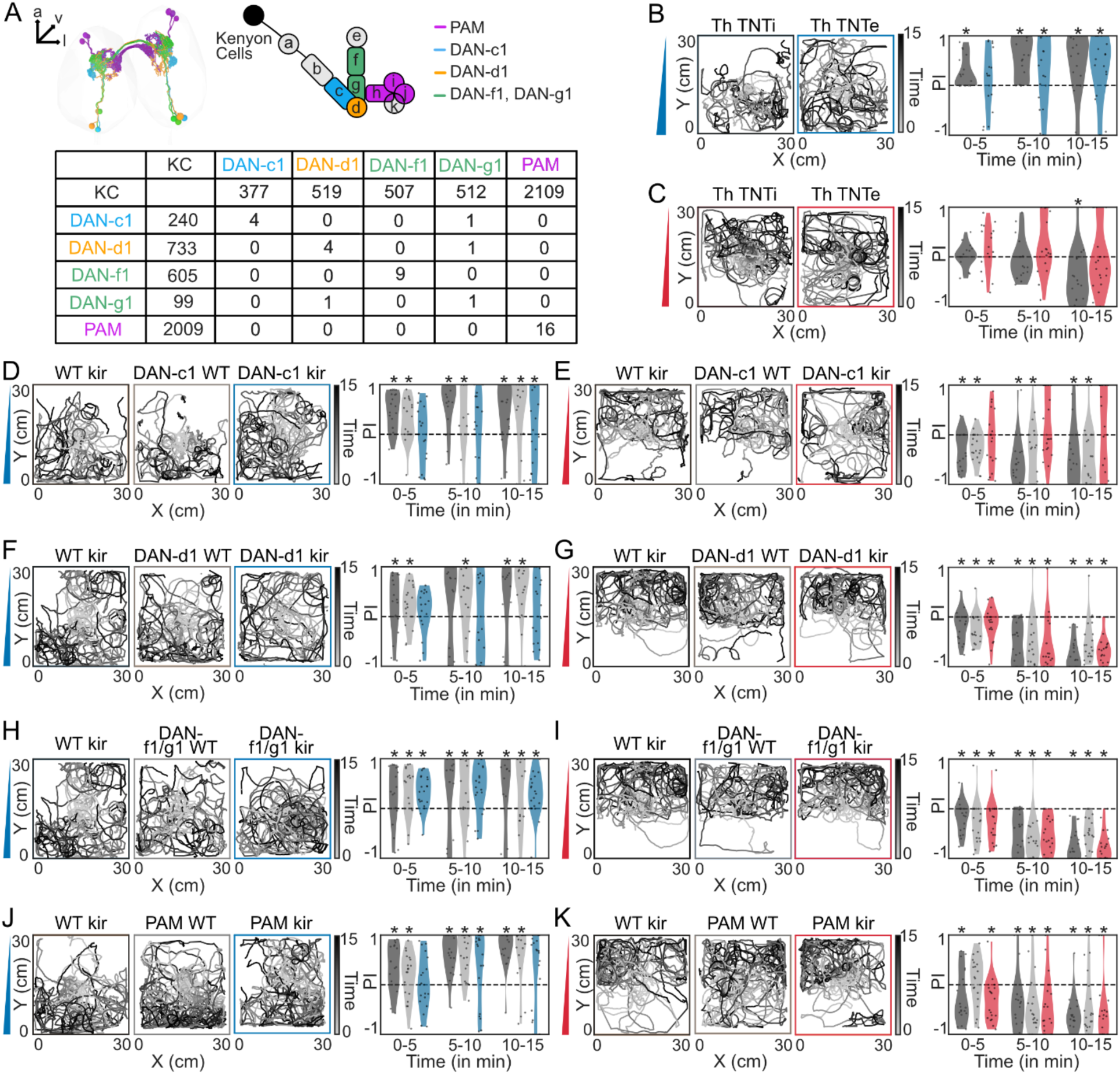
Specific dopamine neurons are required for fructose and salt gradient navigation: (**A**) Electron microscopy (EM) reconstructions of DAN-c1 (blue), DAN-d1 (yellow), DAN-f1, DAN-g1 (green) and PAM (purple) neurons. The number of synapses between cell types was calculated from EM data. (**B**) Trajectories of Th/TNTi (black) and Th/TNTe (blue) larvae on fructose gradients (N = 20). Binned preference index (5 min intervals; violin plots with each dot representing an individual larva; bootstrapped confidence interval (CI) test against zero = Cohen’s d > 0.35, see Data S1) (**C**) Trajectories of Th/TNTi (black) and Th/TNTe (red) larvae on salt gradients (N = 20). Binned preference index. (**D**) Trajectories of WT/kir (black), DAN-c1/WT (gray) and DAN-c1/kir (blue) larvae on fructose gradients (N = 20). Binned preference index. (**E**) Trajectories of WT/kir (black), DAN-c1/WT (gray) and DAN-c1/kir (red) larvae on salt gradients (N = 20). Binned preference index. (**F**) Trajectories of WT/kir (black), DAN-d1/WT (gray) and DAN-d1/kir (blue) larvae on fructose gradients (N = 20). Binned preference index. (**G**) Trajectories of WT/kir (black), DAN-d1/WT (gray) and DAN-d1/kir (red) larvae on salt gradients (N = 20). Binned preference index. (**H**) Trajectories of WT/kir (black), DAN-f1/g1/WT (gray) and DAN-f1/g1/kir (blue) larvae on fructose gradients (N = 20). Binned preference index. (**I**) Trajectories of WT kir (black), DAN-f1/g1/WT (gray) and DAN-f1/g1/kir (red) larvae on salt gradients (N = 20). Binned preference index. (**J**) Trajectories of WT/kir (black), PAM/WT (gray) and PAM/kir (blue) larvae on fructose gradients (N = 20). Binned preference index. (**K**) Trajectories of WT/kir (black), PAM/WT (gray) and PAM/kir (red) larvae on salt gradients (N = 20). Binned preference index.

The *TH-Gal4* driver labels the four DANs of the DL1 cluster but does not label the pPAM DANs (Weber et al., 2025). To assess the role of aversive DANs, we crossed *TH-Gal4* with TNTe. These larvae had overall increased speed, had a delay at navigating fructose gradients (Fig. 4B, Fig. S18) and failed to navigate salt gradients (Fig. 4C, Fig. S18). To identify the role of specific MB subcompartments, we silenced individual DL1 DANs using specific Gal4 driver lines crossed with *UAS-Kir2.0*.

Silencing DAN-c1 (*SS02160-Split Gal4*) delayed fructose gradient navigation (Fig. 4D, Fig. S19A). Larvae took longer to reach high fructose regions and spent less time there (Fig. S19B). During navigation they exhibited increased run length and speed and reduced turn angle, which they maintained after reaching high fructose regions (Fig. S19C, D). DAN-c1 silenced larvae also failed to navigate salt gradients (Fig. 4E, Fig. S19E) and had a much lower success rate at finding low salt regions (Fig. S19F). Like on fructose, they had increased run length and reduced turn angle during navigation on salt gradients (Fig. S19G). However, in low salt regions, their behavior matched controls (Fig. S19H).

Silencing DAN-d1 (*MB328B-Gal4*) abolished only fructose gradient navigation (Fig. 4F, Fig. S20A). These larvae had a lower success rate at reaching high fructose regions and tended to spend less time there (Fig. S20B). They also failed to modulate their behavior when moving toward high fructose, showing a decreased run frequency in that direction (Fig. S20C). After reaching high fructose, they displayed increased run length and speed (Fig. S20D). DAN-d1 silenced larvae could navigate salt gradients (Fig. 4G, Fig. S20). Silencing DAN-f1/g1 (*MB054B-Gal4*) had no effect on fructose or salt gradient navigation (Fig. 4H, I, S21A-H).

Silencing the pPAM cluster (*GMR58E02-Gal4*) impaired only fructose gradient navigation (Fig. 4J, Fig. S22A). pPAM silenced larvae showed a lower success rate for reaching high fructose regions (Fig. S22B). They had an increased turn angle during navigation, which persisted after reaching the high fructose regions (Fig. S22C, D) Silencing the pPAM cluster had no effect on salt gradient navigation, larvae even had lower travel time and higher dwell time compared to controls (Fig. 4K, Fig. S22E-H).

### Mushroom body output neurons are required for taste gradient navigation

KCs and DANs converge onto mushroom body output neurons (MBONs) (Eichler et al., 2017; Eschbach et al., 2020; Thum & Gerber, 2019). Specifically, DAN-c1 and DAN-d1 synapse onto MBON-c1 and MBON-d1 respectively (Fig. 5A), while pPAM DANs synapse onto MBON-h1/h2 and MBON-i1. MBONs also form inter-compartmental connections. MBON-c1 and MBON-d1 are reciprocally connected. MBON-h1/h2 provides input to MBON-c1 and MBON-d1, and together with MBON-i1 they synapse onto MBON-m1 (Eschbach et al., 2021, 2025). In addition, MBON-m1 also receives direct input from the cobra mPN, bypassing the MB circuit.

**Fig 5.**
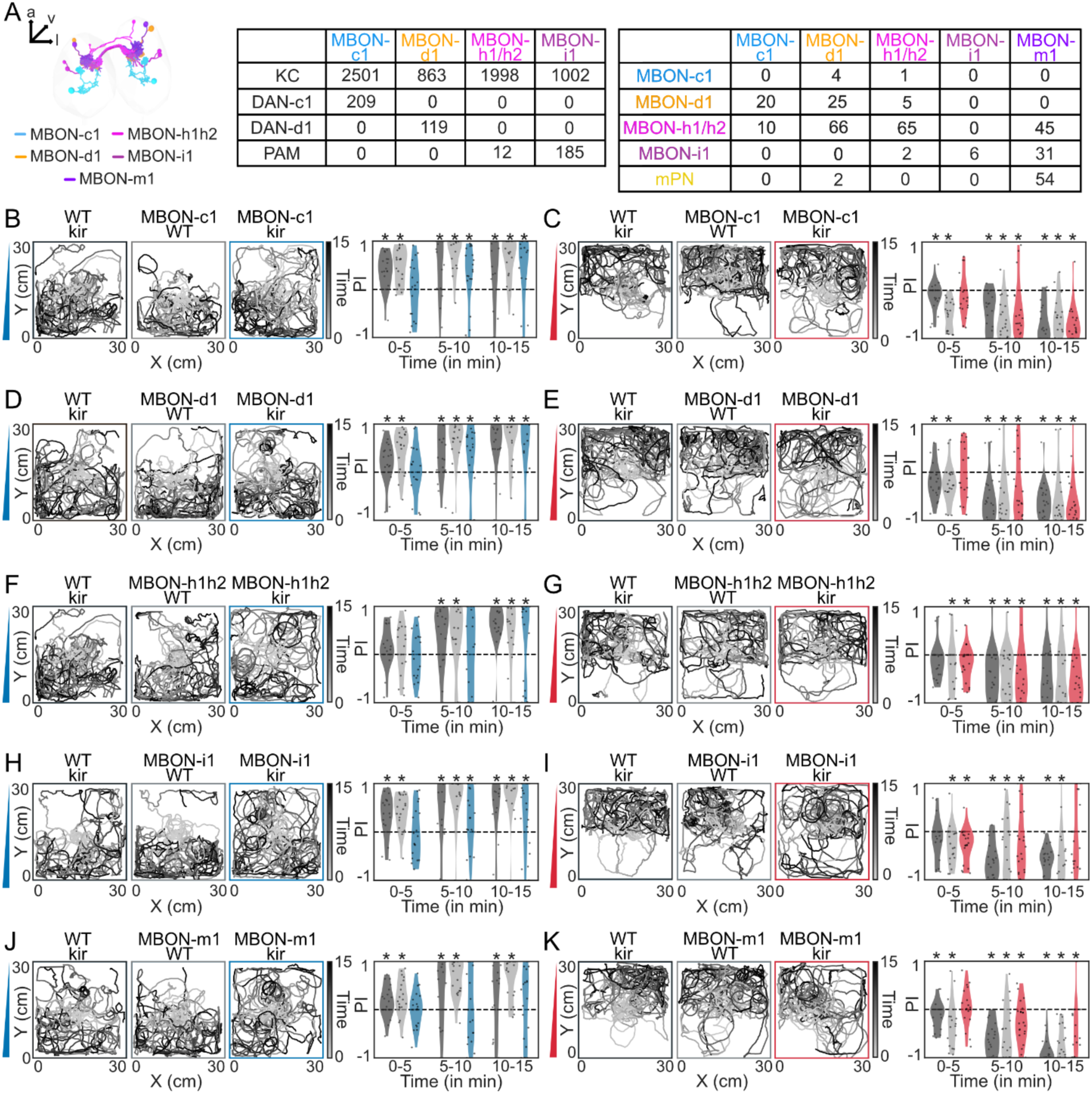
Mushroom body output neurons are required for fructose and salt gradient navigation: (**A**) Electron microscopy (EM) reconstructions of MBON-c1 (blue), MBON-d1 (yellow), MBON-h1/h2 (pink), MBON-i1 (purple) and MBON-m1 (violet) neurons. The number of synapses between cell types was calculated from EM data. (**B**) Trajectories of WT/kir (black), MBON-c1/WT (gray) and MBON-c1/kir (blue) larvae on fructose gradients (N = 20). Binned preference index (5 min intervals; violin plots with each dot representing an individual larva; bootstrapped confidence interval (CI) test against zero = Cohen’s d > 0.35, see Data S1) (**C**) Trajectories of WT/kir (black), MBON-c1/WT (gray) and MBON-c1/kir (red) larvae on salt gradients (N = 20). Binned preference index. (**D**) Trajectories of WT kir/(black), MBON-d1/WT (gray) and MBON-d1/kir (blue) larvae on fructose gradients (N = 20). Binned preference index. (**E**) Trajectories of WT/kir (black), MBON-d1/WT (gray) and MBON-d1/kir (red) larvae on salt gradients (N = 20). Binned preference index. (**F**) Trajectories of WT/kir (black), MBON-h1/h2/WT (gray) and MBON-h1/h2/kir (blue) larvae on fructose gradients (N = 20). Binned preference index. (**G**) Trajectories of WT/kir (black), MBON-h1/h2/WT (gray) and MBON-h1/h2/kir (red) larvae on salt gradients (N = 20). Binned preference index. (**H**) Trajectories of WT/kir (black), MBON-i1/WT (gray) and MBON-i1/kir (blue) larvae on fructose gradients (N = 20). Binned preference index. (**I**) Trajectories of WT/kir (black), MBON-i1/WT (gray) and MBON-i1/kir (red) larvae on salt gradients (N = 20). Binned preference index. (**J**) Trajectories of WT/kir (black), MBON-m1/WT (gray) and MBON-m1/kir (blue) larvae on fructose gradients (N = 20). Binned preference index. (**K**) Trajectories of WT/kir (black), MBON-m1/WT (gray) and MBON-m1/kir (red) larvae on salt gradients (N = 20). Binned preference index.

We tested the requirement of MBONs for taste gradient navigation. Silencing MBON-c1 (*SS01776-Split Gal4*) delayed fructose and salt gradient navigation (Fig. 5B/C, Fig. S23). Larvae showed reduced success rates, longer travel times, and weaker modulation of behavioral parameters while navigating fructose gradients (Fig. S23B, C). After reaching high fructose regions, they exhibited increased run lengths (Fig. S23D). They also had reduced success reaching low salt, potentially due to increased reorientation frequency (Fig. S23F, G).

Silencing MBON-d1 (*SS01705-Split Gal4*) delayed fructose navigation (Fig. 5D, Fig. S24A). Larvae showed reduced success at reaching high fructose, tended to take longer to arrive and stayed there for shorter periods (Fig. S24B). During navigation they increased run length and decreased turn angle, which persisted after reaching high fructose (Fig. S24C, D). Silencing MBON-d1 delayed salt navigation (Fig. 5E, Fig. S24E, F), showing decreased run frequency and reduced turn angles during navigation (Fig. S24G). However, larval behavior after reaching low salt was comparable to controls (Fig. S24H).

Silencing MBON-h1/h2 (*SS00894-Split Gal4*) delayed fructose navigation (Fig. 5F, Fig. S25A). Larvae had reduced success reaching high fructose concentrations (Fig. S25B), increased run length and decreased turn angle during navigation, and failed to modulate speed and turn angle appropriately when moving toward higher fructose (Fig. S25C). They continued to show increased run length after reaching high fructose regions (Fig. S25D). MBON-h1/h2 silencing did not affect salt gradient navigation (Fig. 5G, S25E-H).

Silencing MBON-i1 (*SS01726-Split Gal4*) delayed fructose navigation (Fig. 5H, Fig. S26A) and reduced success at reaching high fructose concentrations (Fig. S26B). Larvae showed reduced behavioral modulation during navigation and maintained increased speed and reduced stop frequency after reaching high fructose regions (Fig. S26C, D). MBON-i1 silenced larvae could navigate salt gradients but were not retained in low salt regions, with a tendency toward reduced dwell time (Fig. 5I, Fig. S26E, F). After reaching low salt regions, they had an increased run length and speed and decreased turn angle compared to the controls (Fig. S26G, H).

Silencing MBON-m1 (*SS02163-Split Gal4*) abolished fructose gradient navigation (Fig. 5J, Fig. S27A). Larvae had reduced success in reaching high fructose regions and shorter dwell times (Fig. S27B), with increased run frequency and run length and decreased turn angle during navigation (Fig. S27C). They had an increase in run frequency and turn angle towards low fructose after reaching high fructose regions (Fig. S27D). MBON-m1 silencing reduced and delayed salt gradient navigation (Fig. 5K, Fig. S27E), reducing success at reaching low salt (Fig. S27F), and increasing run length and turn angle during navigation (Fig. S27G). However, after reaching low salt regions, their behavior was similar to the controls (Fig. S27H).

Together, our data indicates that silencing MBON-c1, MBON-d1, MBON-h1/h2 and MBON-i1 delays fructose gradient navigation, while MBON-c1, MBON-d1 and MBON-i1 are required for salt gradient navigation. These MBONs converge onto MBON-m1 and silencing of MBON-m1 disrupts navigation in both fructose and salt gradients.

### Neural circuit underlying taste gradient navigation

In summary, we find that fructose and salt gradient navigation in *Drosophila* larvae requires distinct but partially overlapping neural pathways (Fig. 6A). Fructose detection is mediated by Gr43a sensors, whereas salt detection requires Ir76b expressing sensory neurons. Fructose sensory information seems to get integrated into early processing centers, such as via the keystone local interneuron in the larval antennal lobe and relayed to the MB calyx via the octopaminergic OAN-a1/a2 projection neuron. Two other types of projection neurons, uPNs and the cobra mPN are required for both fructose and salt gradient navigation.

**Fig 6.**
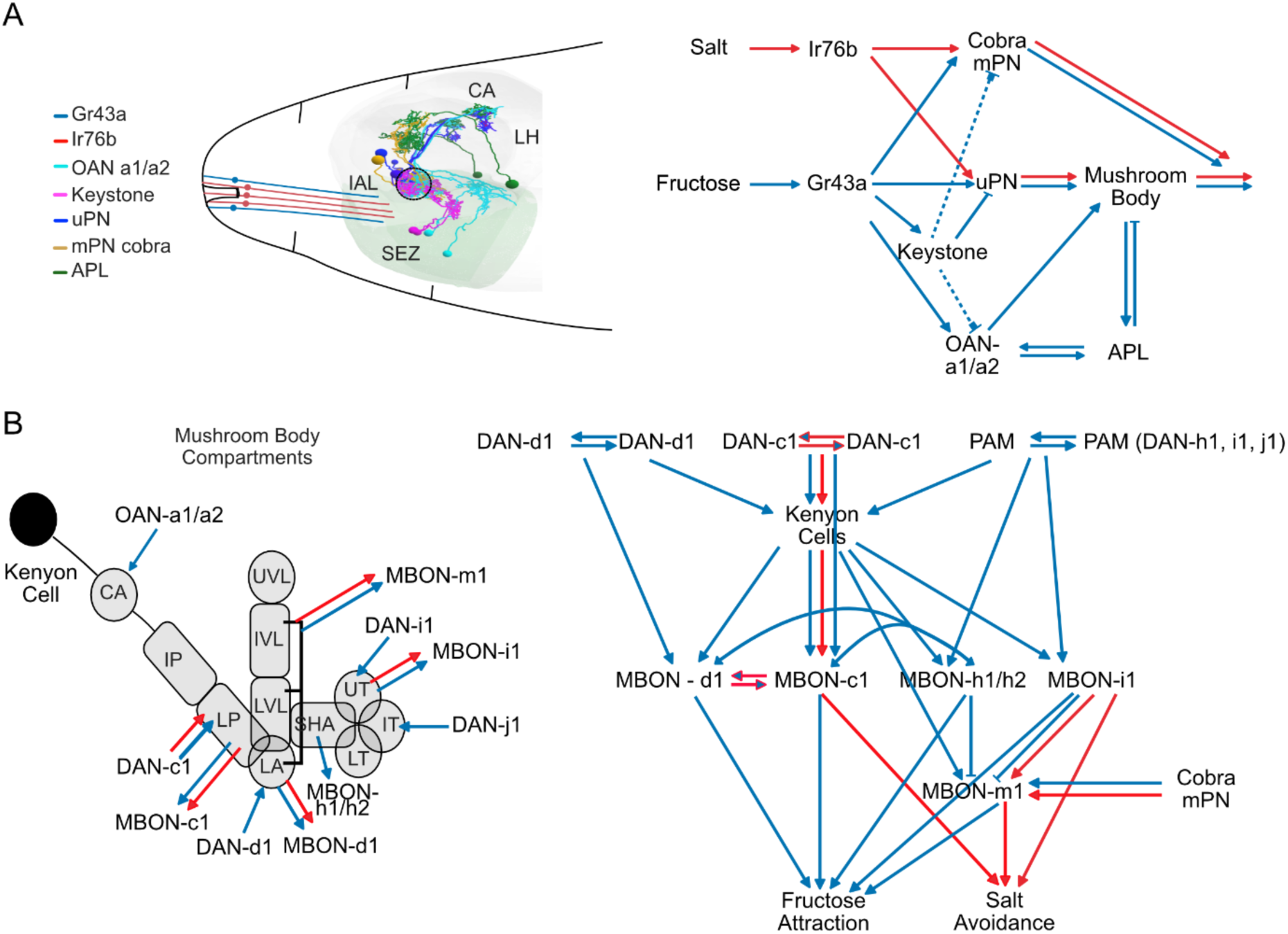
Neural circuit underlying taste gradient navigation: (**A**) Electron microscopy (EM) reconstructions of sensory neurons, interneurons and projection neurons (larval antennal lobe (IAL), subesophageal zone (SEZ), mushroom body calyx (CA), lateral horn (LH)). Fructose and salt are detected by Gr43a and Ir76b respectively. The sensory signals are transmitted via projection neurons to the mushroom body calyx. (**B**) Schematic of the larval mushroom body compartments: CA - calyx, IP - intermediate peduncle, LP - lower peduncle, UVL - upper vertical lobe, IVL - intermediate vertical lobe, LVL - lower vertical lobe, LA - lateral appendix, SHA - shaft, UT - upper toe, IT - intermediate toe, LT - lower toe. Distinct dopaminergic neurons and mushroom body output neurons drive attraction to high fructose and avoidance of high salt. Blue arrows in the circuit denote pathways required for fructose gradient navigation and red arrows denote pathways required for salt gradient navigation. Dotted lines indicate non-direct synaptic connections.

On the MB input level, the APL neuron and different dopaminergic neurons, such as DAN-c1, DAN-d1, also labeled by TH-GAL4, and pPAM cluster neurons are required for fructose navigation, while only DAN-c1 (TH-GAL4) is additionally required for salt gradient navigation (Fig. 6B). Several MB output neurons are necessary for proper navigation of both fructose and salt gradients. Many of these MBONs converge onto MBON-m1, which we find essential for both fructose attraction and salt avoidance.

## Discussion

Animals must continuously evaluate environmental cues during foraging to guide approach or avoidance. How gustatory information is processed to enable navigation of taste gradients is not well understood. We demonstrate that *Drosophila* larvae are capable of navigating both attractive and aversive taste gradients. Having established a robust behavior assay allowed us to systematically map the basic neural circuit requirements underlying taste navigation, beginning from peripheral chemosensory neurons that detect taste cues, extending through parallel projection pathways that convey taste information to higher-order brain regions, such as the MB (Luo et al., 2014; Vogt, 2020). We find that navigating both fructose and salt gradients requires a partially shared MB circuit, suggesting a common valence-independent neural architecture for taste gradient information processing. Generally, navigating an attractive gradient requires a broader population of neurons compared to aversive conditions. We tested the requirement of cell types from the sensory periphery to the MBON-m1, where output from the MB converges and which shapes activity of descending neurons and the behavioral output toward approach or avoidance. Our findings provide a circuit-level account of how information of taste gradients of different valences is detected, processed and transformed into navigational decisions.

### *Drosophila* larvae can navigate taste gradients

Larvae navigating fructose gradients show a positive preference index, high success rates, and increased dwell time in high-fructose regions compared to pure agar navigation (Fig. 1, Fig. S1). During navigation we find directional modulation of locomotor parameters, a navigational strategy called klinotaxis (Gomez-Marin & Louis, 2012), characterized by increased run frequency and reduced turn angles when oriented toward higher fructose concentrations. These responses are consistent with navigation strategies described for olfactory (Gomez-Marin et al., 2011), thermal (Luo et al., 2010), and phototactic (Sawin et al., 1994) gradients in *Drosophila* larvae. Successful navigation to the favorable regions requires spatial gradient information for the larvae to rely on, as success is substantially reduced when high fructose is presented in a strip without a gradient (Fig. S3).

In contrast, larvae exhibit avoidance behavior in salt gradients, as indicated by a negative preference index, accumulation preferentially in low-salt regions and high success rates in reaching these zones (Fig. 1, Fig. S2). Dwell time on low salt is not different from agar, likely due to the minimal physiological requirement for salt (King, 1953; Russell et al., 2011). During navigation toward low salt regions, larvae use similar locomotor modulations as in fructose gradients, including increased run frequency and run length, along with reduced turn angles when oriented toward lower salt concentrations.

Across both gradient types, larvae increase their speed upon reaching favorable conditions, in contrast to previous studies reporting reduced speed as a mechanism to promote retention within such environments (Wosniack et al., 2022; Mudunuri et al., 2026). This behavior is, however, consistent with a rover-like phenotype, where locomotion increases in resource-rich environments to enhance sampling over a broader spatial scale (Padilla Perez et al., 2025). In the context of gradient navigation, increased speed within the favorable environment may function as an active sampling strategy, allowing larvae to more thoroughly characterize the spatial structure of the gradient. Furthermore, our gradient environments lack protein, which is essential for larval growth and development (Wu et al., 2014), therefore, an increase in speed could also reflect an ongoing search for additional resources with protein.

A gradient of 0-1M fructose was selected for neural circuit dissection experiments. 0-0.5M gradients were too shallow to guide effective navigation, whereas 0-2M gradients seem to render much of the arena uniformly favorable, reducing the need for directed movement (Fig. S4). Similarly, 0-1M salt gradients were used, as 0-2M gradients make the high salt regions easily avoidable, whereas 0-0.5M gradients create broadly favorable conditions that diminish the effectiveness of gradient-guided behavior.

### Requirement of early processing centers for taste gradient navigation

Taste gradient navigation in *Drosophila* larvae should depend on the detection of appetitive and aversive gustatory cues. Our results confirm that the receptors Gr43a and Ir76b are required for fructose and salt sensing (Fig. 1F-J, Fig. S5, 6), respectively, consistent with previous studies (Mishra et al., 2013; Zhang et al., 2013). The absence of navigation deficits in Orco^-/-^ mutants further demonstrates that fructose and salt gradients are sensed via gustatory and not olfactory inputs (Fig. S7).

We find that downstream of sensory detection, gustatory information is relayed to higher brain regions through projection neurons with distinct and partially overlapping functional roles. OAN-a1/a2 neurons, which receive predominantly non-olfactory input from the SEZ, the early gustatory processing region in fly larvae, and project to the MB calyx (Fig. 2A), are selectively required for fructose but not salt gradient behavior (Fig. 2B-C, Fig. S8). Although larvae with impaired neurons reach high fructose regions, they fail to remain there, indicating that OAN-a1/a2 are not required for gradient navigation but for sustained local exploitation. This aligns with their role in context-dependent reinforcement, where they modulate the mushroom body circuit during both attractive and aversive classical learning (Schroll et al., 2006; Selcho et al., 2014; Saumweber et al., 2018; Franke et al., 2026).

Also neurons in the early olfactory processing center, the larval antennal lobe, receive substantial non-olfactory sensory input (Fig. 2A, Berck et al. 2016). We found that the local inhibitory keystone neuron is required for fructose gradient navigation, but not the picky LNs. These findings suggest that even early processing centers, directly downstream of sensory neurons, can have multimodal functions (Odell et al., 2022).

Larval antennal lobe output neurons, the uPNs, are required for fructose and salt navigation, with larvae failing to remain in low salt regions upon arrival (Fig. 2D-E, Fig. S10). KCs, which integrate major input from uPNs among other projection neurons (Fig. 3A), are required for navigation on both fructose and salt gradients (Fig. 3B-C, Fig. S15, S16), indicating a general role for MB dependent processing across gustatory modalities. KC silencing does not have an effect on fructose or salt preference in a simple two choice preference assay (Pauls et al., 2010; Widmann et al., 2016). Consistent with this, KC silencing results in prolonged run lengths during both navigation and post-arrival phases, suggesting a defect in locomotor decision-making, such as stopping and reorienting, as in adult flies (Martin et al., 1998; Sakai & Kitamoto, 2006), rather than sensory detection. Effective gradient navigation requires comparison of current sensory input with recent experience, implying a role for MB mediated working memory, in addition to its role in classical short-term associative learning (Rohwedder et al. 2016; Saumweber et al. 2018). The observed decreased stopping phenotype may therefore reflect an inability to update the sensory input that normally signals when to terminate a run (Tastekin et al., 2018).

The APL neuron, which receives input from OAN-a1/a2 and KC axons and provides GABAergic feedback to the MB calyx, is selectively required for fructose but not salt gradient navigation (Fig. 3D-E, Fig. S17), consistent with its role in appetitive short-term memory (Mancini et al., 2023). APL silencing does not impair fructose preference in a two-choice assay, further supporting its role in navigational computation rather than sensory detection (Saumweber et al., 2018).

Finally, the multiglomerular cobra mPN is required for navigation in both fructose and salt gradients (Fig. 2H, I, Fig. S11). Additionally, cobra mPN has been shown to be required for aversive olfactory gradient navigation (Vogt et al., 2021), consistent with its proposed function as a multisensory integrator of gustatory, olfactory and thermal information based on EM connectivity (Eschbach et al., 2025). Silencing of cobra mPN reduces success rates and disrupts locomotor modulation during the taste gradient navigation phase. Cobra mPN provides direct input to MBON-m1, bypassing intrinsic MB circuitry, which may enable parallel, state- or context-specific processing (Vogt et al., 2021).

We found requirements of three distinct projection neuron types for gustatory navigation, which however differ in their amount of multisensory input and seem to serve different functional roles (Miroschnikow et al., 2018).

### Valence-specific dopaminergic signals converge onto mushroom body output neurons to guide gradient navigation

Taste gradient navigation requires larvae to compare stimulus concentrations as they move through the gradient, implying that the underlying circuit must encode not only stimulus presence but its relative value across space. In adult *Drosophila*, dopaminergic neurons encode the relative values of attractive and aversive stimuli (Villar et al., 2022; Martinez-Cordera et al., 2025). Comparable dopaminergic circuits are present in larvae (Rohwedder et al., 2016; Weber et al., 2025).

Silencing DAN-c1 (and *TH-GAL4*, which also labels DAN-c1) impaired both fructose and salt navigation (Fig. 4D-E, Fig. S19), consistent with results from functional imaging showing that these neurons respond to both stimuli with excitation (Weber et al., 2025). This involvement across modalities suggests that DAN-c1 neurons may contribute to relative value coding independently of stimulus valence. Notably, DAN-c1 has previously been shown to not be needed for most short-term appetitive or aversive associative memory assays (Saumweber et al., 2018; Eschbach et al., 2020; Weber et al., 2025), indicating that sensory detection and reinforcement processing are intact, and instead points to a function in short-term value comparison during navigation. Consistent with dopaminergic value processing, dopamine receptors in DAN-c1 have been implicated in larval aversive conditioning, suggesting that receptor-mediated modulation may shape how dopaminergic neurons contribute to learned value representations (Qi et al., 2025). In adults, this idea is further supported by evidence that dopamine receptors in pPAM neurons differentially modulate reward learning depending on stimulus intensity (Saito et al., 2026).

Silencing pPAM neurons impaired fructose navigation but had no effect on salt navigation (Fig. 4J-K, Fig. S22), different from earlier findings in the fructose two-choice assay (Rohwedder et al., 2016). The selective requirement for fructose gradients is, however, consistent with the established role of pPAM neurons as a reward-encoding population responsive to fructose (Eschbach et al., 2020; Saumweber et al., 2018; Weber et al., 2025).

Silencing DAN-d1 also selectively abolished fructose navigation while leaving salt navigation intact (Fig. 4F-G, Fig. S20), despite the absence of a detectable calcium response to fructose (Weber et al., 2025). This dissociation suggests that DAN-d1 does not directly encode fructose but might instead contribute indirectly, likely through downstream circuit interactions or indirect modulation. Activation of FAN-7, a postsynaptic partner of DAN-d1, induces appetitive memory, raising the possibility that DAN-d1 shapes reward processing via its output targets rather than by encoding stimulus value itself (Eschbach et al., 2020).

Finally, silencing DAN-f1/g1 had no effect on fructose or salt gradient navigation (Fig. 4H-I, Fig. S21), despite their established role in aversive associative memory (Eschbach et al., 2021; Weber et al., 2025). Our findings indicate that gradient navigation relies on dopaminergic mechanisms that are partially shared but also distinct from those underlying classical associative learning, depending on short-term comparisons of relative stimulus value.

The compartment-specific DANs and KCs converge onto mushroom body output neurons (MBONs). Silencing MBON-c1, MBON-d1, MBON-h1/h2, and MBON-i1 individually delays fructose navigation (Fig. 5). These results indicate that functional coding and cross-talk among MBONs seems to be required (Fig. 5A, Aso et al. 2014). MBON-c1 and MBON-d1 are additionally required for salt navigation, consistent with their multimodal input from DAN-c1. Beyond their roles in chemosensory evaluation, MBON-c1 and MBON-d1 may contribute to navigation by being part of the larval central complex (Lungu et al., 2025). The c1 compartment of the larval MB is associated with the putative fan-shaped body cell-type and MBON-d1 signals via MB2ON-63, a putative protocerebral bridge horizontal fiber cell-type, potentially integrating spatial or motor-related information (Lungu et al., 2025).

In contrast, larvae with silenced MBON-i1 successfully navigate toward low-salt regions but fail to remain there, pointing to a function in goal retention rather than navigation itself (Fig. 5H-I, Fig. S26). This is consistent with its position downstream of pPAM, suggesting that reward related signals may be required to evaluate low salt as a positive reward signal.

All four MBONs converge onto MBON-m1, which also receives direct sensory input from the cobra mPN, bypassing the MB circuitry. Silencing MBON-m1 abolishes fructose navigation and impairs salt navigation (Fig. 5J-K, Fig. S27), confirming its role as a key integrator that encodes modality-independent stimulus valence and translates distributed MBON signals into behavioral decisions (Eschbach et al., 2025).

The neural circuit underlying taste navigation overlaps with the neural circuit underlying associative learning, however, the computational demands of the two processes differ substantially. Gradient navigation relies on continuous, real-time evaluation and comparison of present and past sensory input, whereas classical learning depends on the storage and retrieval of past associative experiences. Stimulus specific information is initially processed in peripheral sensory neurons, but subsequently converges onto shared computational nodes that encode modality-independent stimulus valence. Rather than maintaining dedicated circuits for each sensory modality or behavioral function, the larval brain appears to process diverse sensory inputs through a common evaluative framework (Eschbach et al., 2025). Consequently, different functional requirements between behaviors might arise not only from distinct circuit architectures, such as for DAN-c1, but also from differences in temporal dynamics and contextual modulation.

## Methods

### Animal stocks and husbandry

Flies were raised on a standard cornmeal diet under controlled conditions (25 °C, 60% relative humidity, 12 h light/12 h dark cycle). Egg laying was allowed for 48h, after which adults were removed from the vials. Larvae were allowed to develop for 4-6 days, and early third-instar individuals were used for experiments.

### Transgenic lines

Mutant lines and GAL4/UAS crosses were maintained under the same rearing conditions as mentioned above. Crosses were set up with a controlled number of adults to ensure comparable larval density during development. Mutant lines were not backcrossed to the WT background. Experimental crosses were established by combining at least 10 GAL4-driver line males with 20 *UAS-hid/reaper* female virgins. Control crosses consisted of both the UAS-hid/reaper and the GAL4-driver line crossed to WT, using similar sex ratios. WT experiments were performed using the Canton-S strain. Fly stocks were obtained from the Bloomington *Drosophila* Stock Center (BDSC) or provided by other labs (see table S1).

### Behavioral experiments

#### Gradient

A 2% agarose solution was prepared by dissolving 4 g of agarose in 200 ml of distilled water and heating the mixture until the agar was fully dissolved. To prepare the taste solutions, 18 g of fructose (Sigma-Aldrich, CAS #57-48-7) or 5.85 g of NaCl (Sigma-Aldrich, CAS #7647-14-5) was added to 100 ml of the agarose solution to obtain a 1M fructose or salt solution respectively. For other concentrations, 9 g and 36 g of fructose were used to prepare 0.5 M and 2 M fructose solutions, respectively, while 2.92 g and 11.7 g of NaCl were used to prepare 0.5 M and 2 M salt solutions.

Behavioral experiments were conducted in a 25 cm x 25 cm assay arena. To make the fructose or salt gradients, the arena was tilted by placing a 50 ml Falcon tube lid under one edge. Subsequently, 100 ml of the agarose solution containing fructose or salt was poured onto the tilted plate and allowed to solidify for around 10 minutes. After solidification, the plate was returned to a horizontal position, and an additional 100 ml of plain agarose solution was poured on top. This layer was allowed to solidify for another 10 minutes. The plate was then covered, inverted, and left overnight (15-18 hours) to allow diffusion of fructose or salt and the establishment of a taste gradient. Before the experiment, the plate was re-inverted and placed in the light-tight box, maintained at a constant temperature of 25°C and 60% humidity.

#### Strip

A 2% agarose solution was prepared by dissolving 4 g of agarose in 100 ml of distilled water and heating the mixture until the agar was fully dissolved. The solution was then poured onto the 25 cm x 25 cm assay plate and allowed to solidify for approximately 10 minutes. Once solidified, a 5 cm-wide strip of agarose was carefully removed from one side of the plate. Into the gap, 15 ml of 1 M fructose solution was added and allowed to solidify.

For all behavioral experiments, early third-instar larvae of a similar size were used (4-6 days after egg laying). Larvae were collected from fly vials and rinsed with distilled water to remove all traces of food. An individual larva was then placed in the center of the assay arena, and its behavior was recorded for 15 minutes using a Basler camera (acA2040-90umNIR) equipped with a lens (Kowa Lens LM16HC F1.4 f15mm1”). Videos were acquired at 1fps with a red light filter (Edmund optics #89-837) positioned above the arena.

### Behavioral and Statistical Analysis

Videos were processed using the open-source tracking software TRex (Walter & Couzin, 2021) to extract larval positions (X/Y) and turn angle. The preference index was calculated as:

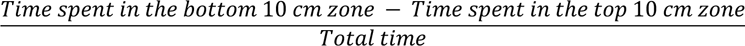

Larvae that reached the high concentration (bottom 5 cm) zone in fructose gradients and low concentration (top 5 cm) zone in salt gradients were considered to be successful. Each larva was classified based on whether it reached the high concentration zone, the low concentration zone, or neither zone during this period. The percentage of larvae in each category was then calculated for each condition. Larvae that ever reached the successful zone during the experiment were assigned to the high concentration zone for the fructose gradients and low concentration zone for the salt gradients respectively. For fructose gradients, the travel time was defined as the time required for a larva to first reach the high concentration zone, whereas for salt gradients it was defined as the time required to first reach the low concentration zone. Dwell time was defined as the cumulative time spent within the respective target zone after entry. To compare metrics across conditions we used a bootstrapped confidence-interval test and considered results significant when the effect size was moderate (Cohen’s d > 0.35, Data S1).

Speed was computed as the displacement in X and Y coordinates sampled at 1 s intervals. Stopping events were counted as frames with instantaneous speed < 0.02 cm/s, and stop frequency was reported as the number of such frames per trial. Turn angles were calculated from the tracking data. Runs were defined as continuous segments between two frames in which the turn angle exceeded 0.25 rad; run frequency was the number of runs per trial and run length was the distance covered in each run.

We calculated these locomotor parameters overall and separately for movements up and down the arena, both until larvae reached the high fructose concentration or the low salt concentration (during navigation) and for the period after they reached the respective zone (post successful navigation). To compare metrics across conditions we used a bootstrapped confidence-interval test and considered results significant when the effect size was small as the data was very variable (Cohen’s d > 0.20, Data S2).

## Author Contributions

Conceptualization: A.M., K.V. Methodology: A.M. Software: A.M. Validation: A.M. Formal analysis: A.M. Investigation: A.M. Data curation: A.M. Resources: K.V. Writing - original draft: A.M., K.V. Writing - review and editing: A.M., K.V. Visualization: A.M. Supervision: K.V. Project administration: K.V. Funding acquisition: K.V.

## Acknowledgments

We would like to thank Andreas Thum, Einat Couzin-Fuchs and Iain Couzin for valuable discussions. A.M. and K.V. were supported by the DFG German Research Foundation (EXC 2117-422037984, FOR5424 - 466488864).

## Supplementary Information

**Table S1:**
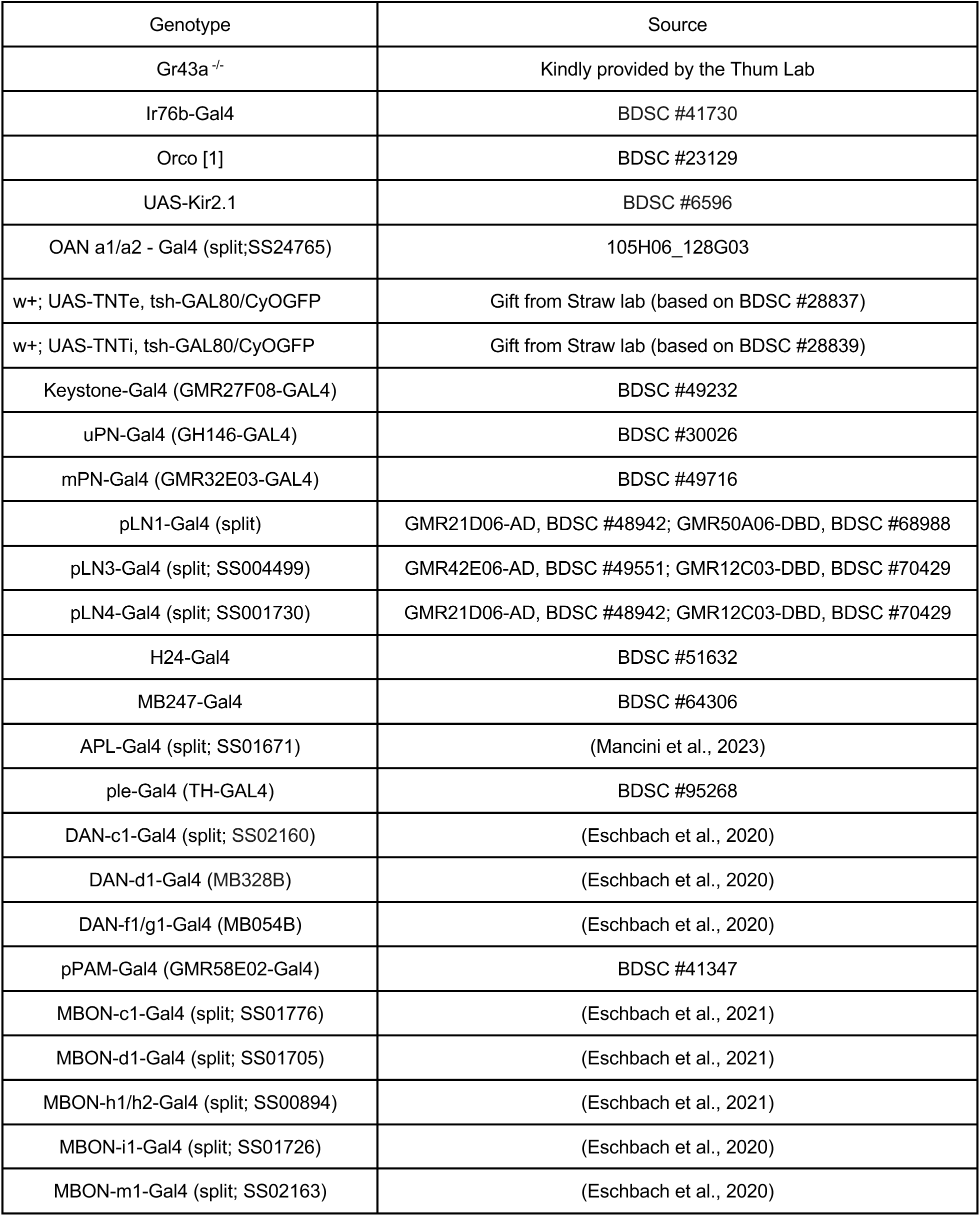
Fly lines used in this study.

**Fig S1.**
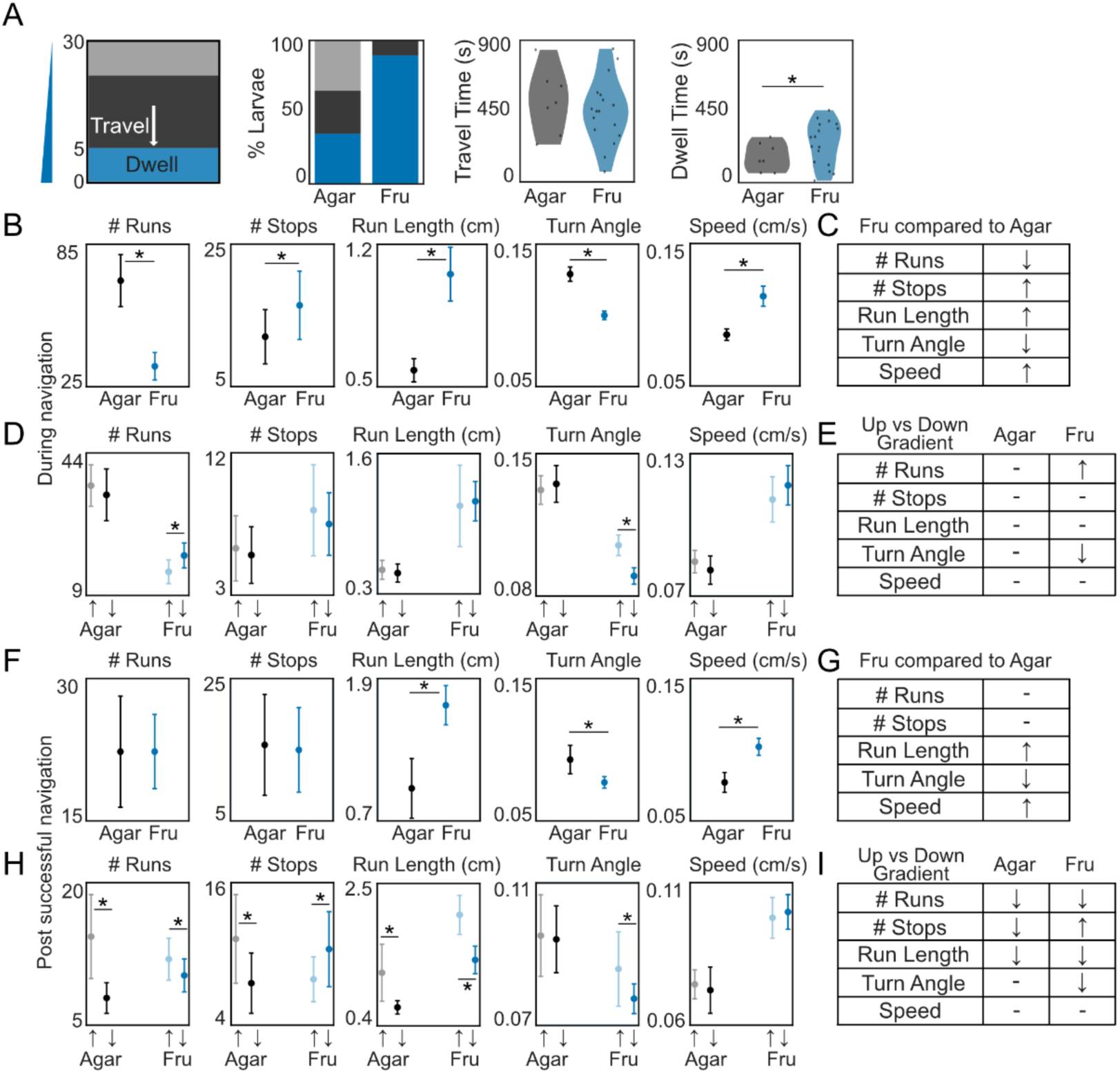
Larval behavior during fructose gradient navigation: (**A**) Percentage of larvae either reaching the bottom (blue), top (gray) or neither region (black) on agar and fructose gradients. Time to reach the high-fructose zone and time spent in the zone on agar (black) and fructose gradients (blue) (Bootstrapped confidence interval (CI) test = * Cohen’s d > 0.35, see Data S1). (**B**) Locomotor parameters before reaching either top or bottom of the arena. (**C**) Summary of larval behavior on fructose compared to agar (↑ : increase, ↓ : decrease; – : no difference). (**D**) Parameters during movement toward low fructose (dim) vs toward high fructose (bright) prior to reaching either top or bottom of the arena. (**E**) Summary of larval behavioral modulation. (**F**) Parameters after reaching high fructose regions. (**G**) Summary of larval behavior on fructose compared to agar. (**H**) Locomotor parameters while larvae move in and against the gradient direction. (**I**) Summary of larval behavioral modulation (Bootstrapped confidence interval (CI) test = * Cohen’s d > 0.2, see Data S2).

**Fig S2.**
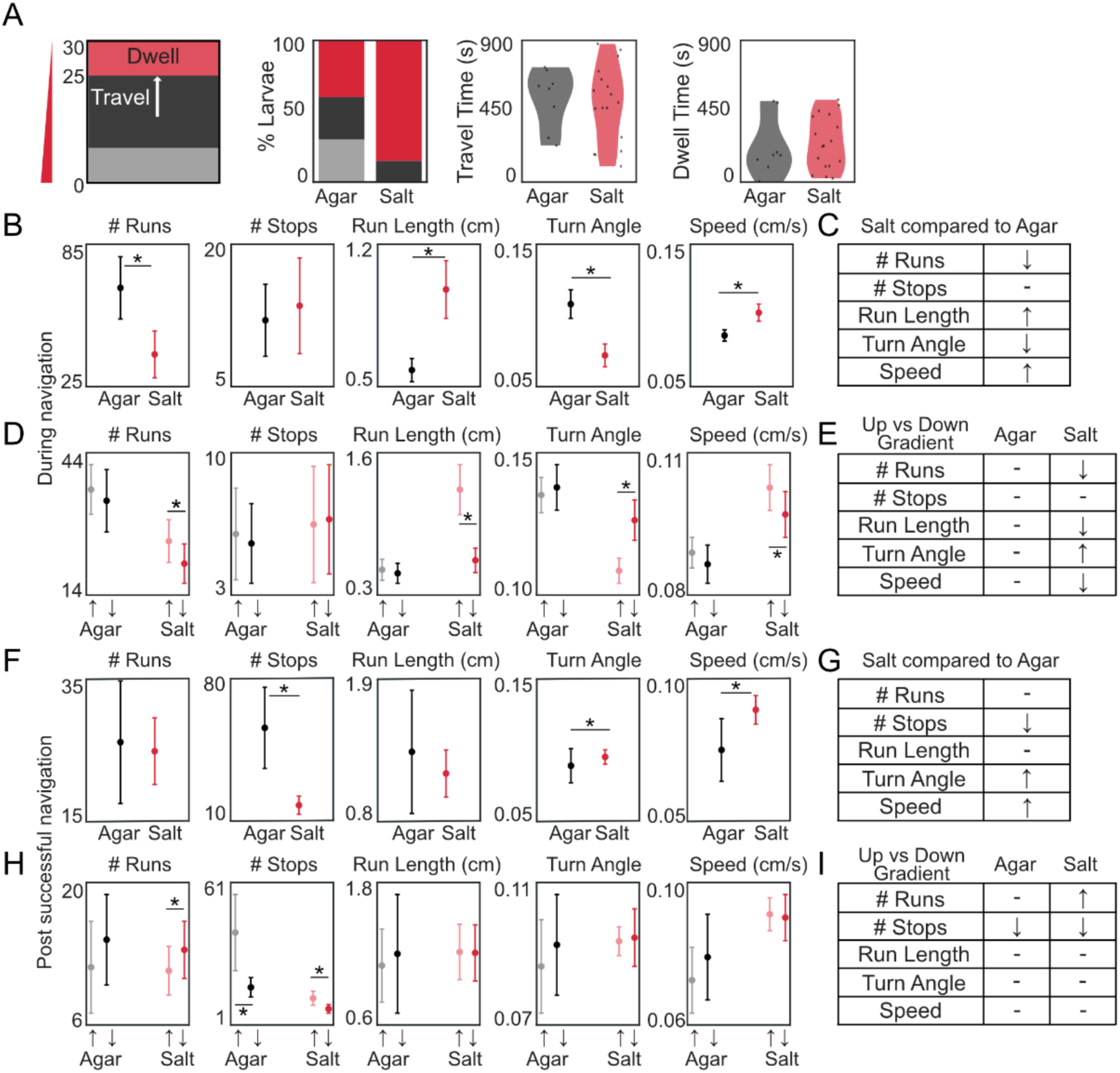
Larval behavior during salt gradient navigation: (**A**) Percentage of larvae either reaching the bottom (blue), top (gray) or neither region (red) on agar and salt gradients. Time to reach the high-fructose zone and time spent in the zone on agar (black) and salt gradients (red) (Bootstrapped confidence interval (CI) test = * Cohen’s d > 0.35, see Data S1). (**B**) Locomotor parameters before reaching either top or bottom of the arena. (**C**) Summary of larval behavior on salt compared to agar (↑ : increase, ↓ : decrease; – : no difference). (**D**) Parameters during movement toward low salt (dim) vs toward high salt (bright) prior to reaching either top or bottom of the arena. (**E**) Summary of larval behavioral modulation. (**F**) Parameters after reaching low salt regions. (**G**) Summary of larval behavior on salt compared to agar. (**H**) Locomotor parameters while larvae move in and against the gradient direction. (**I**) Summary of larval behavioral modulation (Bootstrapped confidence interval (CI) test = * Cohen’s d > 0.2, see Data S2).

**Fig S3.**
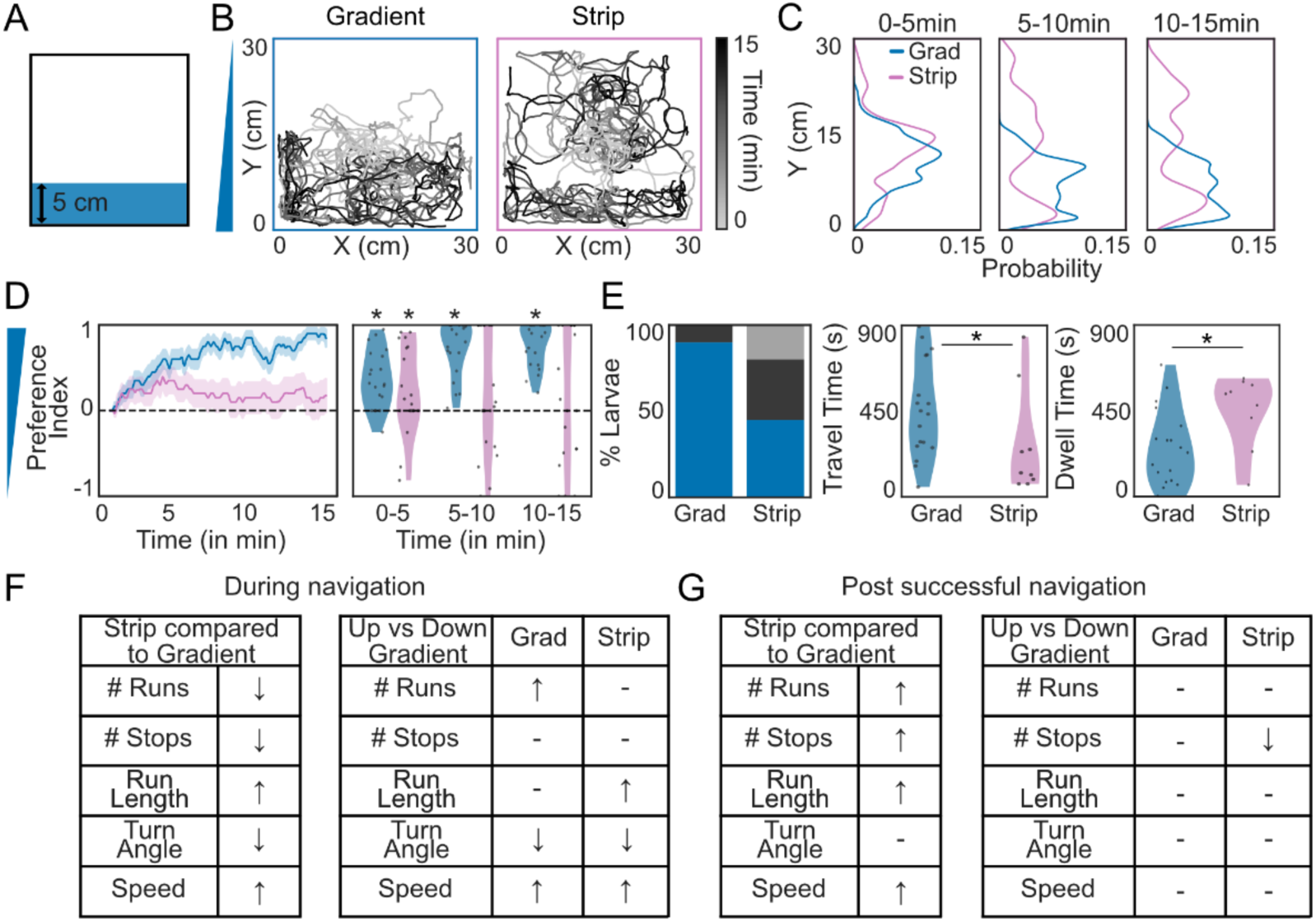
Fructose gradient navigation requires continuous gradient information: (**A**) Schematic of the arena with a 5 cm-wide 1M fructose strip (rest: agar). (**B**) Sample trajectories of individual larvae on fructose gradient (blue) and strip control (pink) (N = 20 larvae per condition). (**C**) Larval distribution along Y-axis. (**D**) Preference index over time. (mean: solid line, SEM: shaded) Binned preference index (5-min intervals, shown as violin plots; each dot: individual larva; bootstrapped CI * = Cohen’s d > 0.35, see Data S1). (**E**) Percentage of larvae either reaching the bottom, top or neither region, time to reach the high-fructose zone and time spent in the zone. (**F**) Locomotor parameters before reaching either end of the arena (↑ : increase, ↓ : decrease; – : no difference). Parameters during movement with vs against gradient direction. (**G**) Locomotor parameters after reaching high fructose regions. Parameters during movement with vs against gradient direction (Bootstrapped CI test = * Cohen’s d > 0.2, see Data S2).

**Fig S4.**
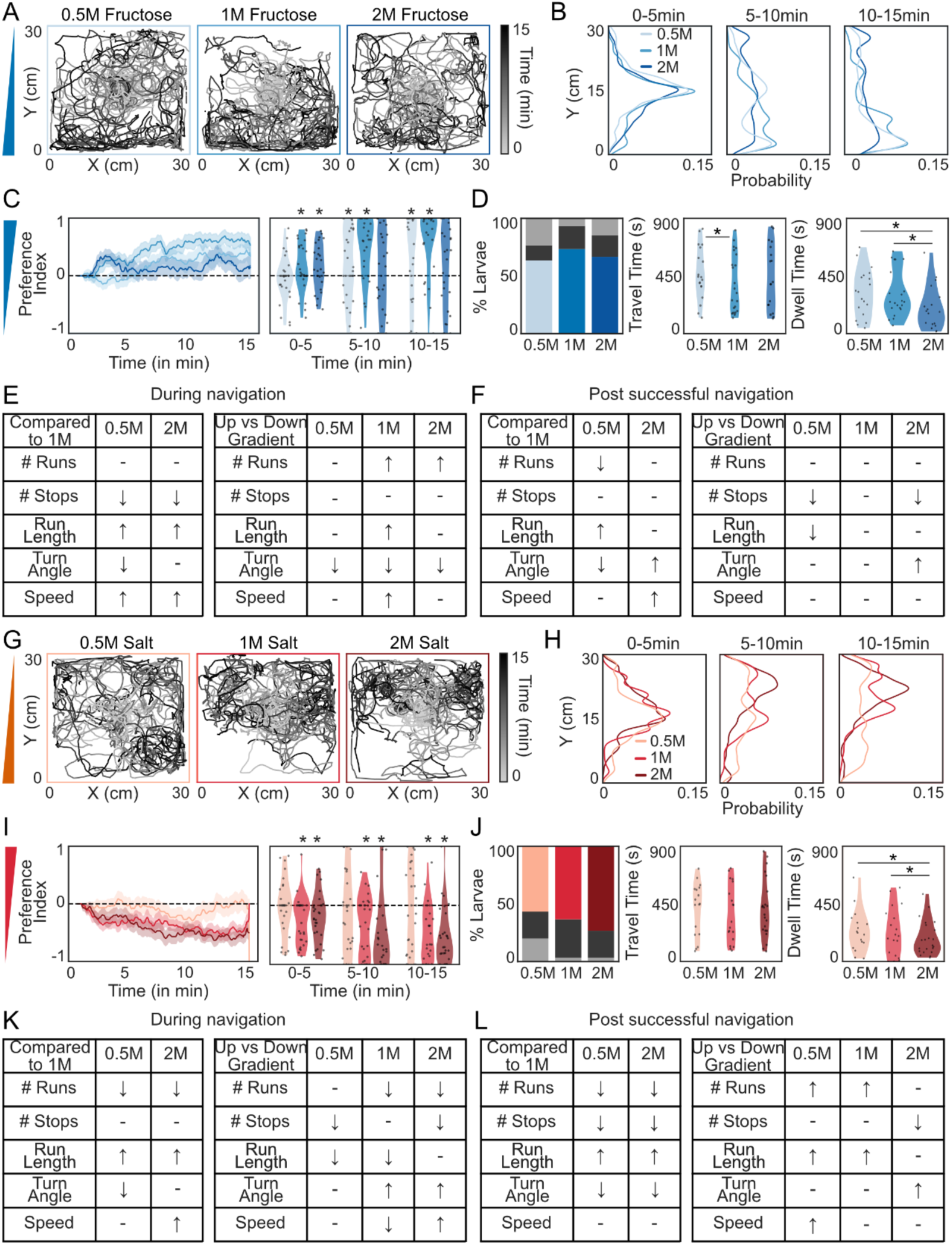
Concentration-dependent taste gradient navigation: (**A**) Trajectories of individual larvae on 0.5M fructose (light blue), 1M fructose (blue) and 2M fructose (dark blue) gradients (N = 30 per condition). (**B**) Larval distribution along Y-axis. (**C**) Preference index over time (solid line: mean; shaded: SEM) and binned preference index (5min intervals, dots: individual larvae). (**D**) Percentage of larvae either reaching the bottom, top or neither region; time to reach the high-fructose zone and time spent in the zone (Bootstrapped CI test; Cohen’s d > 0.35, see Data S1). (**E**) Locomotor parameters before reaching either top or bottom of the arena (↑ : increase, ↓ : decrease; – : no difference) and when moving with vs against the gradient (Bootstrapped CI test = * Cohen’s d > 0.2, see Data S2). (**F**) Locomotor parameters after successful navigation and when moving with vs against the gradient. (**G**) Trajectories on 0.5M salt (light red), 1M salt (red) and 2M salt (dark red) gradients (N = 30). (**H**) Distribution of larvae. (**I**) Preference index over time and binned preference index (**J**) Percentage of larvae either reaching the bottom, top or neither region; time to reach the low-salt zone and time spent in the zone. (**K**) Locomotor parameters before reaching either top or bottom of the arena and when moving with vs against the gradient. (**L**) Locomotor parameters after successful navigation and when moving with vs against the gradient.

**Fig S5.**
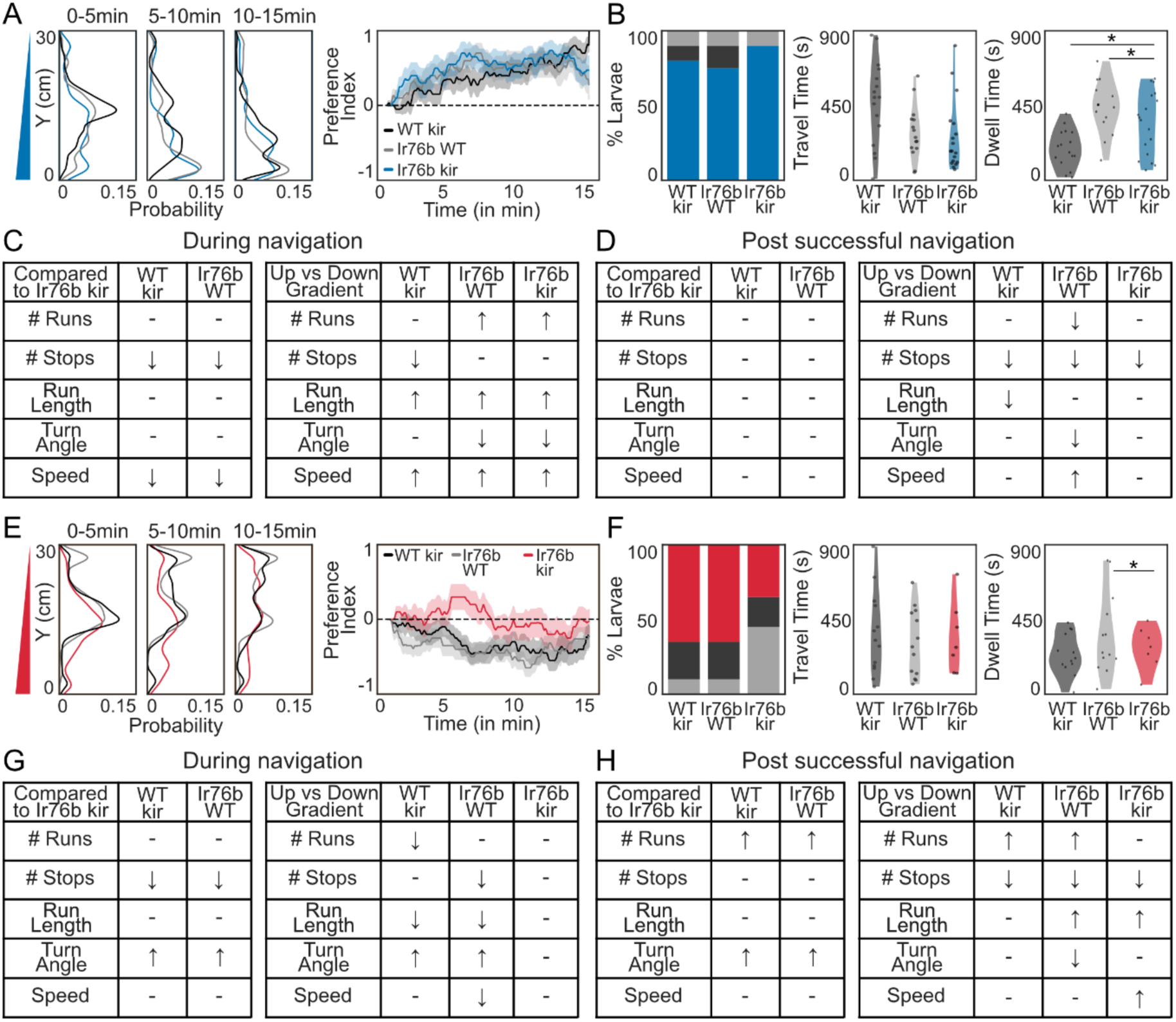
Larval behavior of WT/kir, Ir76b/WT and Ir76b/kir larvae during fructose and salt gradient navigation: (**A**) Distribution of WT/kir (black), Ir76b/WT (gray) and Ir76b/kir (blue) larvae on fructose gradient along the Y-axis. Preference index over time (solid line: mean; shaded: SEM). (**B**) Percentage of larvae either reaching the bottom, top or neither region; time to reach the high-fructose zone and time spent in the zone (Bootstrapped CI test = Cohen’s d > 0.35, see Data S1). (**C**) Locomotor parameters before reaching either top or bottom of the arena (↑ : increase, ↓ : decrease; – : no difference). Locomotor parameters when moving with vs against the gradient (Bootstrapped CI test = * Cohen’s d > 0.2, see Data S2). (**D**) Locomotor parameters after successful navigation and when moving with vs against the gradient. (**E**) Distribution of WT/kir (black), Ir76b/WT (gray) and Ir76b/kir (red) larvae on salt gradient along the Y-axis. Preference index over time. (**F**) Percentage of larvae either reaching the bottom, top or neither region; time to reach the low-salt zone and time spent in the zone. (**G**) Locomotor parameters before reaching either top or bottom of the arena. Locomotor parameters when moving with vs against the gradient. (**H**) Locomotor parameters after successful navigation and when moving with vs against the gradient.

**Fig S6.**
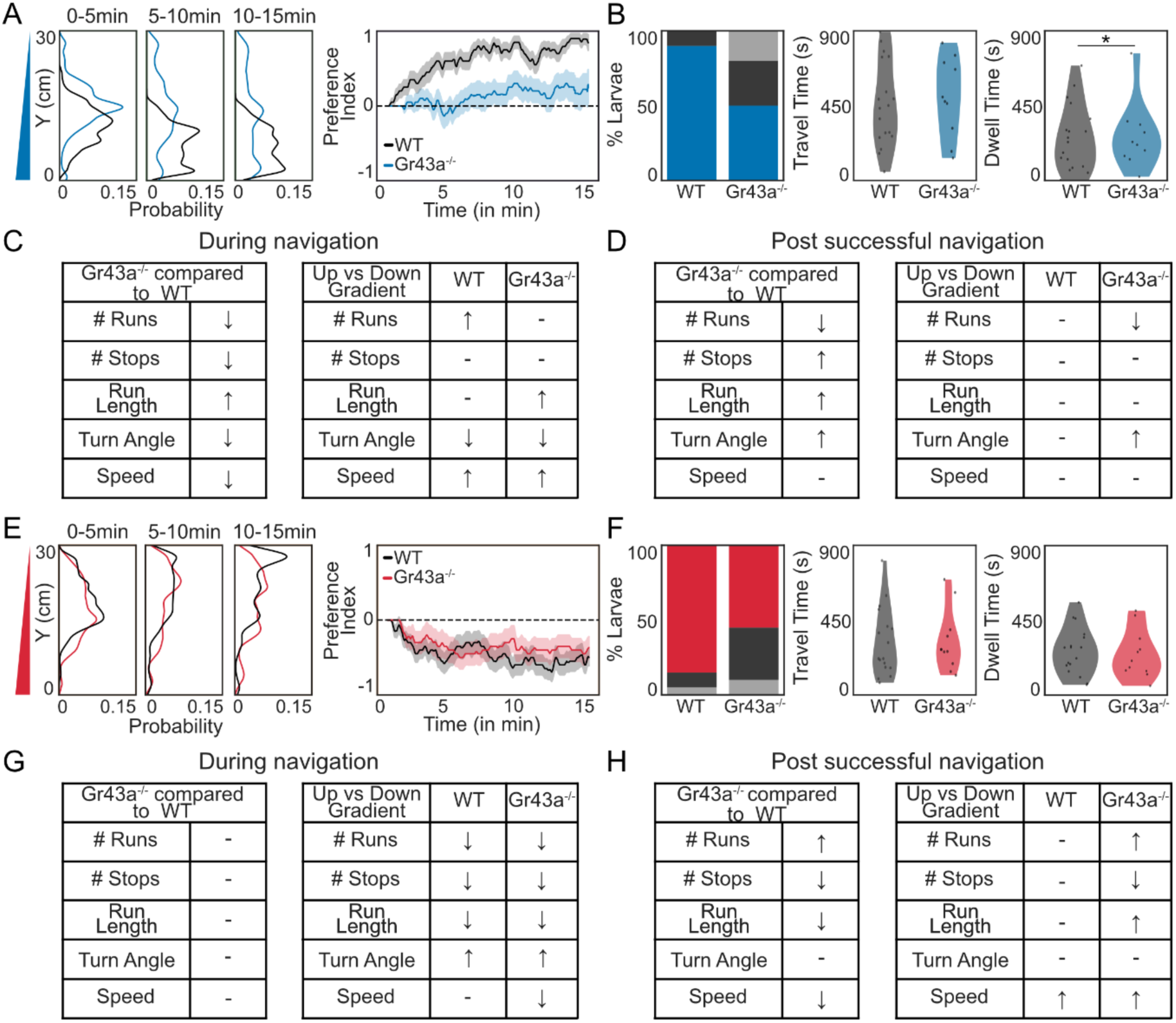
Larval behavior of WT and Gr43a^-/-^ larvae during fructose and salt gradient navigation: (**A**) Distribution of WT (black) and Gr43a^-/-^ (blue) larvae on fructose gradient along the Y-axis. Preference index over time (solid line: mean; shaded: SEM). (**B**) Percentage of larvae either reaching the bottom, top or neither region; time to reach the high-fructose zone and time spent in the zone (Bootstrapped CI test = Cohen’s d > 0.35, see Data S1). (**C**) Locomotor parameters before reaching either top or bottom of the arena (↑ : increase, ↓ : decrease; – : no difference). Locomotor parameters when moving with vs against the gradient (Bootstrapped CI test = * Cohen’s d > 0.2, see Data S2). (**D**) Locomotor parameters after successful navigation and when moving with vs against the gradient. (**E**) Distribution of WT (black) and Gr43a^-/-^ (red) larvae on salt gradient along the Y-axis. Preference index over time. (**F**) Percentage of larvae either reaching the bottom, top or neither region; time to reach the low-salt zone and time spent in the zone. (**G**) Locomotor parameters before reaching either top or bottom of the arena. Locomotor parameters when moving with vs against the gradient. (**H**) Locomotor parameters after successful navigation and when moving with vs against the gradient.

**Fig S7.**
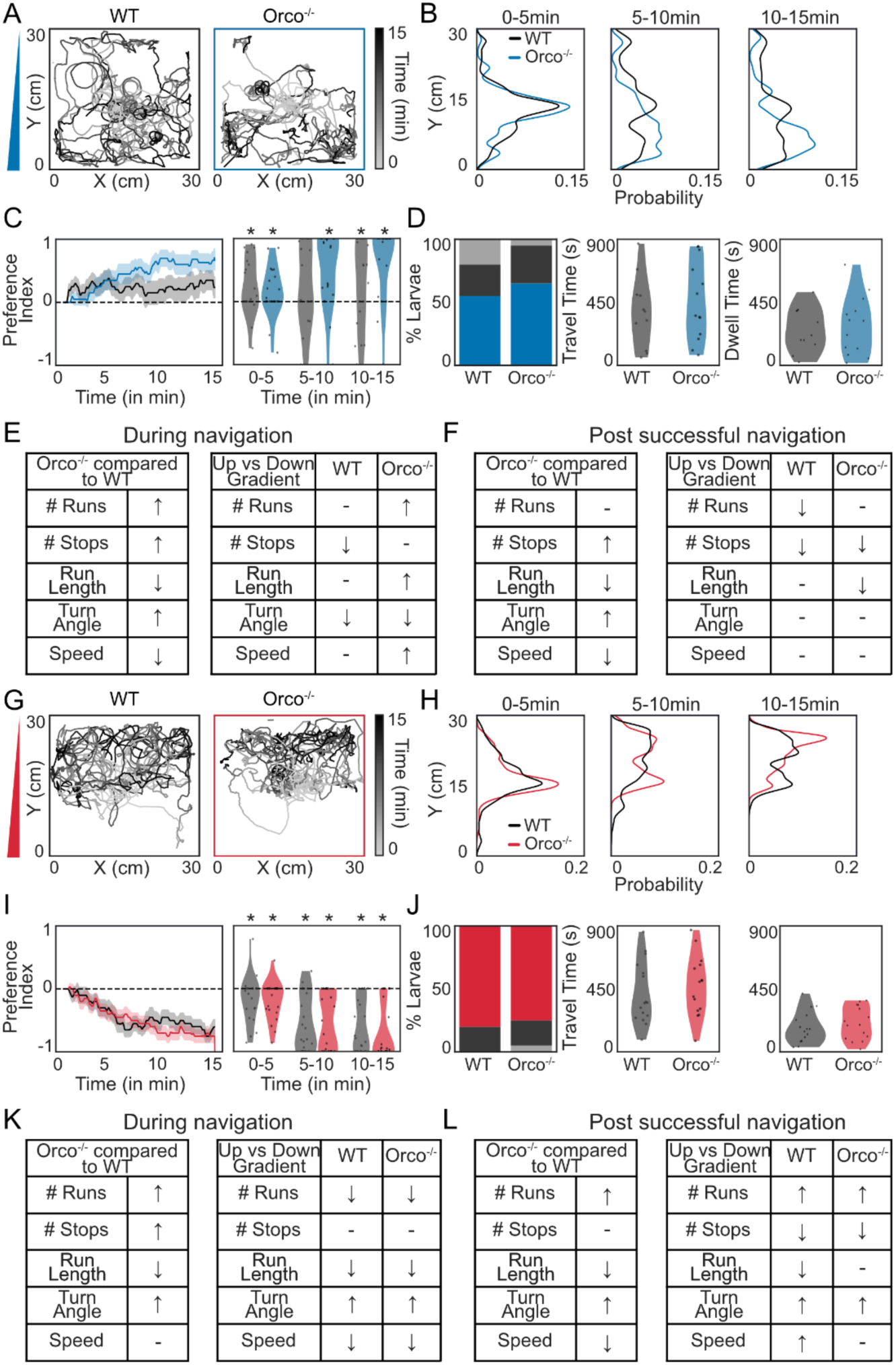
Olfactory cues are not needed for taste gradient navigation: (**A**) Trajectories of WT (black) and Orco^-/-^ (blue) during fructose gradient navigation (N = 20). (**B**) Larval distribution along the Y-axis. (**C**) Preference index over time (solid line: mean; shaded: SEM) and binned preference index (5min intervals, dots: individual larvae). (**D**) Percentage of larvae either reaching the bottom, top or neither region; time to reach the high-fructose zone and time spent in the zone (Bootstrapped CI test = Cohen’s d > 0.35, see Data S1). (**E**) Locomotor parameters before reaching either top or bottom of the arena (↑ : increase, ↓ : decrease; – : no difference). Locomotor parameters when moving with vs against the gradient (Bootstrapped CI test = * Cohen’s d > 0.2, see Data S2). (**F**) Locomotor parameters after successful navigation and when moving with vs against the gradient. (**G**) Trajectories of WT (black) and Orco^-/-^ (red) during salt gradient navigation (N = 20). (**H**) Larval distribution along the Y-axis. (**I**) Preference index over time and binned preference index. (**J**) Percentage of larvae either reaching the bottom, top or neither region; time to reach the low-salt zone and time spent in the zone. (**K**) Locomotor parameters before reaching either top or bottom of the arena compared to WT. Locomotor parameters when moving with vs against the gradient. (**L**) Locomotor parameters after successful navigation and when moving with vs against the gradient.

**Fig S8.**
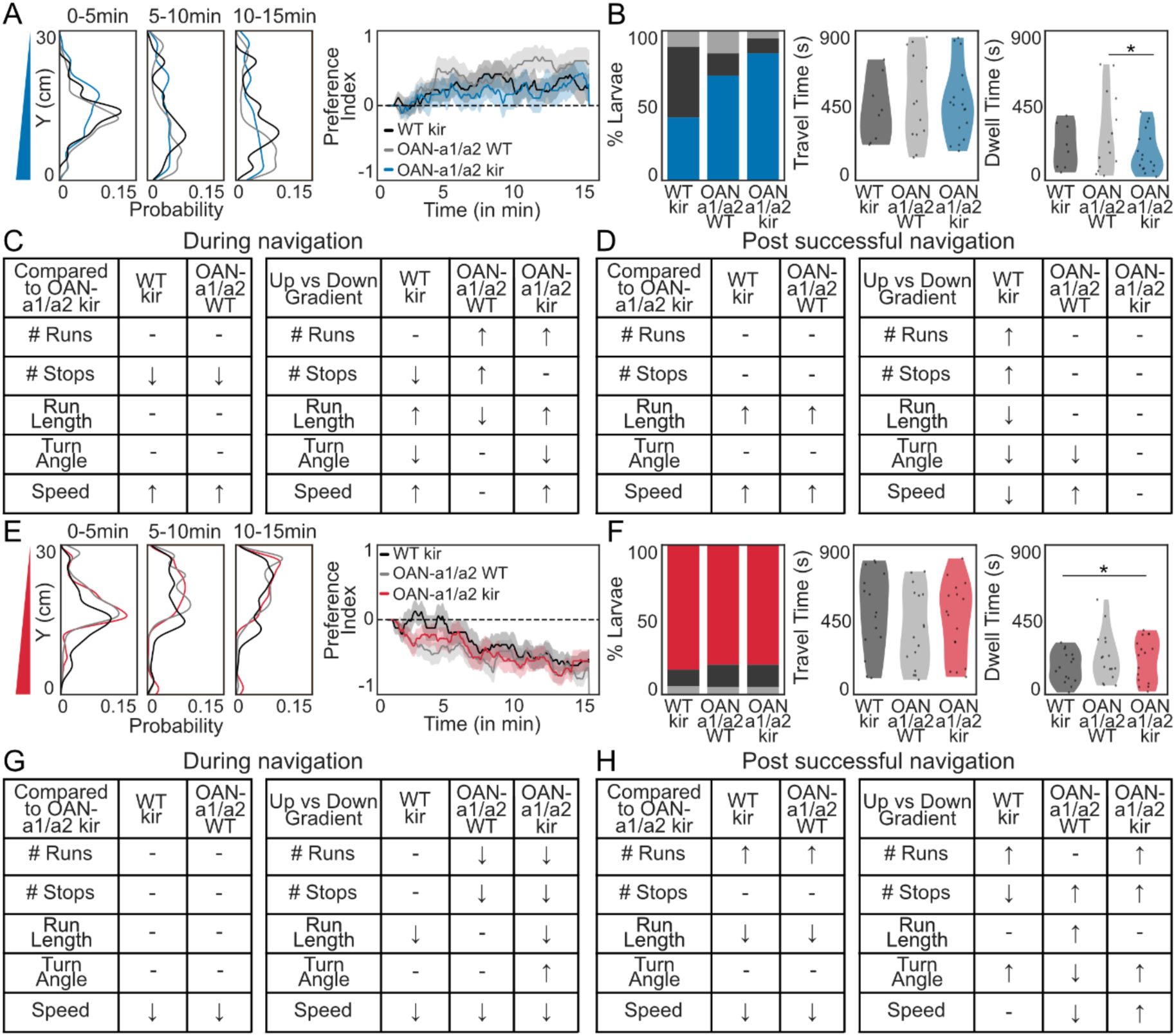
Larval behavior of WT-kir, OAN-a1/a2/WT and OAN-a1/a2/kir larvae during fructose and salt gradient navigation: (**A**) Distribution of WT/kir (black), OAN-a1/a2/WT (gray) and OAN-a1/a2/kir (blue) larvae on fructose gradient along the Y-axis. Preference index over time (solid line: mean; shaded: SEM). (**B**) Percentage of larvae either reaching the bottom, top or neither region; time to reach the high-fructose zone and time spent in the zone (Bootstrapped CI test = Cohen’s d > 0.35, see Data S1). (**C**) Locomotor parameters before reaching either top or bottom of the arena (↑ : increase, ↓ : decrease; – : no difference). Locomotor parameters when moving with vs against the gradient (Bootstrapped CI test = * Cohen’s d > 0.2, see Data S2). (**D**) Locomotor parameters after successful navigation and when moving with vs against the gradient. (**E**) Distribution of WT/kir (black), OAN-a1/a2 WT (gray) and OAN-a1/a2/kir (red) larvae on salt gradient along the Y-axis. Preference index over time. (**F**) Percentage of larvae either reaching the bottom, top or neither region; time to reach the low-salt zone and time spent in the zone. (**G**) Locomotor parameters before reaching either top or bottom of the arena. Locomotor parameters when moving with vs against the gradient. (**H**) Locomotor parameters after successful navigation and when moving with vs against the gradient.

**Fig S9.**
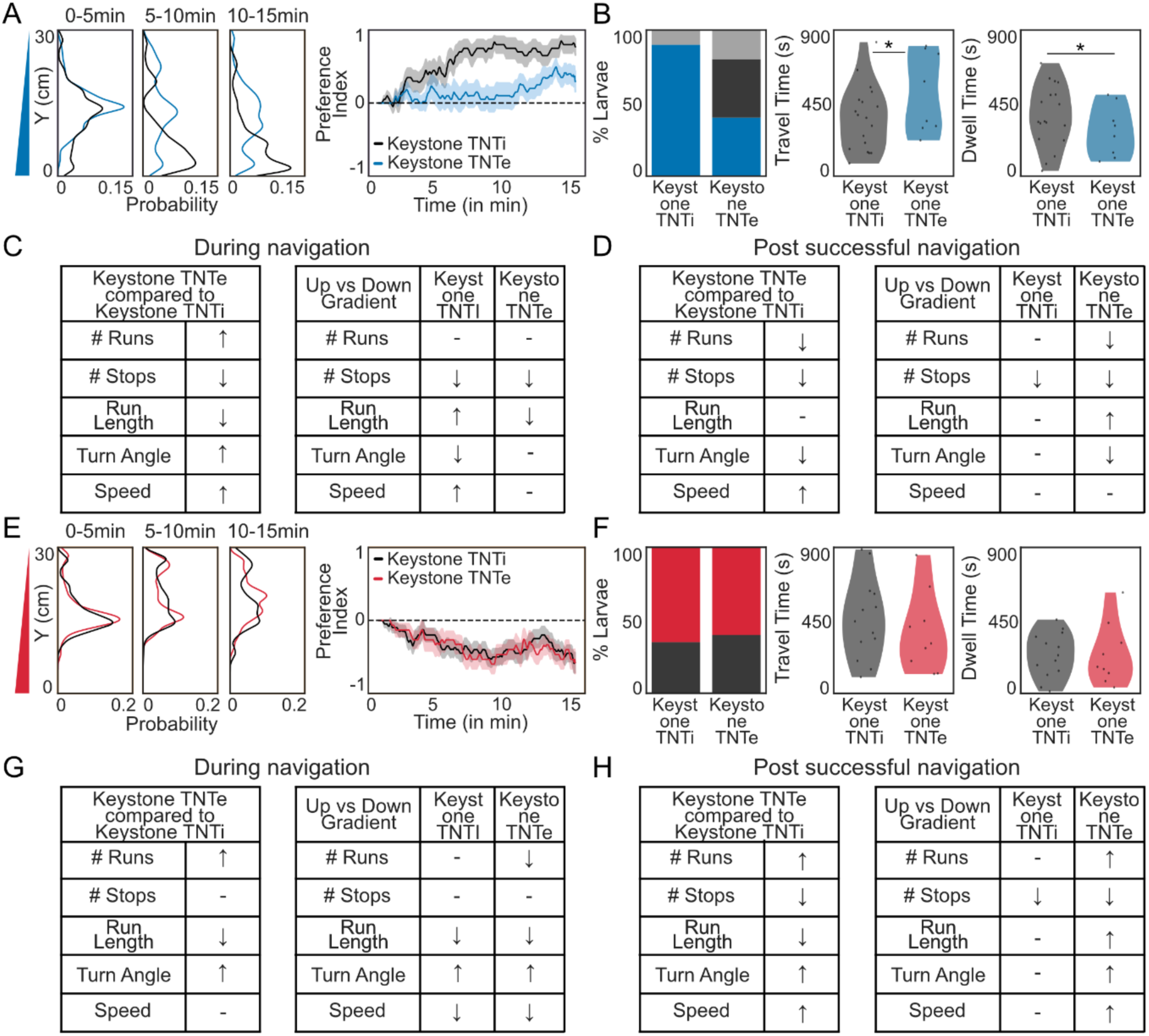
Larval behavior of Keystone/TNTi and Keystone/TNTe larvae during fructose and salt gradient navigation: (**A**) Distribution of Keystone/TNTi (black) and Keystone/TNTe (blue) larvae on fructose gradient along the Y-axis. Preference index over time (solid line: mean; shaded: SEM). (**B**) Percentage of larvae either reaching the bottom, top or neither region; time to reach the high-fructose zone and time spent in the zone (Bootstrapped CI test = Cohen’s d > 0.35, see Data S1). (**C**) Locomotor parameters before reaching either top or bottom of the arena (↑ : increase, ↓ : decrease; – : no difference). Locomotor parameters when moving with vs against the gradient (Bootstrapped CI test = * Cohen’s d > 0.2, see Data S2). (**D**) Locomotor parameters after successful navigation and when moving with vs against the gradient. (**E**) Distribution of Keystone/TNTi (black) and Keystone/TNTe (red) larvae on salt gradient along the Y-axis. Preference index over time. (**F**) Percentage of larvae either reaching the bottom, top or neither region; time to reach the low-salt zone and time spent in the zone. (**G**) Locomotor parameters before reaching either top or bottom of the arena. Locomotor parameters when moving with vs against the gradient. (**H**) Locomotor parameters after successful navigation and when moving with vs against the gradient.

**Fig S10.**
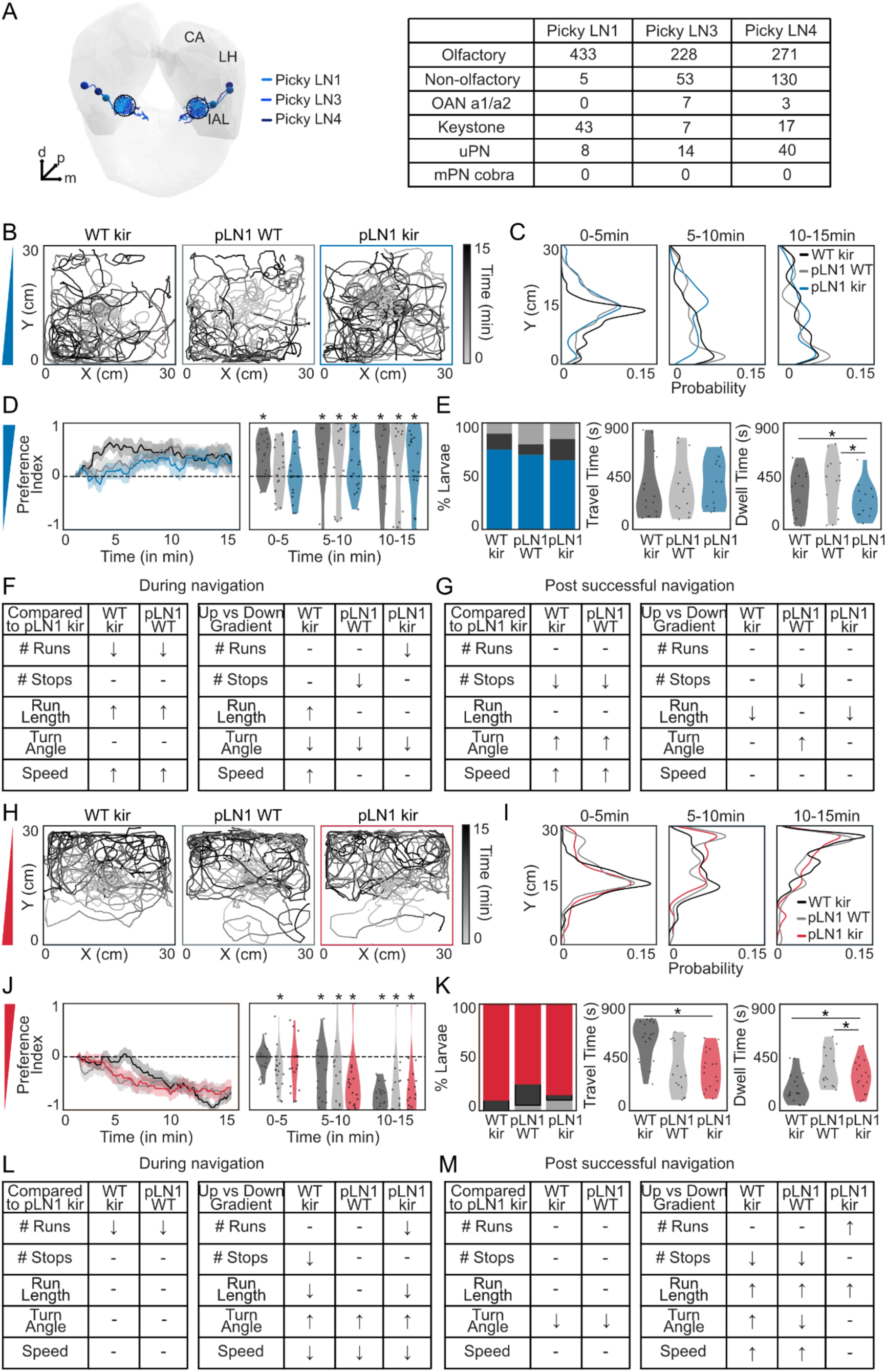
Picky local interneuron pLN1 are not involved in taste gradient navigation: (**A**) EM reconstructions of picky 1, picky 3, picky 4 neurons (larval antennal lobe (IAL), mushroom body calyx (CA), lateral horn (LH)). Olfactory, non-olfactory and projection neurons input to these neurons was calculated from EM data. (**B**) Trajectories of WT/kir (black), pLN1/WT (gray) and pLN1/kir (blue) during fructose gradient navigation (N = 20). (**C**) Larval distribution along the Y-axis. (**D**) Preference index over time (solid line: mean; shaded: SEM) and binned preference index (5min intervals, dots: individual larvae). (**E**) Percentage of larvae either reaching the bottom, top or neither region; time to reach the high-fructose zone and time spent in the zone (Bootstrapped CI test = Cohen’s d > 0.35, see Data S1). (**F**) Locomotor parameters before reaching either top or bottom of the arena (↑ : increase, ↓ : decrease; – : no difference). Locomotor parameters when moving with vs against the gradient (Bootstrapped CI test = * Cohen’s d > 0.2, see Data S2). (**G**) Locomotor parameters after successful navigation and when moving with vs against the gradient. (**H**) Trajectories of WT/ kir (black), pLN1/WT (gray) and pLN1/kir (red) during salt gradient navigation (N = 20). (**I**) Larval distribution along the Y-axis. (**J**) Preference index over time and binned preference index. (**K**) Percentage of larvae either reaching the bottom, top or neither region; time to reach the low-salt zone and time spent in the zone. (**L**) Locomotor parameters before reaching either top or bottom of the arena compared to WT. Locomotor parameters when moving with vs against the gradient. (**M**) Locomotor parameters after successful navigation and when moving with vs against the gradient.

**Fig S11.**
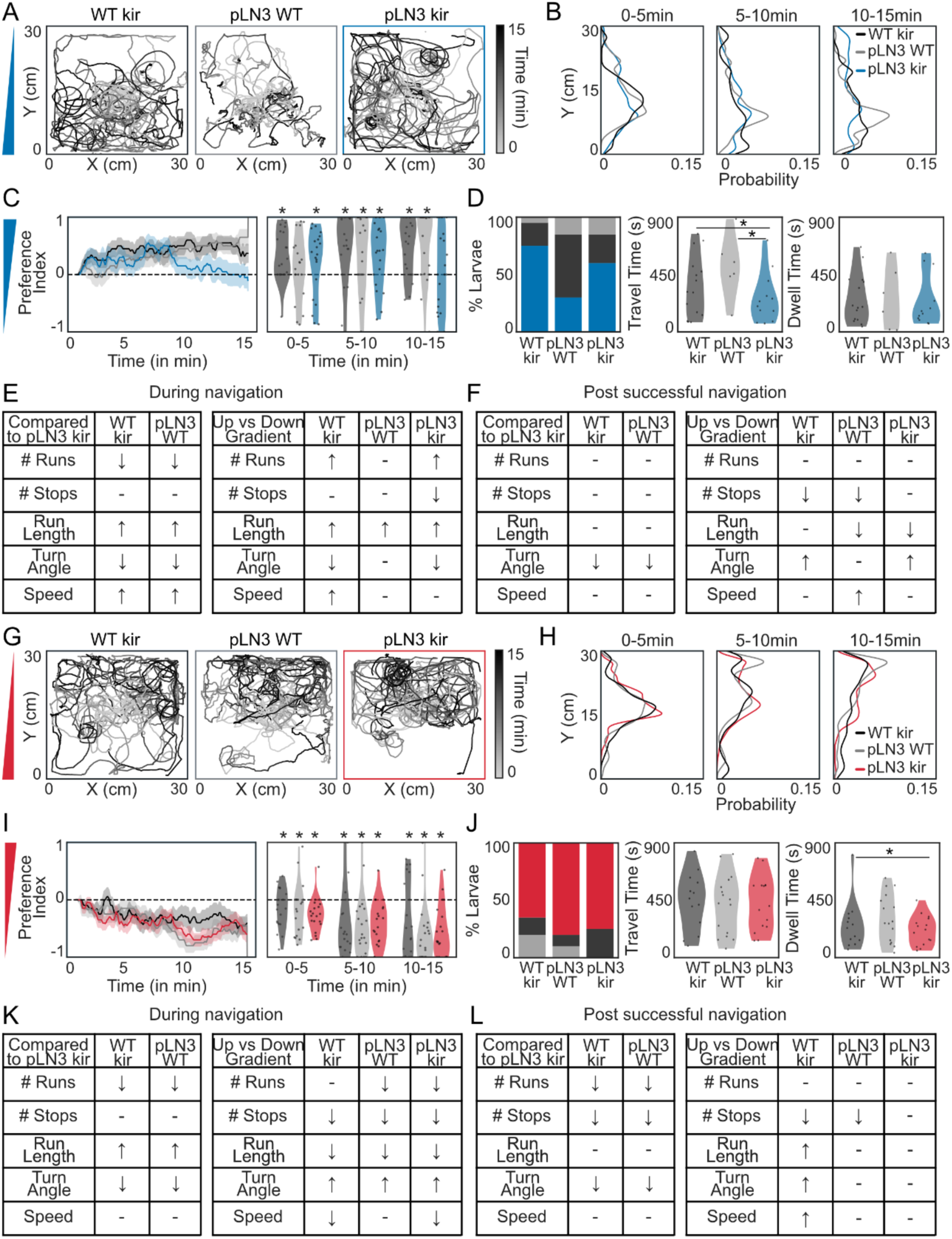
Picky local interneuron pLN3 are not involved in taste gradient navigation: (**A**) Trajectories of WT/kir (black), pLN3/WT (gray) and pLN3/kir (blue) during fructose gradient navigation (N = 20). (**B**) Larval distribution along the Y-axis. (**C**) Preference index over time (solid line: mean; shaded: SEM) and binned preference index (5min intervals, dots: individual larvae). (**D**) Percentage of larvae either reaching the bottom, top or neither region; time to reach the high-fructose zone and time spent in the zone (Bootstrapped CI test = Cohen’s d > 0.35, see Data S1). (**E**) Locomotor parameters before reaching either top or bottom of the arena (↑ : increase, ↓ : decrease; – : no difference). Locomotor parameters when moving with vs against the gradient (Bootstrapped CI test = * Cohen’s d > 0.2, see Data S2). (**F**) Locomotor parameters after successful navigation and when moving with vs against the gradient. (**G**) Trajectories of WT/kir (black), pLN3/WT (gray) and pLN3/kir (red) during salt gradient navigation (N = 20). (**H**) Larval distribution along the Y-axis. (I) Preference index over time and binned preference index. (**J**) Percentage of larvae either reaching the bottom, top or neither region; time to reach the low-salt zone and time spent in the zone. (**K**) Locomotor parameters before reaching either top or bottom of the arena compared to WT. Locomotor parameters when moving with vs against the gradient. (**L**) Locomotor parameters after successful navigation and when moving with vs against the gradient.

**Fig S12.**
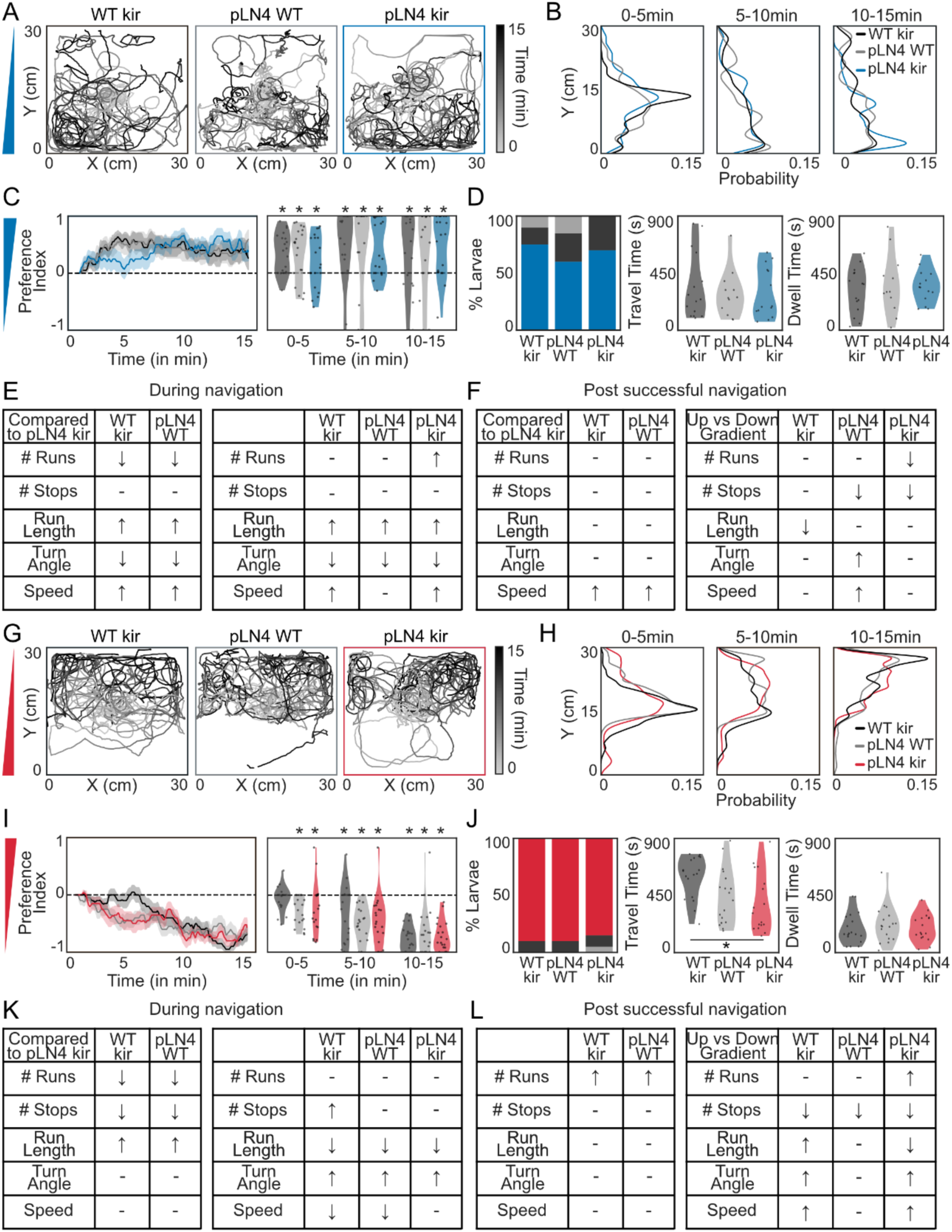
Picky local interneuron pLN4 are not involved in taste gradient navigation: (**A**) Trajectories of WT/kir (black), pLN4/WT (gray) and pLN4/kir (blue) during fructose gradient navigation (N = 20). (**B**) Larval distribution along the Y-axis. (**C**) Preference index over time (solid line: mean; shaded: SEM) and binned preference index (5min intervals, dots: individual larvae). (**D**) Percentage of larvae either reaching the bottom, top or neither region; time to reach the high-fructose zone and time spent in the zone (Bootstrapped CI test = Cohen’s d > 0.35, see Data S1). (**E**) Locomotor parameters before reaching either top or bottom of the arena (↑ : increase, ↓ : decrease; – : no difference). Locomotor parameters when moving with vs against the gradient (Bootstrapped CI test = * Cohen’s d > 0.2, see Data S2). (**F**) Locomotor parameters after successful navigation and when moving with vs against the gradient. (**G**) Trajectories of WT/kir (black), pLN4/WT (gray) and pLN4/kir (red) during salt gradient navigation (N = 20). (**H**) Larval distribution along the Y-axis. (I) Preference index over time and binned preference index. (**J**) Percentage of larvae either reaching the bottom, top or neither region; time to reach the low-salt zone and time spent in the zone. (**K**) Locomotor parameters before reaching either top or bottom of the arena compared to WT. Locomotor parameters when moving with vs against the gradient. (**L**) Locomotor parameters after successful navigation and when moving with vs against the gradient.

**Fig S13.**
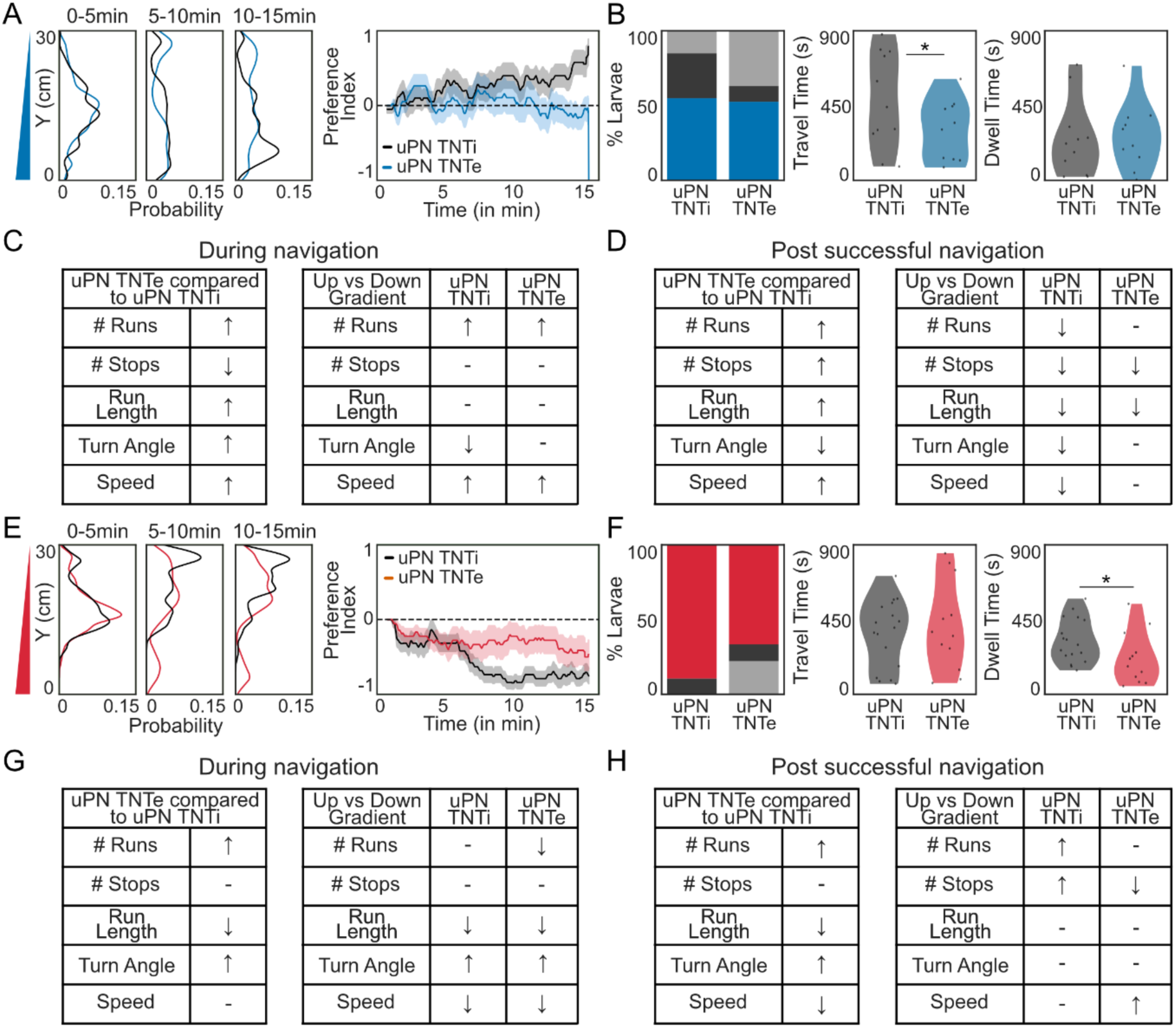
Larval behavior of uPN/TNTi and uPN/TNTe larvae during fructose and salt gradient navigation: (**A**) Distribution of uPN/TNTi (black) and uPN/TNTe (blue) larvae on fructose gradient along the Y-axis. Preference index over time (solid line: mean; shaded: SEM). (**B**) Percentage of larvae either reaching the bottom, top or neither region; time to reach the high-fructose zone and time spent in the zone (Bootstrapped CI test = Cohen’s d > 0.35, see Data S1). (**C**) Locomotor parameters before reaching either top or bottom of the arena (↑ : increase, ↓ : decrease; – : no difference). Locomotor parameters when moving with vs against the gradient. (**D**) Locomotor parameters after successful navigation and when moving with vs against the gradient (Bootstrapped CI test = * Cohen’s d > 0.2, see Data S2). (**E**) Distribution of uPN/TNTi (black) and uPN/TNTe (red) larvae on salt gradient along the Y-axis. Preference index over time. (**F**) Percentage of larvae either reaching the bottom, top or neither region; time to reach the low-salt zone and time spent in the zone. (**G**) Locomotor parameters before reaching either top or bottom of the arena. Locomotor parameters when moving with vs against the gradient. (**H**) Locomotor parameters after successful navigation and when moving with vs against the gradient.

**Fig S14.**
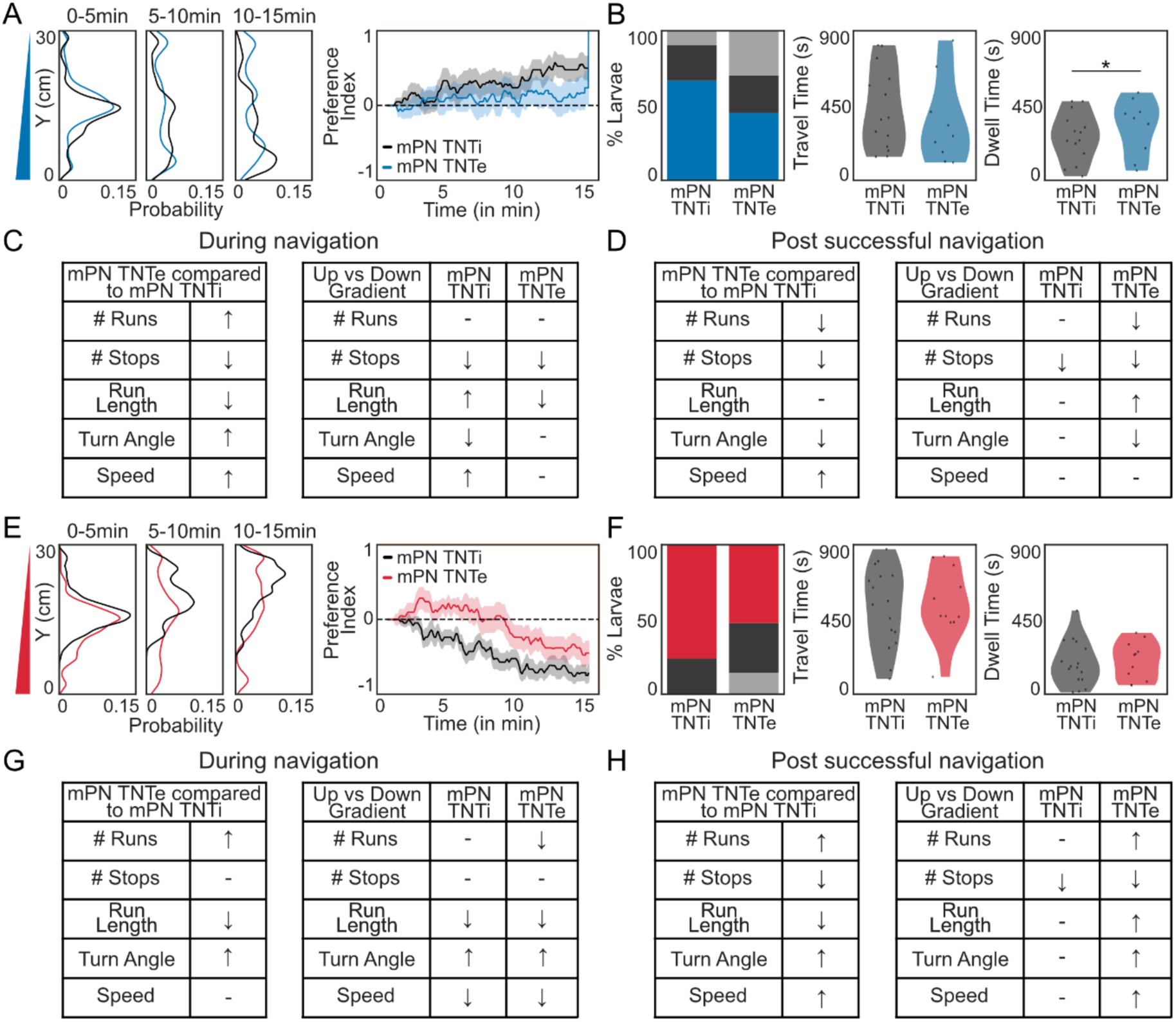
Larval behavior of mPN/TNTi and mPN/TNTe larvae during fructose and salt gradient navigation: (**A**) Distribution of mPN/TNTi (black) and mPN/TNTe (blue) larvae on fructose gradient along the Y-axis. Preference index over time (solid line: mean; shaded: SEM). (**B**) Percentage of larvae either reaching the bottom, top or neither region; time to reach the high-fructose zone and time spent in the zone. (Bootstrapped CI test = Cohen’s d > 0.35, see Data S1). (**C**) Locomotor parameters before reaching either top or bottom of the arena (↑ : increase, ↓ : decrease; – : no difference). Locomotor parameters when moving with vs against the gradient (Bootstrapped CI test = * Cohen’s d > 0.2, see Data S2). (**D**) Locomotor parameters after successful navigation and when moving with vs against the gradient. (**E**) Distribution of mPN/TNTi (black) and mPN/TNTe (red) larvae on salt gradient along the Y-axis. Preference index over time. (**F**) Percentage of larvae either reaching the bottom, top or neither region; time to reach the low-salt zone and time spent in the zone. (**G**) Locomotor parameters before reaching either top or bottom of the arena. Locomotor parameters when moving with vs against the gradient. (**H**) Locomotor parameters after successful navigation and when moving with vs against the gradient.

**Fig S15.**
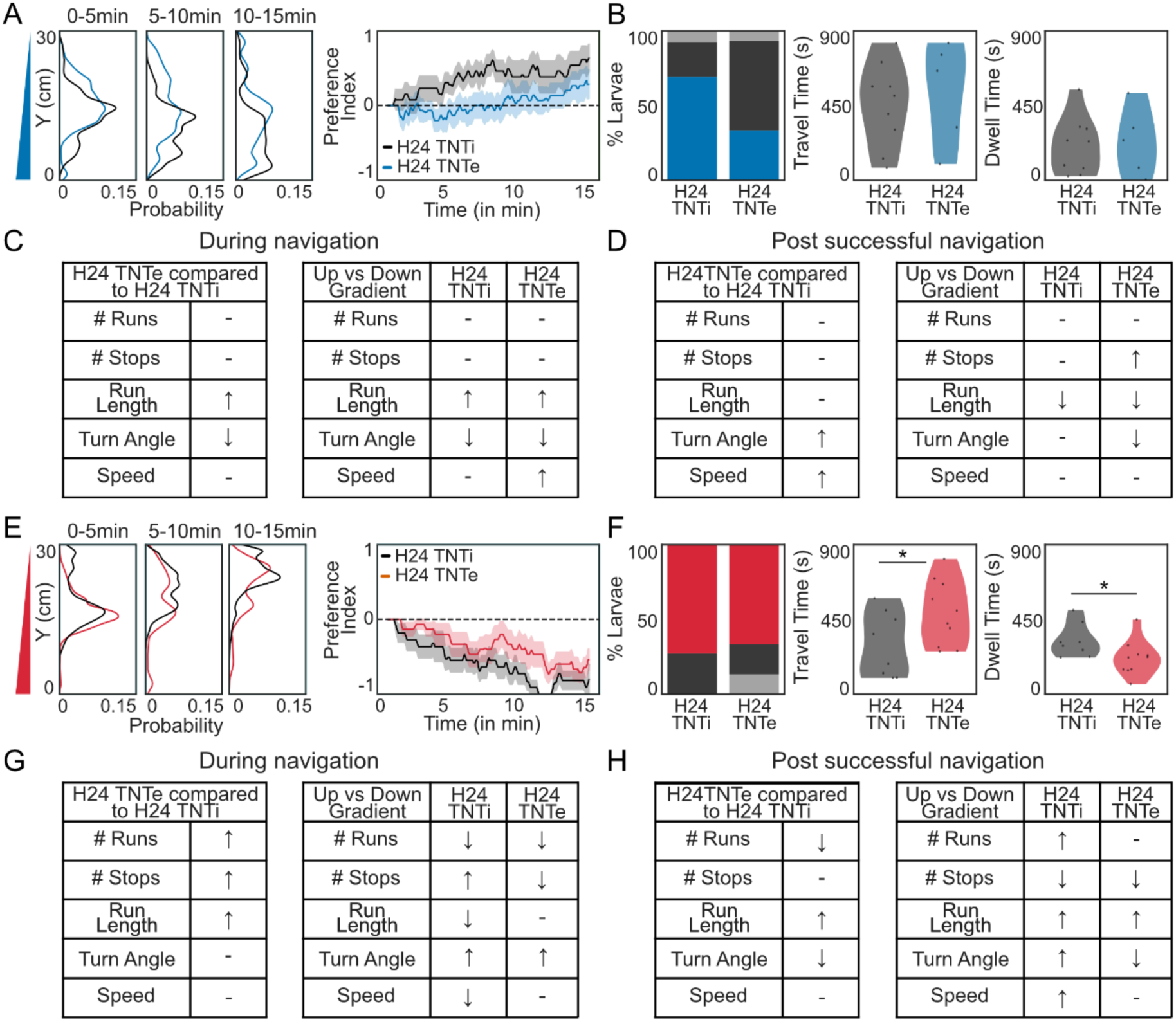
Larval behavior of H24/TNTi and H24/TNTe larvae during fructose and salt gradient navigation: (**A**) Distribution of H24/TNTi (black) and H24/TNTe (blue) larvae on fructose gradient along the Y-axis. Preference index over time (solid line: mean; shaded: SEM). (**B**) Percentage of larvae either reaching the bottom, top or neither region; time to reach the high-fructose zone and time spent in the zone (Bootstrapped CI test = Cohen’s d > 0.35, see Data S1). (**C**) Locomotor parameters before reaching either top or bottom of the arena compared to TNTi (↑ : increase, ↓ : decrease; – : no difference). Locomotor parameters when moving with vs against the gradient (Bootstrapped CI test = * Cohen’s d > 0.2, see Data S2). (**D**) Locomotor parameters after successful navigation and when moving with vs against the gradient. (**E**) Distribution of H24/TNTi (black) and H24/TNTe (red) larvae on salt gradient along the Y-axis. Preference index over time. (**F**) Percentage of larvae either reaching the bottom, top or neither region; time to reach the low-salt zone and time spent in the zone. (**G**) Locomotor parameters before reaching either top or bottom of the arena. Locomotor parameters when moving with vs against the gradient. (**H**) Locomotor parameters after successful navigation and when moving with vs against the gradient.

**Fig S16.**
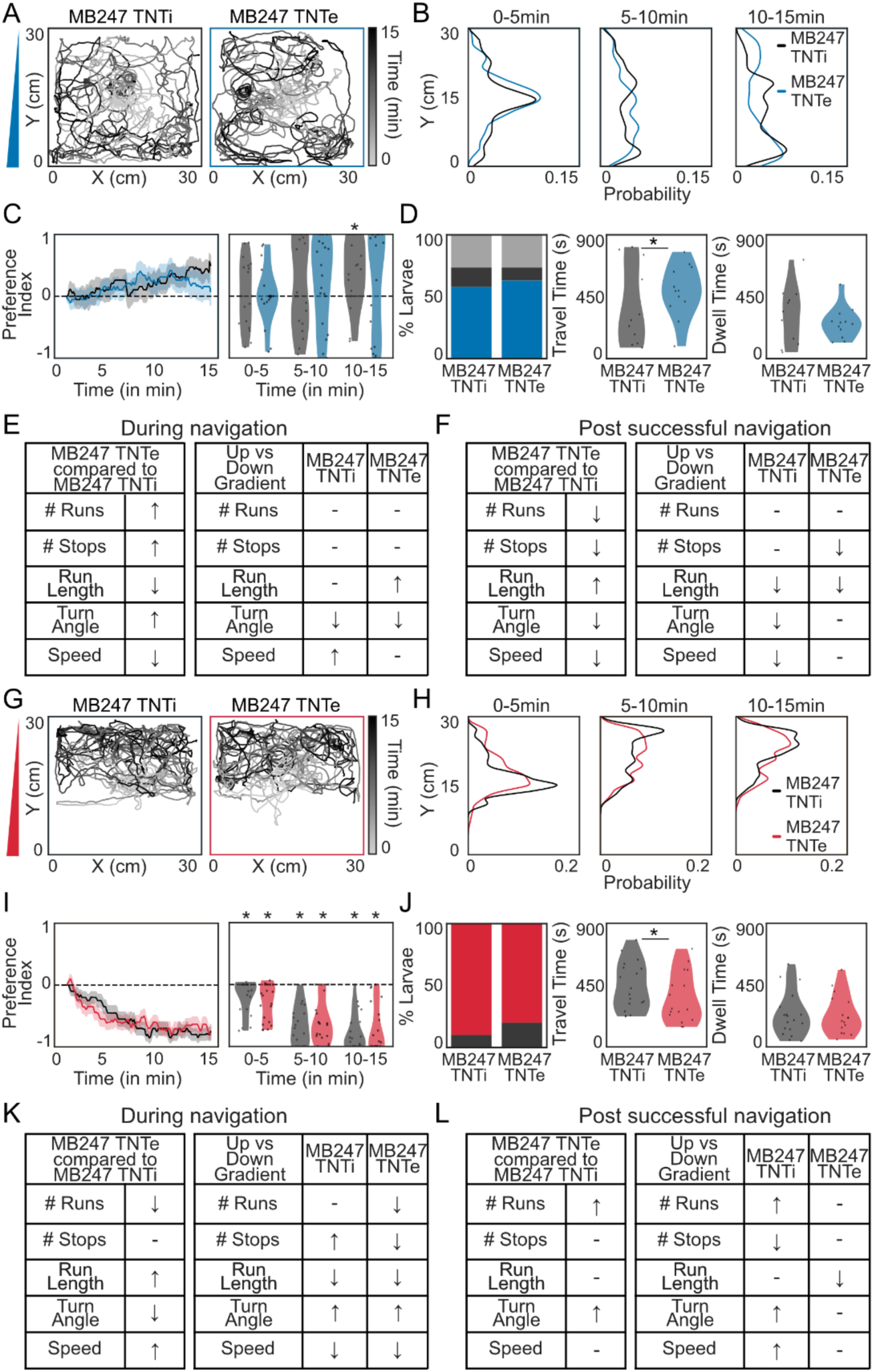
Kenyon cells are required for taste gradient navigation: (**A**) Trajectories of MB247/TNTi (black) and MB247/TNTe (blue) during fructose gradient navigation (N = 20). (**B**) Larval distribution along the Y-axis. (**C**) Preference index over time (solid line: mean; shaded: SEM) and binned preference index (5min intervals, dots: individual larvae). (**D**) Percentage of larvae either reaching the bottom, top or neither region; time to reach the high-fructose zone and time spent in the zone (Bootstrapped CI test = Cohen’s d > 0.35, see Data S1). (**E**) Locomotor parameters before reaching either top or bottom of the arena (↑ : increase, ↓ : decrease; – : no difference). Locomotor parameters when moving with vs against the gradient (Bootstrapped CI test = * Cohen’s d > 0.2, see Data S2). (**F**) Locomotor parameters after successful navigation and when moving with vs against the gradient. (**G**) Trajectories of MB247/TNTi (black) and MB247/TNTe (red) during salt gradient navigation (N = 20). (**H**) Larval distribution along the Y-axis. (I) Preference index over time and binned preference index. (**J**) Percentage of larvae either reaching the bottom, top or neither region; time to reach the low-salt zone and time spent in the zone. (**K**) Locomotor parameters before reaching either top or bottom of the arena. Locomotor parameters when moving with vs against the gradient. (**L**) Locomotor parameters after successful navigation and when moving with vs against the gradient.

**Fig S17.**
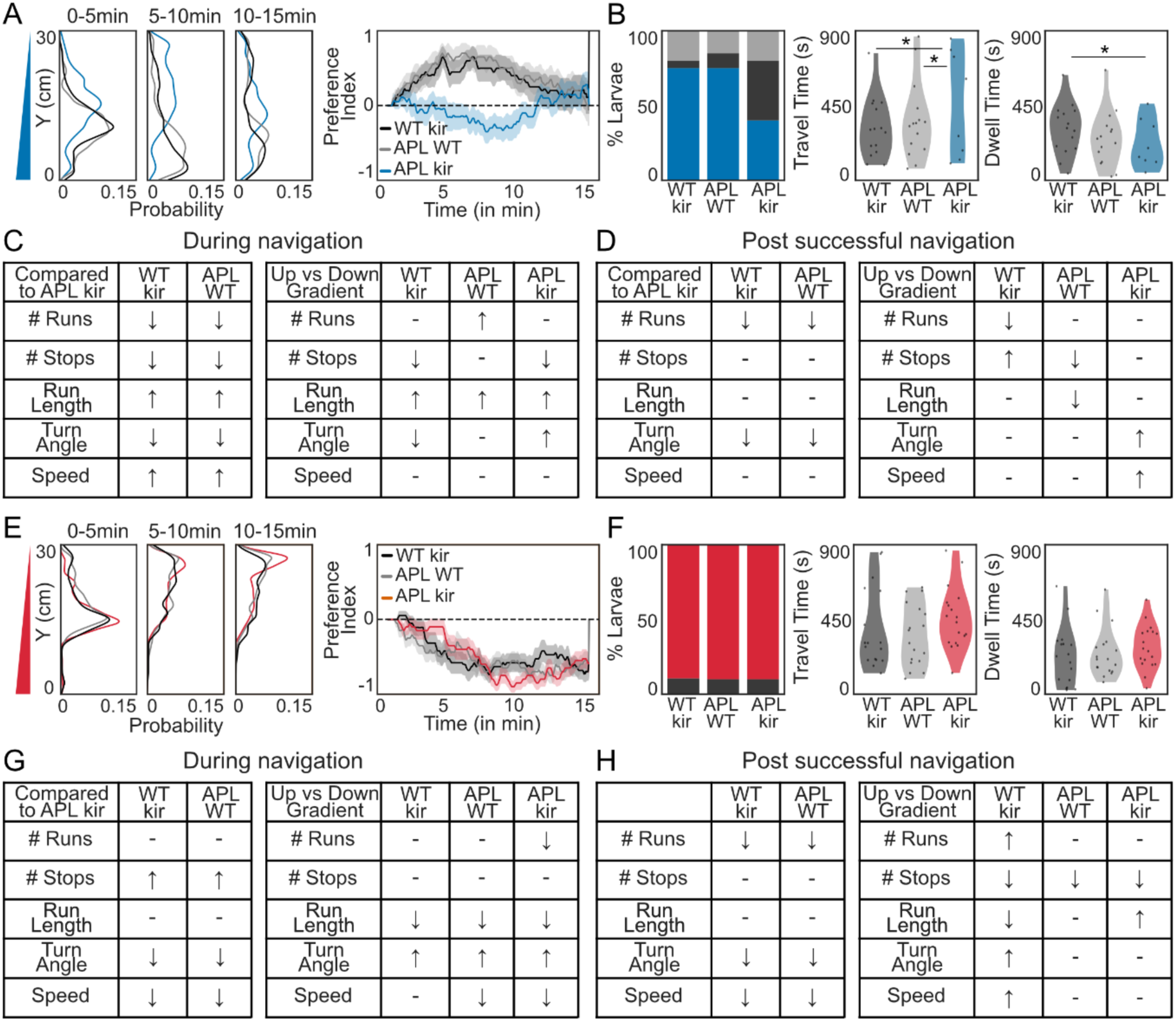
Larval behavior of WT/kir, APL/WT and APL/kir larvae during fructose and salt gradient navigation: (**A**) Distribution of WT/kir (black), APL/WT (gray) and APL/kir (blue) larvae on fructose gradient along the Y-axis. Preference index over time (solid line: mean; shaded: SEM). (**B**) Percentage of larvae either reaching the bottom, top or neither region; time to reach the high-fructose zone and time spent in the zone (Bootstrapped CI test = Cohen’s d > 0.35, see Data S1). (**C**) Locomotor parameters before reaching either top or bottom of the arena (↑ : increase, ↓ : decrease; – : no difference). Locomotor parameters when moving with vs against the gradient (Bootstrapped CI test = * Cohen’s d > 0.2, see Data S2). (**D**) Locomotor parameters after successful navigation and when moving with vs against the gradient. (**E**) Distribution of WT/kir (black), APL/WT (gray) and APL/kir (red) larvae on salt gradient along the Y-axis. Preference index over time. (**F**) Percentage of larvae either reaching the bottom, top or neither region; time to reach the low-salt zone and time spent in the zone. (**G**) Locomotor parameters before reaching either top or bottom of the arena. Locomotor parameters when moving with vs against the gradient. (**H**) Locomotor parameters after successful navigation and when moving with vs against the gradient.

**Fig S18.**
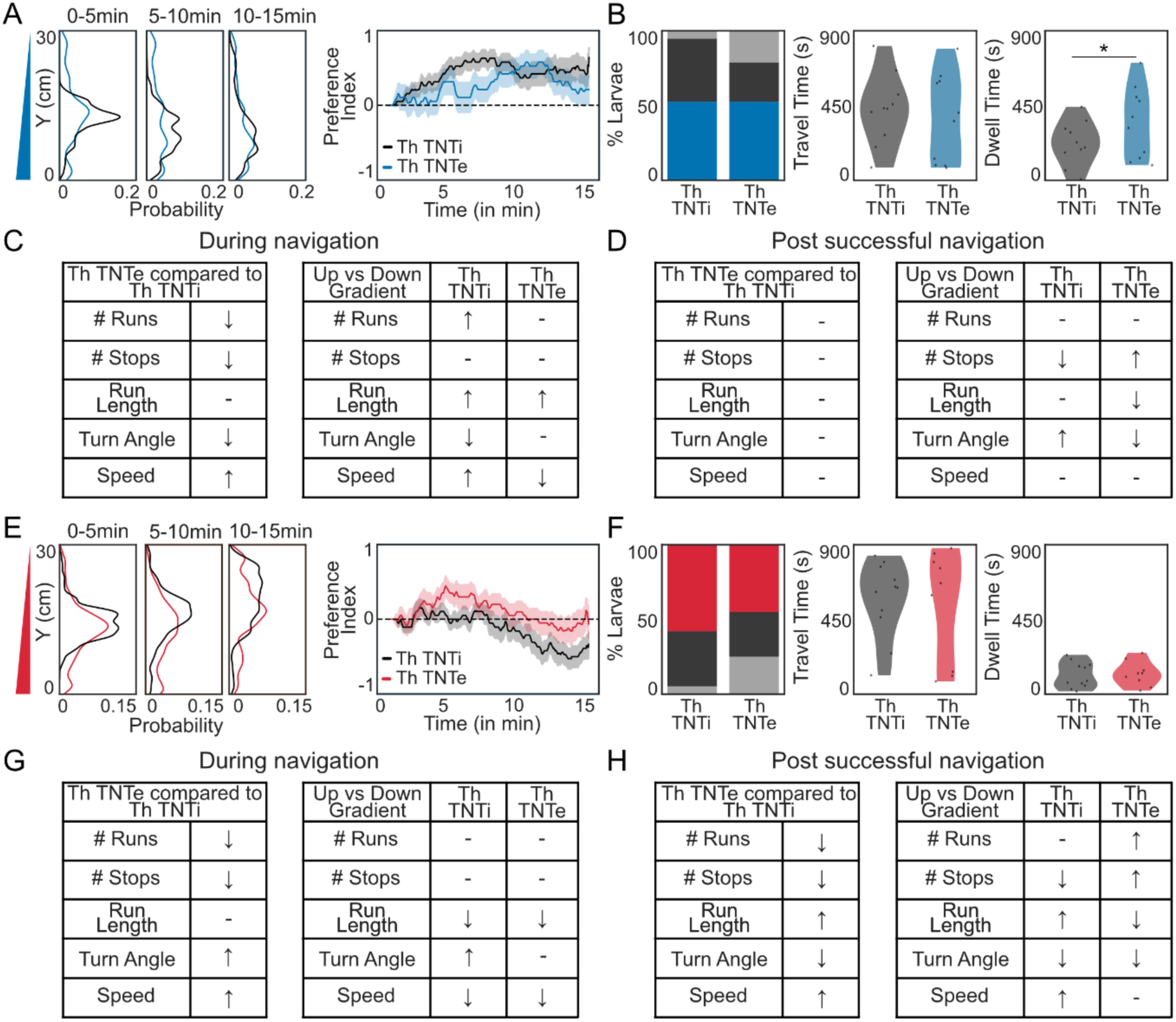
Larval behavior of Th/TNTi and Th/TNTe larvae during fructose and salt gradient navigation: (**A**) Distribution of Th/TNTi (black) and Th/TNTe (blue) larvae on fructose gradient along the Y-axis. Preference index over time (solid line: mean; shaded: SEM). (**B**) Percentage of larvae either reaching the bottom, top or neither region; time to reach the high-fructose zone and time spent in the zone (Bootstrapped CI test = Cohen’s d > 0.35, see Data S1). (**C**) Locomotor parameters before reaching either top or bottom of the arena (↑ : increase, ↓ : decrease; – : no difference). Locomotor parameters when moving with vs against the gradient (Bootstrapped CI test = * Cohen’s d > 0.2, see Data S2). (**D**) Locomotor parameters after successful navigation and when moving with vs against the gradient. (**E**) Distribution of Th/TNTi (black) and Th/TNTe (red) larvae on salt gradient along the Y-axis. Preference index over time. (**F**) Percentage of larvae either reaching the bottom, top or neither region; time to reach the low-salt zone and time spent in the zone. (**G**) Locomotor parameters before reaching either top or bottom of the arena. Locomotor parameters when moving with vs against the gradient. (**H**) Locomotor parameters after successful navigation and when moving with vs against the gradient.

**Fig S19.**
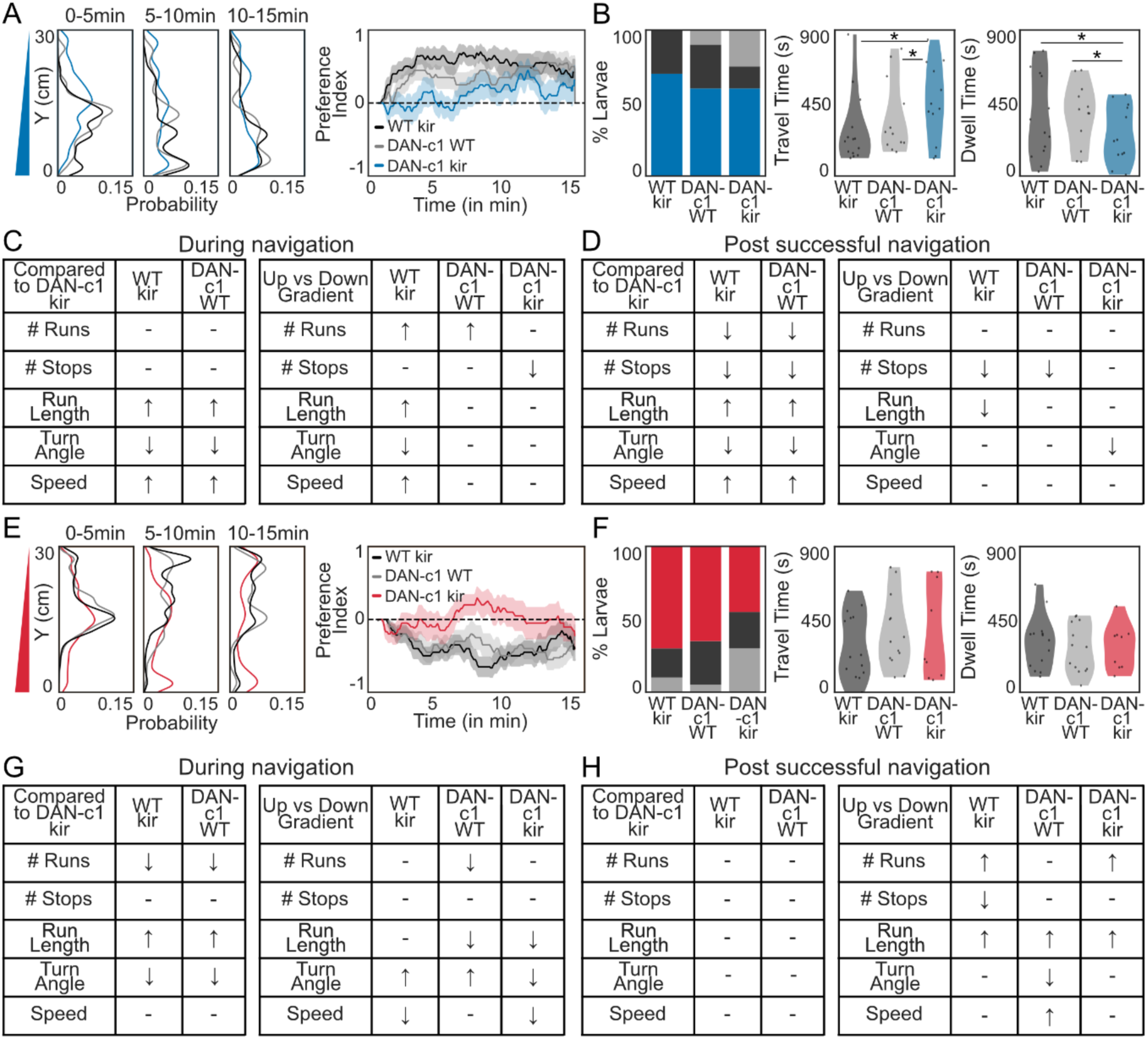
Larval behavior of WT/kir, DAN-c1/WT and DAN-c1/kir larvae during fructose and salt gradient navigation: (**A**) Distribution of WT/kir (black), DAN-c1/WT (gray) and DAN-c1/kir (blue) larvae on fructose gradient along the Y-axis. Preference index over time (solid line: mean; shaded: SEM). (**B**) Percentage of larvae either reaching the bottom, top or neither region; time to reach the high-fructose zone and time spent in the zone (Bootstrapped CI test = Cohen’s d > 0.35, see Data S1). (**C**) Locomotor parameters before reaching either top or bottom of the arena (↑ : increase, ↓ : decrease; – : no difference). Locomotor parameters when moving with vs against the gradient (Bootstrapped CI test = * Cohen’s d > 0.2, see Data S2). (**D**) Locomotor parameters after successful navigation and when moving with vs against the gradient. (**E**) Distribution of WT/kir (black), DAN-c1/WT (gray) and DAN-c1/kir (red) larvae on salt gradient along the Y-axis. Preference index over time. (**F**) Percentage of larvae either reaching the bottom, top or neither region; time to reach the low-salt zone and time spent in the zone. (**G**) Locomotor parameters before reaching either top or bottom of the arena. Locomotor parameters when moving with vs against the gradient. (**H**) Locomotor parameters after successful navigation and when moving with vs against the gradient.

**Fig S20.**
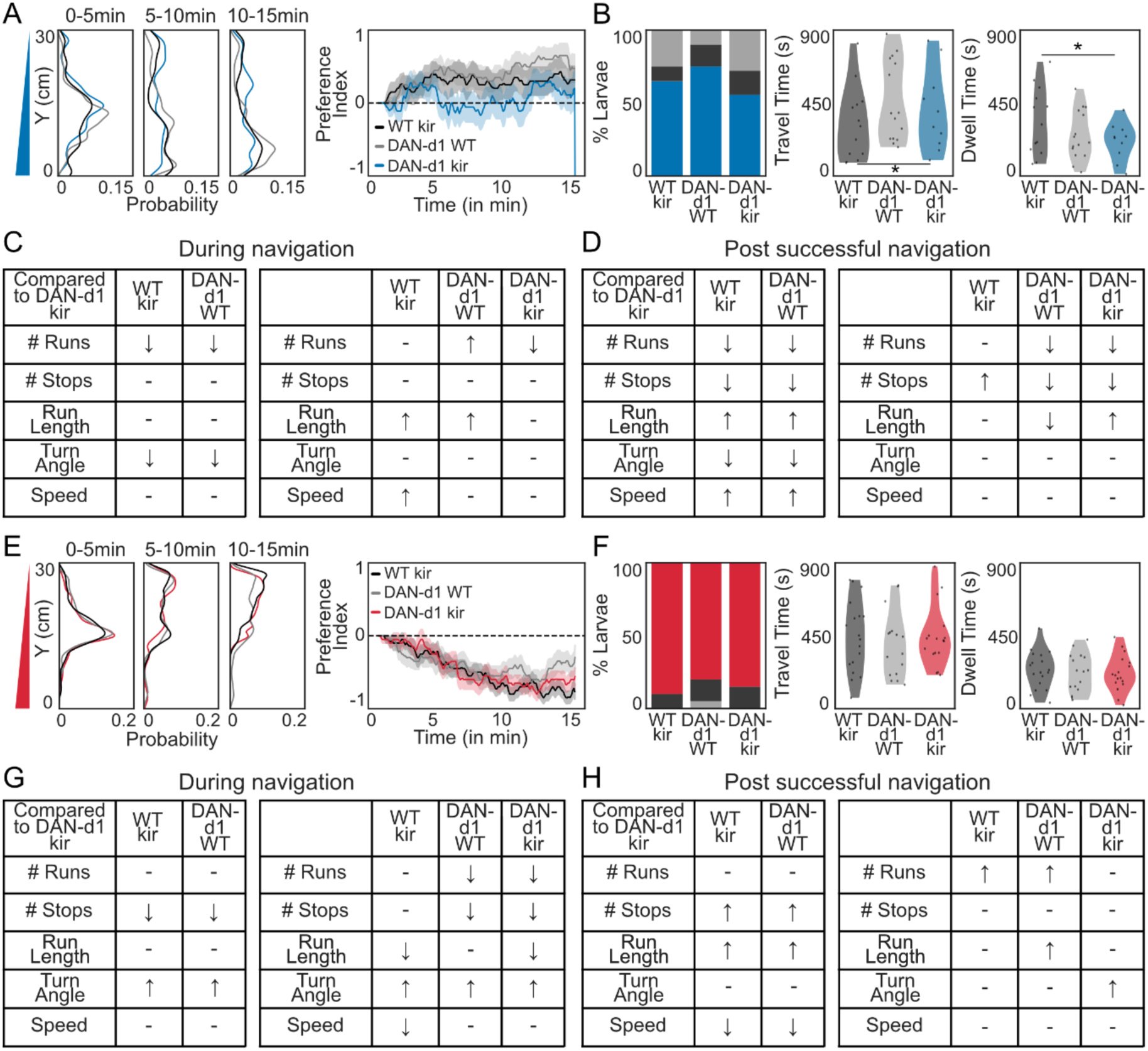
Larval behavior of WT/kir, DAN-d1/WT and DAN-d1/kir larvae during fructose and salt gradient navigation: (**A**) Distribution of WT/kir (black), DAN-d1/WT (gray) and DAN-d1/kir (blue) larvae on fructose gradient along the Y-axis. Preference index over time (solid line: mean; shaded: SEM). (**B**) Percentage of larvae either reaching the bottom, top or neither region; time to reach the high-fructose zone and time spent in the zone (Bootstrapped CI test = Cohen’s d > 0.35, see Data S1). (**C**) Locomotor parameters before reaching either top or bottom of the arena (↑ : increase, ↓ : decrease; – : no difference). Locomotor parameters when moving with vs against the gradient (Bootstrapped CI test = * Cohen’s d > 0.2, see Data S2). (**D**) Locomotor parameters after successful navigation and when moving with vs against the gradient. (**E**) Distribution of WT/kir (black), DAN-d1/WT (gray) and DAN-d1/kir (red) larvae on salt gradient along the Y-axis. Preference index over time. (**F**) Percentage of larvae either reaching the bottom, top or neither region; time to reach the low-salt zone and time spent in the zone. (**G**) Locomotor parameters before reaching either top or bottom of the arena. Locomotor parameters when moving with vs against the gradient. (**H**) Locomotor parameters after successful navigation and when moving with vs against the gradient.

**Fig S21.**
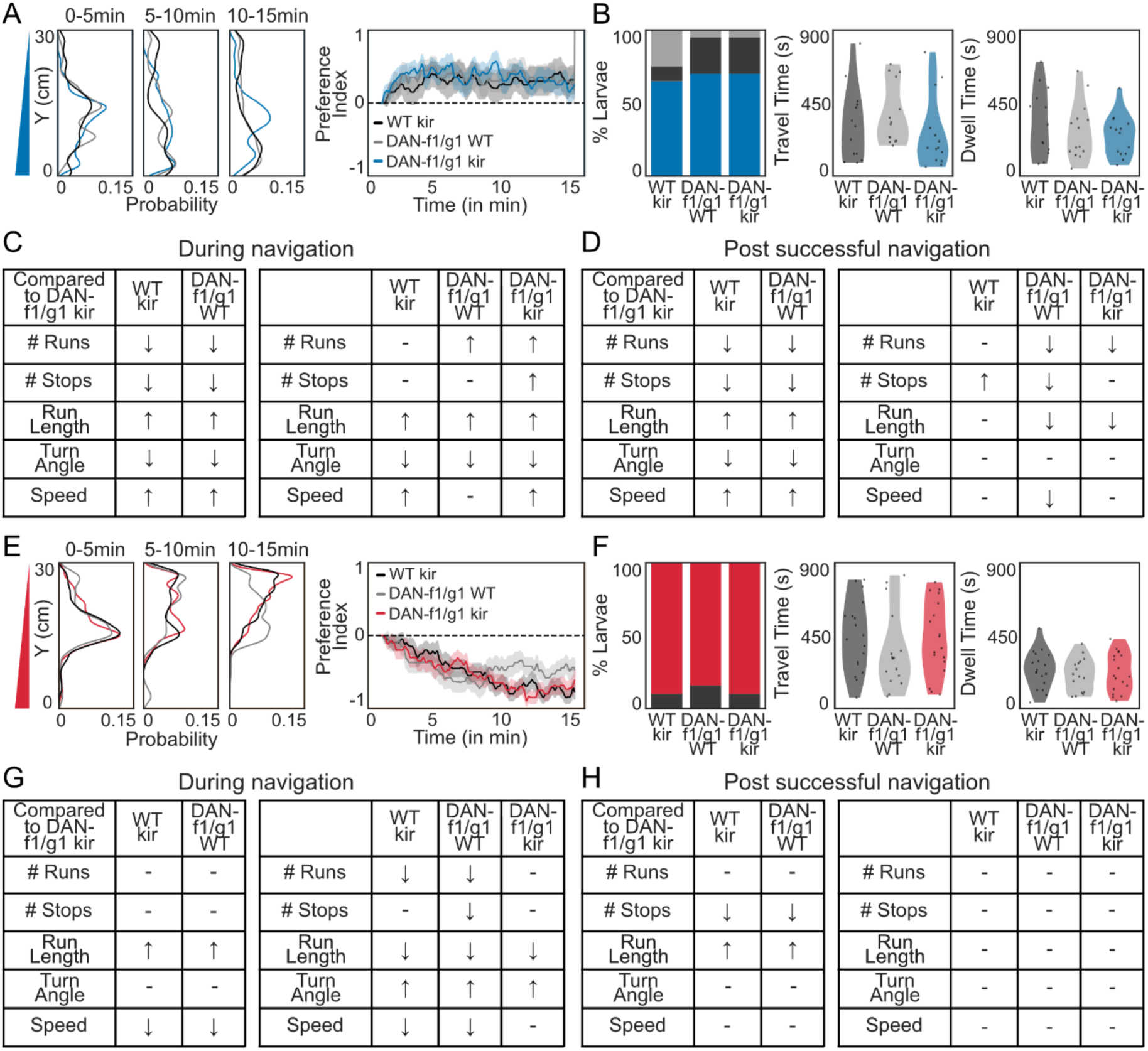
Larval behavior of WT kir, DAN-f1/g1 WT and DAN-f1/g1 kir larvae during fructose and salt gradient navigation: (**A**) Distribution of WT/kir (black), DAN-f1/g1/WT (gray) and DAN-f1/g1/kir (blue) larvae on fructose gradient along the Y-axis. Preference index over time (solid line: mean; shaded: SEM). (**B**) Percentage of larvae either reaching the bottom, top or neither region; time to reach the high-fructose zone and time spent in the zone (Bootstrapped CI test = Cohen’s d > 0.35, see Data S1). (**C**) Locomotor parameters before reaching either top or bottom of the arena (↑ : increase, ↓ : decrease; – : no difference). Locomotor parameters when moving with vs against the gradient (Bootstrapped CI test = * Cohen’s d > 0.2, see Data S2). (**D**) Locomotor parameters after successful navigation and when moving with vs against the gradient. (**E**) Distribution of WT/kir (black), DAN-f1/g1/WT (gray) and DAN-f1/g1/kir (red) larvae on salt gradient along the Y-axis. Preference index over time. (**F**) Percentage of larvae either reaching the bottom, top or neither region; time to reach the low-salt zone and time spent in the zone. (**G**) Locomotor parameters before reaching either top or bottom of the arena. Locomotor parameters when moving with vs against the gradient. (**H**) Locomotor parameters after successful navigation and when moving with vs against the gradient.

**Fig S22.**
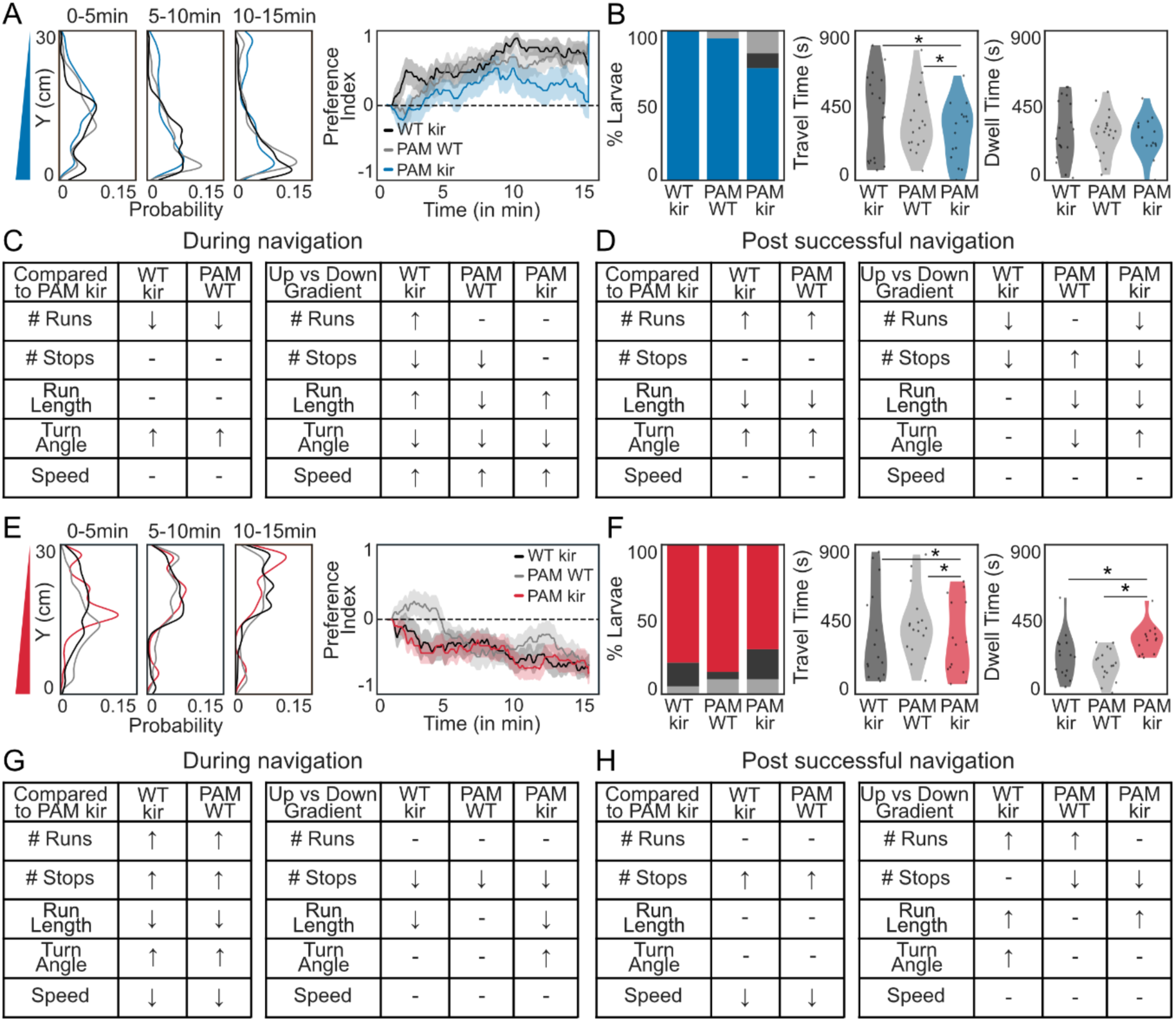
Larval behavior of WT kir, PAM WT and PAM kir larvae during fructose and salt gradient navigation: (**A**) Distribution of WT/kir (black), PAM/WT (gray) and PAM/kir (blue) larvae on fructose gradient along the Y-axis. Preference index over time (solid line: mean; shaded: SEM). (**B**) Percentage of larvae either reaching the bottom, top or neither region; time to reach the high-fructose zone and time spent in the zone (Bootstrapped CI test = Cohen’s d > 0.35, see Data S1). (**C**) Locomotor parameters before reaching either top or bottom of the arena (↑ : increase, ↓ : decrease; – : no difference). Locomotor parameters when moving with vs against the gradient (Bootstrapped CI test = * Cohen’s d > 0.2, see Data S2). (**D**) Locomotor parameters after successful navigation and when moving with vs against the gradient. (**E**) Distribution of WT/kir (black), PAM/WT (gray) and PAM/kir (red) larvae on salt gradient along the Y-axis. Preference index over time. (**F**) Percentage of larvae either reaching the bottom, top or neither region; time to reach the low-salt zone and time spent in the zone. (**G**) Locomotor parameters before reaching either top or bottom of the arena. Locomotor parameters when moving with vs against the gradient. (**H**) Locomotor parameters after successful navigation and when moving with vs against the gradient.

**Fig S23.**
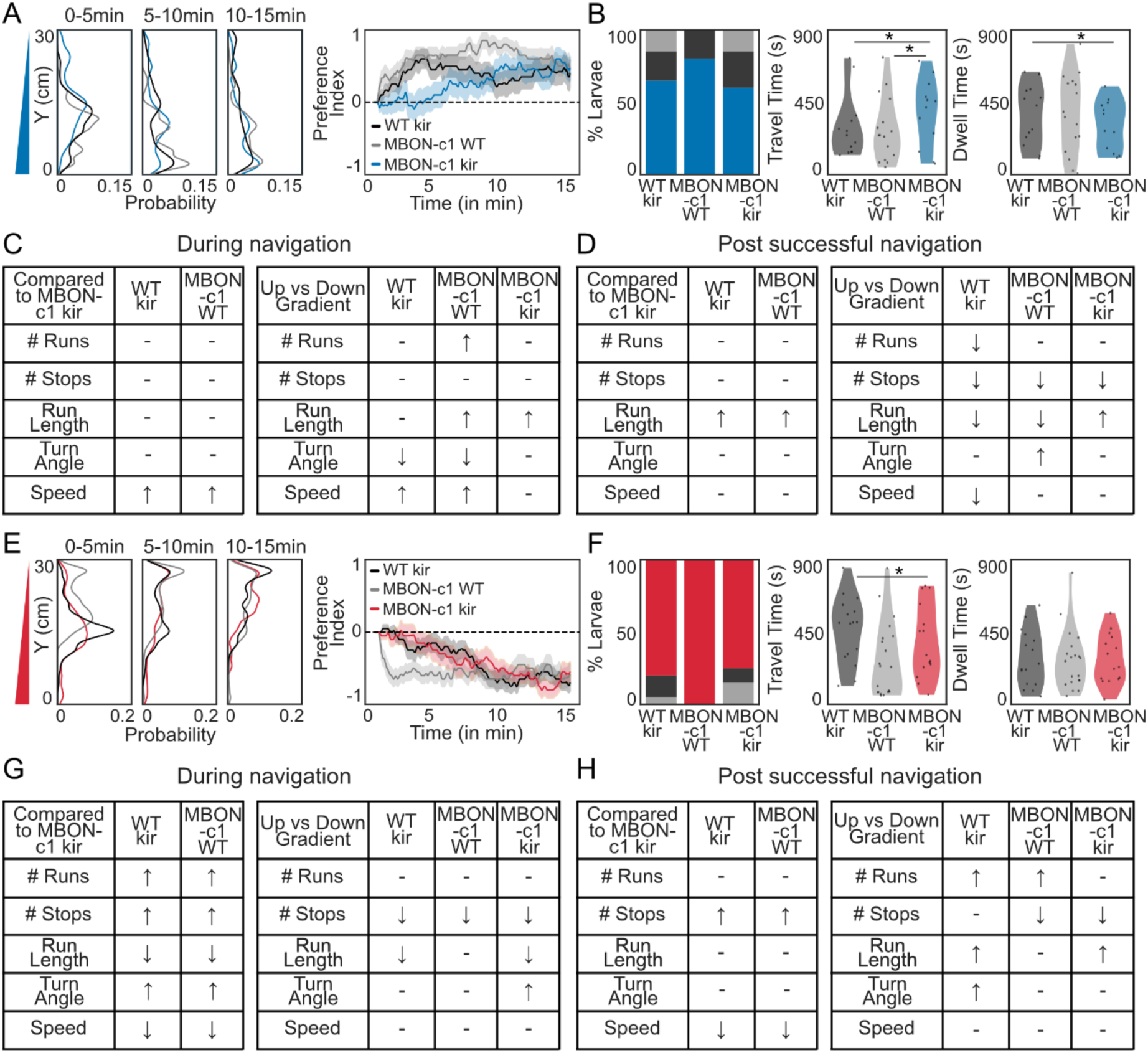
Larval behavior of WT kir, MBON-c1 WT and MBON-c1 kir larvae during fructose and salt gradient navigation: (**A**) Distribution of WT/kir (black), MBON-c1/WT (gray) and MBON-c1/kir (blue) larvae on fructose gradient along the Y-axis. Preference index over time (solid line: mean; shaded: SEM). (**B**) Percentage of larvae either reaching the bottom, top or neither region; time to reach the high-fructose zone and time spent in the zone. (Bootstrapped CI test = Cohen’s d > 0.35, see Data S1). (**C**) Locomotor parameters before reaching either top or bottom of the arena (↑ : increase, ↓ : decrease; – : no difference). Locomotor parameters when moving with vs against the gradient (Bootstrapped CI test = * Cohen’s d > 0.2, see Data S2). (**D**) Locomotor parameters after successful navigation and when moving with vs against the gradient. (**E**) Distribution of WT/kir (black), MBON-c1/WT (gray) and MBON-c1/kir (red) larvae on salt gradient along the Y-axis. Preference index over time. (**F**) Percentage of larvae either reaching the bottom, top or neither region; time to reach the low-salt zone and time spent in the zone. (**G**) Locomotor parameters before reaching either top or bottom of the arena. Locomotor parameters when moving with vs against the gradient. (**H**) Locomotor parameters after successful navigation and when moving with vs against the gradient.

**Fig S24.**
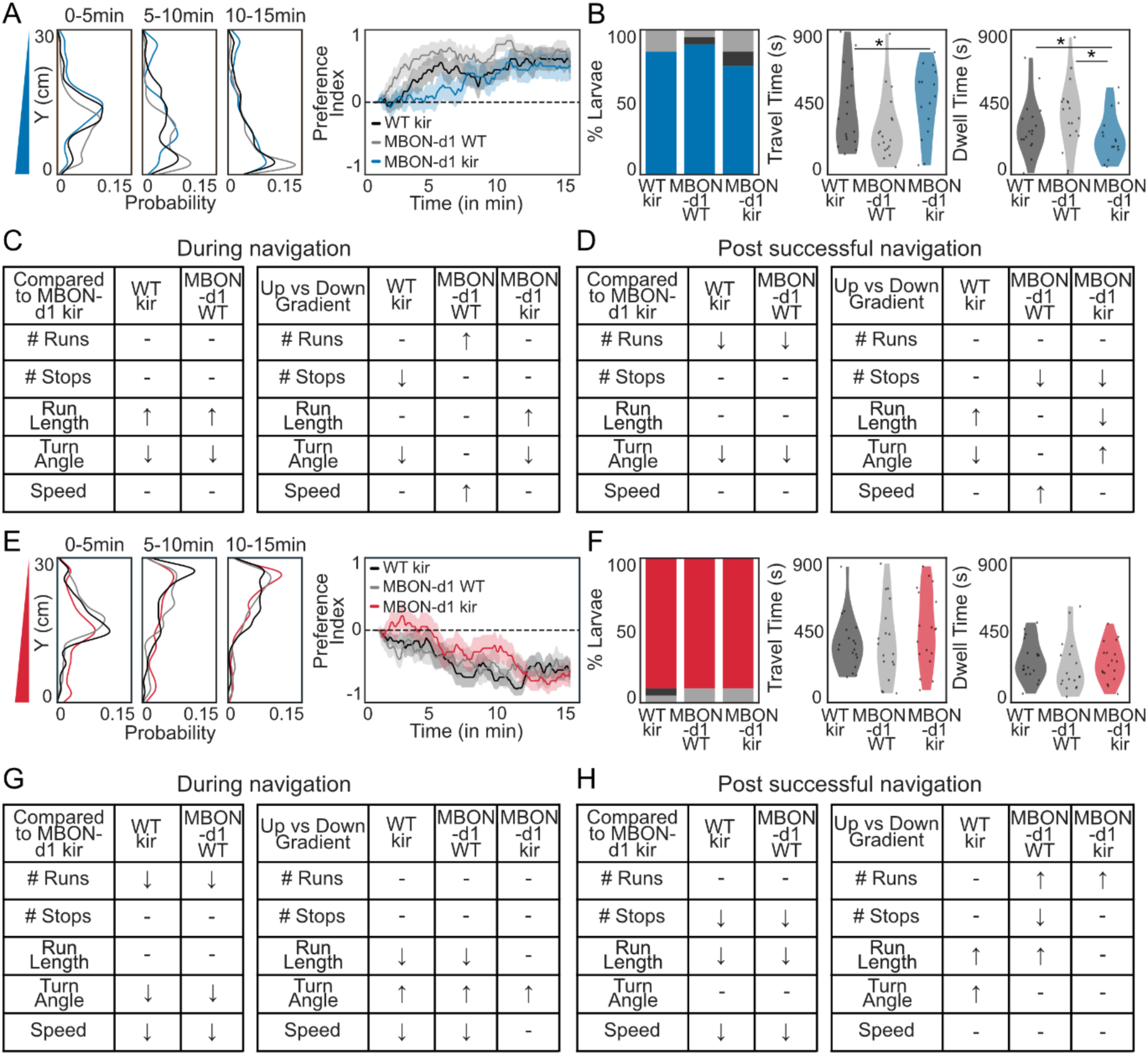
Larval behavior of WT kir, MBON-d1 WT and MBON-d1 kir larvae during fructose and salt gradient navigation: (**A**) Distribution of WT/kir (black), MBON-d1/WT (gray) and MBON-d1/kir (blue) larvae on fructose gradient along the Y-axis. Preference index over time (solid line: mean; shaded: SEM). (**B**) Percentage of larvae either reaching the bottom, top or neither region; time to reach the high-fructose zone and time spent in the zone. (Bootstrapped CI test = Cohen’s d > 0.35, see Data S1). (**C**) Locomotor parameters before reaching either top or bottom of the arena (↑ : increase, ↓ : decrease; – : no difference). Locomotor parameters when moving with vs against the gradient (Bootstrapped CI test = * Cohen’s d > 0.2, see Data S2). (**D**) Locomotor parameters after successful navigation and when moving with vs against the gradient. (**E**) Distribution of WT/kir (black), MBON-d1/WT (gray) and MBON-d1/kir (red) larvae on salt gradient along the Y-axis. Preference index over time. (**F**) Percentage of larvae either reaching the bottom, top or neither region; time to reach the low-salt zone and time spent in the zone. (**G**) Locomotor parameters before reaching either top or bottom of the arena. Locomotor parameters when moving with vs against the gradient. (**H**) Locomotor parameters after successful navigation and when moving with vs against the gradient.

**Fig S25.**
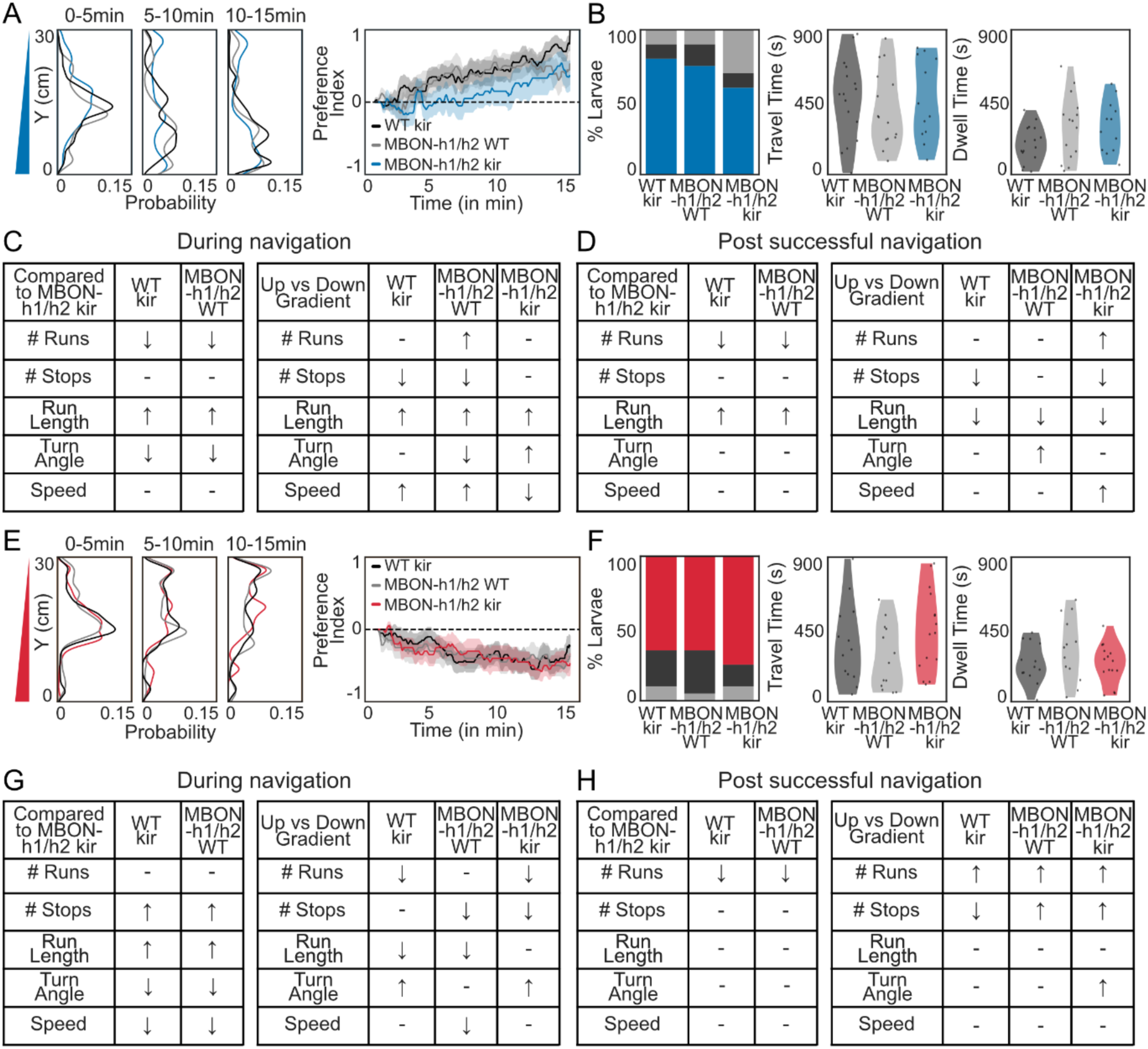
Larval behavior of WT kir, MBON-h1/h2 WT and MBON-h1/h2 kir larvae during fructose and salt gradient navigation: (**A**) Distribution of WT/kir (black), MBON-h1/h2/WT (gray) and MBON-h1/h2/kir (blue) larvae on fructose gradient along the Y-axis. Preference index over time (solid line: mean; shaded: SEM). (**B**) Percentage of larvae either reaching the bottom, top or neither region; time to reach the high-fructose zone and time spent in the zone (Bootstrapped CI test = Cohen’s d > 0.35, see Data S1). (**C**) Locomotor parameters before reaching either top or bottom of the arena (↑ : increase, ↓ : decrease; – : no difference). Locomotor parameters when moving with vs against the gradient (Bootstrapped CI test = * Cohen’s d > 0.2, see Data S2). (**D**) Locomotor parameters after successful navigation and when moving with vs against the gradient. (**E**) Distribution of WT/kir (black), MBON-h1/h2/WT (gray) and MBON-h1/h2/kir (red) larvae on salt gradient along the Y-axis. Preference index over time. (**F**) Percentage of larvae either reaching the bottom, top or neither region; time to reach the low-salt zone and time spent in the zone. (**G**) Locomotor parameters before reaching either top or bottom of the arena. Locomotor parameters when moving with vs against the gradient. (**H**) Locomotor parameters after successful navigation and when moving with vs against the gradient.

**Fig S26.**
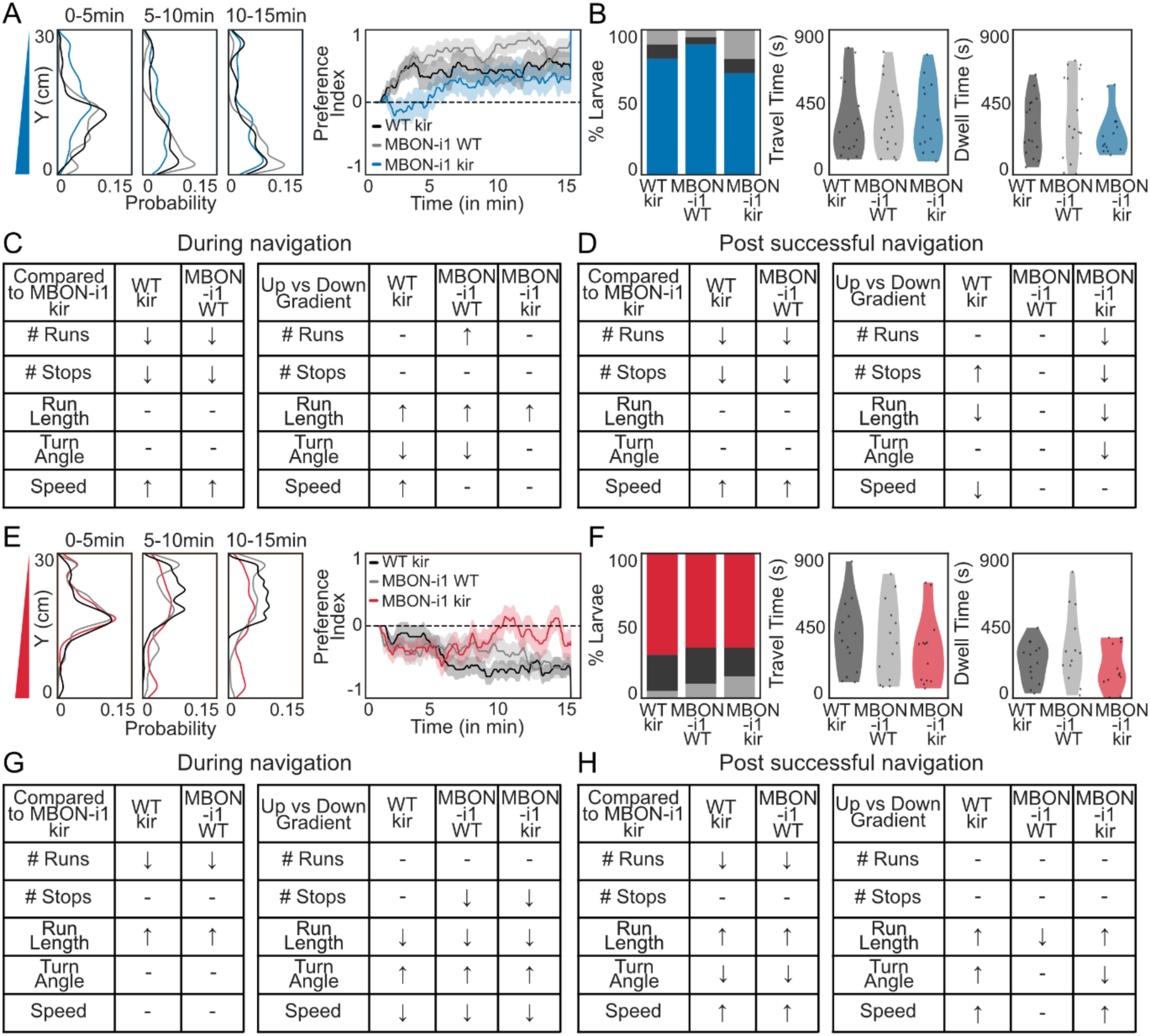
Larval behavior of WT/kir, MBON-i1/WT and MBON-i1/kir larvae during fructose and salt gradient navigation: (**A**) Distribution of WT/kir (black), MBON-i1/WT (gray) and MBON-i1/kir (blue) larvae on fructose gradient along the Y-axis. Preference index over time (solid line: mean; shaded: SEM). (**B**) Percentage of larvae either reaching the bottom, top or neither region; time to reach the high-fructose zone and time spent in the zone. (Bootstrapped CI test = Cohen’s d > 0.35, see Data S1). (**C**) Locomotor parameters before reaching either top or bottom of the arena (↑ : increase, ↓ : decrease; – : no difference). Locomotor parameters when moving with vs against the gradient (Bootstrapped CI test = * Cohen’s d > 0.2, see Data S2). (**D**) Locomotor parameters after successful navigation and when moving with vs against the gradient. (**E**) Distribution of WT/kir (black), MBON-i1/WT (gray) and MBON-i1/kir (red) larvae on salt gradient along the Y-axis. Preference index over time. (**F**) Percentage of larvae either reaching the bottom, top or neither region; time to reach the low-salt zone and time spent in the zone. (**G**) Locomotor parameters before reaching either top or bottom of the arena. Locomotor parameters when moving with vs against the gradient. (**H**) Locomotor parameters after successful navigation and when moving with vs against the gradient.

**Fig S27.**
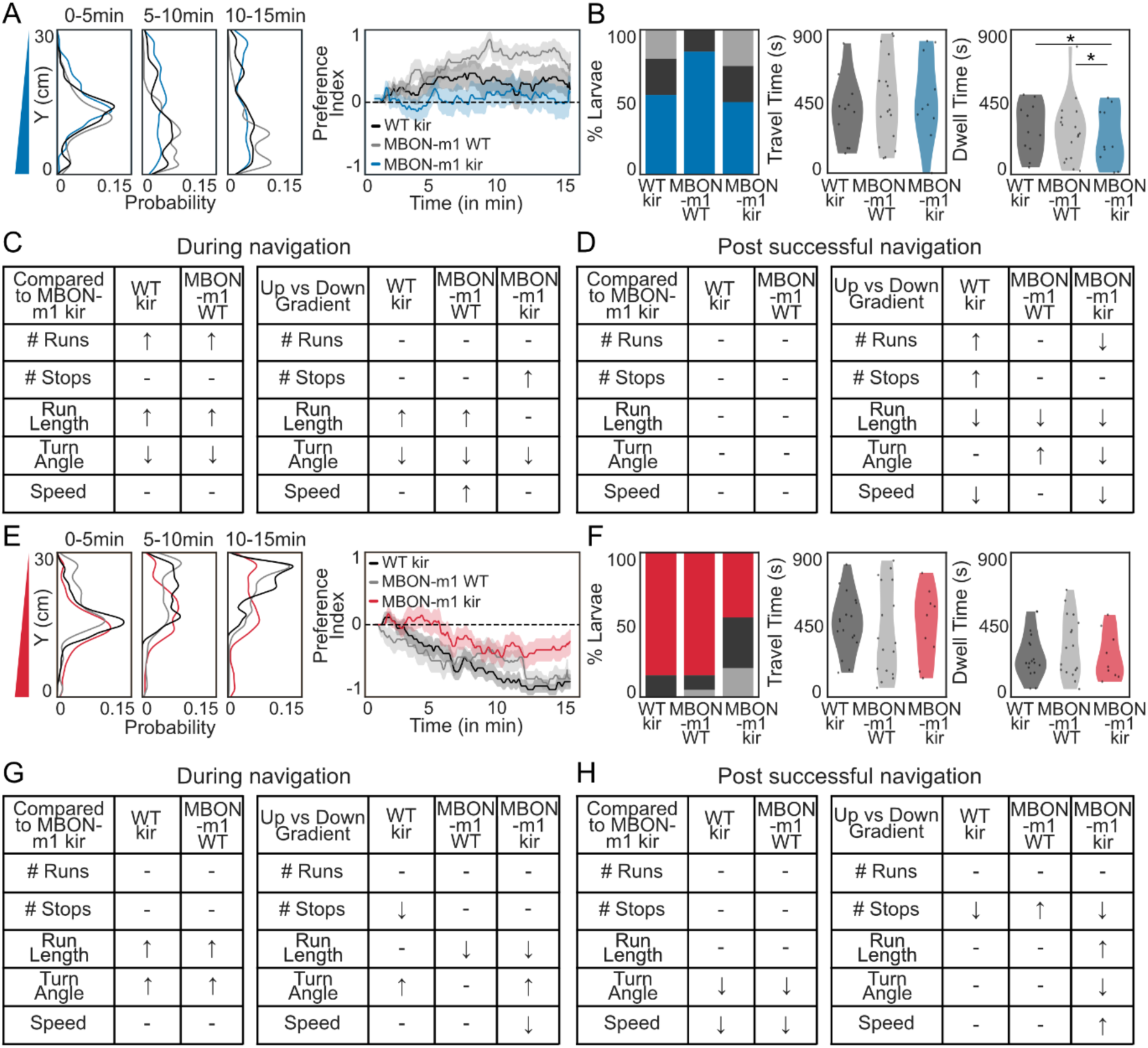
Larval behavior of WT/jukir, MBON-m1/WT and MBON-m1/kir larvae during fructose and salt gradient navigation: (**A**) Distribution of WT/kir (black), MBON-m1/WT (gray) and MBON-m1/kir (blue) larvae on fructose gradient along the Y-axis. Preference index over time (solid line: mean; shaded: SEM). (**B**) Percentage of larvae either reaching the bottom, top or neither region; time to reach the high-fructose zone and time spent in the zone (Bootstrapped CI test = Cohen’s d > 0.35, see Data S1). (**C**) Locomotor parameters before reaching either top or bottom of the arena (↑ : increase, ↓ : decrease; – : no difference). Locomotor parameters when moving with vs against the gradient. (**D**) Locomotor parameters after successful navigation and when moving with vs against the gradient (Bootstrapped CI test = * Cohen’s d > 0.2, see Data S2). (**E**) Distribution of WT/kir (black), MBON-m1/WT (gray) and MBON-m1/kir (red) larvae on salt gradient along the Y-axis. Preference index over time. (**F**) Percentage of larvae either reaching the bottom, top or neither region; time to reach the low-salt zone and time spent in the zone. (**G**) Locomotor parameters before reaching either top or bottom of the arena. Locomotor parameters when moving with vs against the gradient. (**H**) Locomotor parameters after successful navigation and when moving with vs against the gradient.

## Supplementary Videos

**Movie S1:** Superimposed trajectories of individual larvae navigating fructose gradient for 15 minutes (N = 20, shown at 10x).

**Movie S2:** Superimposed trajectories of individual larvae navigating salt gradient for 15 minutes (N = 20, shown at 10x).

**Movie S3:** Superimposed trajectories of individual larvae on agar for 15 minutes (N = 20, shown at 10x).

## Supplementary Files

**Data S1:** Statistical results for data in the main and supplement figures.

**Data S2:** Parameter values (Average +/- STD) and statistical results for all locomotion analysis.

## References

Alves, G., Salle, J., Chaudy, S., Dupas, S., & Maniere, G. (2014). High-NaCl Perception in Drosophila melanogaster. Journal of Neuroscience, 34(33), 10884–10891. 10.1523/JNEUROSCI.4795-13.2014

Apostolopoulou, A. A., Rist, A., & Thum, A. S. (2015). Taste processing in Drosophila larvae. Frontiers in Integrative Neuroscience, 9. 10.3389/fnint.2015.00050

Aso, Y., Sitaraman, D., Ichinose, T., Kaun, K. R., Vogt, K., Belliart-Guérin, G., Plaçais, P.-Y., Robie, A. A., Yamagata, N., Schnaitmann, C., Rowell, W. J., Johnston, R. M., Ngo, T.-T. B., Chen, N., Korff, W., Nitabach, M. N., Heberlein, U., Preat, T., Branson, K. M., … Rubin, G. M. (2014). Mushroom body output neurons encode valence and guide memory-based action selection in Drosophila. eLife, 3, e04580. 10.7554/eLife.04580

Bakker, K. (1959). Feeding period, growth, and pupation in larvae of *Drosophila melanogaster*. Entomologia Experimentalis et Applicata, 2(3), 171–186. 10.1111/j.1570-7458.1959.tb00432.x

Balakrishnan, K. A., & Haesemeyer, M. (2025). Behavioral and circuit principles of temperature gradient navigation. Current Biology, 35(22), 5395–5410.e8. 10.1016/j.cub.2025.09.053

Berck, M. E., Khandelwal, A., Claus, L., Hernandez-Nunez, L., Si, G., Tabone, C. J., Li, F., Truman, J. W., Fetter, R. D., Louis, M., Samuel, A. D., & Cardona, A. (2016). The wiring diagram of a glomerular olfactory system. eLife, 5, e14859. 10.7554/eLife.14859

Chen, X., & Engert, F. (2014). Navigational strategies underlying phototaxis in larval zebrafish. Frontiers in Systems Neuroscience, 8. 10.3389/fnsys.2014.00039

Dancausse, S., Robles, J., Fitzpatrick, J., Garcia, C., Fernando, O., Evans, A., Baker, J. D., & Klein, M. (2026). Nicotine-driven hyperactivation of larval locomotion. 10.7554/eLife.110613.1

Eichler, K., Li, F., Litwin-Kumar, A., Park, Y., Andrade, I., Schneider-Mizell, C. M., Saumweber, T., Huser, A., Eschbach, C., Gerber, B., Fetter, R. D., Truman, J. W., Priebe, C. E., Abbott, L. F., Thum, A. S., Zlatic, M., & Cardona, A. (2017). The complete connectome of a learning and memory centre in an insect brain. Nature, 548(7666), 175–182. 10.1038/nature23455

Eschbach, C., Fushiki, A., Winding, M., Afonso, B., Andrade, I. V., Cocanougher, B. T., Eichler, K., Gepner, R., Si, G., Valdes-Aleman, J., Fetter, R. D., Gershow, M., Jefferis, G. S., Samuel, A. D., Truman, J. W., Cardona, A., & Zlatic, M. (2021). Circuits for integrating learned and innate valences in the insect brain. eLife, 10, e62567. 10.7554/eLife.62567

Eschbach, C., Fushiki, A., Winding, M., Schneider-Mizell, C. M., Shao, M., Arruda, R., Eichler, K., Valdes-Aleman, J., Ohyama, T., Thum, A. S., Gerber, B., Fetter, R. D., Truman, J. W., Litwin-Kumar, A., Cardona, A., & Zlatic, M. (2020). Recurrent architecture for adaptive regulation of learning in the insect brain. Nature Neuroscience, 23(4), 544–555. 10.1038/s41593-020-0607-9

Eschbach, C., Vogt, K., Afonso, B., Polizos, N., Dancausse, S., Evans, A., Verbe, A., Wang, K., Berck, M., Samuel, A., Klein, M., Cardona, A., & Zlatic, M. (2025). A multisensory, bidirectional, valence encoder guides behavioral decisions. Neuroscience. 10.1101/2025.09.26.678749

Franke, U. S., Großjohann, A., Aurich, S., Köhler, I., Lamberty, M., Granato, S., Fallaha, A., Breitfeld, J., Kovacs, P., Selcho, M., Kittel, R. J., Thum, A. S., & Pauls, D. (2026). Selective octopaminergic tuning of mushroom body circuits during memory formation. Proceedings of the National Academy of Sciences, 123(16), e2517403123. 10.1073/pnas.2517403123

Gomez-Marin, A., & Louis, M. (2012). Active sensation during orientation behavior in the Drosophila larva: More sense than luck. Current Opinion in Neurobiology, 22(2), 208–215. 10.1016/j.conb.2011.11.008

Gomez-Marin, A., Stephens, G. J., & Louis, M. (2011). Active sampling and decision making in Drosophila chemotaxis. Nature Communications, 2(1), 441. 10.1038/ncomms1455

Kane, E. A., Gershow, M., Afonso, B., Larderet, I., Klein, M., Carter, A. R., De Bivort, B. L., Sprecher, S. G., & Samuel, A. D. T. (2013). Sensorimotor structure of *Drosophila* larva phototaxis. Proceedings of the National Academy of Sciences, 110(40). 10.1073/pnas.1215295110

Kathman, N. D., Lanz, A. J., Freed, J. D., & Nagel, K. I. (2024). *Neural dynamics for working memory and evidence integration during olfactory navigation in* Drosophila. Neuroscience. 10.1101/2024.10.05.616803

King, R. C. (1953). Effects of Alkali Metal Ions on Development of *Drosophila*, with Special Reference to Lithium-Induced Abnormalities. Proceedings of the National Academy of Sciences, 39(5), 403–407. 10.1073/pnas.39.5.403

Klein, M., Afonso, B., Vonner, A. J., Hernandez-Nunez, L., Berck, M., Tabone, C. J., Kane, E. A., Pieribone, V. A., Nitabach, M. N., Cardona, A., Zlatic, M., Sprecher, S. G., Gershow, M., Garrity, P. A., & Samuel, A. D. T. (2015). Sensory determinants of behavioral dynamics in *Drosophila* thermotaxis. Proceedings of the National Academy of Sciences, 112(2). 10.1073/pnas.1416212112

Komarov, N., Fritsch, C., Maier, G. L., Bues, J., Biočanin, M., Avalos, C. B., Dodero, A., Kwon, J. Y., Deplancke, B., & Sprecher, S. G. (2025). Food hardness preference reveals multisensory contributions of fly larval gustatory organs in behaviour and physiology. PLOS Biology, 23(1), e3002730. 10.1371/journal.pbio.3002730

Kramer, T. S., Wan, F. K., Pugliese, S. M., Atanas, A. A., Pradhan, S., Hiser, A. W., Godinez, L. M., Luo, J., Bueno, E., Felt, T., & Flavell, S. W. (2026). Neural sequences underlying directed turning in Caenorhabditis elegans. Nature Neuroscience. 10.1038/s41593-026-02257-5

Liu, L., Leonard, A. S., Motto, D. G., Feller, M. A., Price, M. P., Johnson, W. A., & Welsh, M. J. (2003). Contribution of Drosophila DEG/ENaC Genes to Salt Taste. Neuron, 39(1), 133–146. 10.1016/S0896-6273(03)00394-5

Louis, M., Huber, T., Benton, R., Sakmar, T. P., & Vosshall, L. B. (2008). Bilateral olfactory sensory input enhances chemotaxis behavior. Nature Neuroscience, 11(2), 187–199. 10.1038/nn2031

Lungu, L., Ceffa, N., Clayton, M., Zlatic, M., & Cardona, A. (2025). The central complex of the larval fruit fly brain. Neuroscience. 10.1101/2025.09.30.679510

Luo, L., Gershow, M., Rosenzweig, M., Kang, K., Fang-Yen, C., Garrity, P. A., & Samuel, A. D. T. (2010). Navigational Decision Making in Drosophila Thermotaxis. The Journal of Neuroscience, 30(12), 4261–4272. 10.1523/JNEUROSCI.4090-09.2010

Luo, L., Wen, Q., Ren, J., Hendricks, M., Gershow, M., Qin, Y., Greenwood, J., Soucy, E. R., Klein, M., Smith-Parker, H. K., Calvo, A. C., Colón-Ramos, D. A., Samuel, A. D. T., & Zhang, Y. (2014). Dynamic Encoding of Perception, Memory, and Movement in a C. elegans Chemotaxis Circuit. Neuron, 82(5), 1115–1128. 10.1016/j.neuron.2014.05.010

Maier, G. L., Komarov, N., Meyenhofer, F., Kwon, J. Y., & Sprecher, S. G. (2021). Taste sensing and sugar detection mechanisms in Drosophila larval primary taste center. eLife, 10, e67844. 10.7554/eLife.67844

Mancini, N., Thoener, J., Tafani, E., Pauls, D., Mayseless, O., Strauch, M., Eichler, K., Champion, A., Kobler, O., Weber, D., Sen, E., Weiglein, A., Hartenstein, V., Chytoudis-Peroudis, C.-C., Jovanic, T., Thum, A. S., Rohwedder, A., Schleyer, M., & Gerber, B. (2023). Rewarding Capacity of Optogenetically Activating a Giant GABAergic Central-Brain Interneuron in Larval *Drosophila*. The Journal of Neuroscience, 43(44), 7393–7428. 10.1523/JNEUROSCI.2310-22.2023

Markow, T. A., & O’Grady, P. (2008). Reproductive ecology of *Drosophila*. Functional Ecology, 22(5), 747–759. 10.1111/j.1365-2435.2008.01457.x

Martin, J. R., Ernst, R., & Heisenberg, M. (1998). Mushroom bodies suppress locomotor activity in Drosophila melanogaster. Learning & Memory, 5(1–2), 179–191.

Martinez-Cordera, M., Sakai, T., Saitoe, M., & Ueno, K. (2025). Comparative experience shapes sucrose preference through memory in Drosophila. Molecular Brain, 18(1), 32. 10.1186/s13041-025-01202-0

Masuda-Nakagawa, L. M., Ito, K., Awasaki, T., & O’Kane, C. J. (2014). A single GABAergic neuron mediates feedback of odor-evoked signals in the mushroom body of larval Drosophila. Frontiers in Neural Circuits, 8. 10.3389/fncir.2014.00035

Miroschnikow, A., Schlegel, P., Schoofs, A., Hueckesfeld, S., Li, F., Schneider-Mizell, C. M., Fetter, R. D., Truman, J. W., Cardona, A., & Pankratz, M. J. (2018). Convergence of monosynaptic and polysynaptic sensory paths onto common motor outputs in a Drosophila feeding connectome. eLife, 7, e40247. 10.7554/eLife.40247

Mishra, D., Miyamoto, T., Rezenom, Y. H., Broussard, A., Yavuz, A., Slone, J., Russell, D. H., & Amrein, H. (2013). The Molecular Basis of Sugar Sensing in Drosophila Larvae. Current Biology, 23(15), 1466–1471. 10.1016/j.cub.2013.06.028

Miyakawa, Y., Fujishiro, N., Kijima, H., & Morita, H. (1980). Differences in feeding response to sugars between adults and larvae in Drosophila melanogaster. Journal of Insect Physiology, 26(10), 685–688. 10.1016/0022-1910(80)90041-4

Mudunuri, A., Tučková, K., El Hady, A., & Vogt, K. (2026). *Environmental statistics and sensory experience shape patch foraging strategies in* Drosophila *larvae*. Systems Biology. 10.64898/2026.03.27.714746

Mudunuri, A., Zadigue-Dubé, É., & Vogt, K. (2026). Multimodal social context modulates larval behavior in *Drosophila*. Science Advances, 12(5), eady0750. 10.1126/sciadv.ady0750

Niewalda, T., Singhal, N., Fiala, A., Saumweber, T., Wegener, S., & Gerber, B. (2008). Salt Processing in Larval Drosophila: Choice, Feeding, and Learning Shift from Appetitive to Aversive in a Concentration-Dependent Way. Chemical Senses, 33(8), 685–692. 10.1093/chemse/bjn037

Odell, S. R., Clark, D., Zito, N., Jain, R., Gong, H., Warnock, K., Carrion-Lopez, R., Maixner, C., Prieto-Godino, L., & Mathew, D. (2022). Internal state affects local neuron function in an early sensory processing center to shape olfactory behavior in Drosophila larvae. Scientific Reports, 12(1), 15767. 10.1038/s41598-022-20147-1

Padilla Perez, D. J., VandenBrooks, J. M., Sokolowski, M. B., & Angilletta Jr, M. J. (2025). Foraging actively can be advantageous in heterogeneous environments. Biology Letters, 21(7), 20250153. 10.1098/rsbl.2025.0153

Pauls, D., Selcho, M., Gendre, N., Stocker, R. F., & Thum, A. S. (2010). *Drosophila* Larvae Establish Appetitive Olfactory Memories via Mushroom Body Neurons of Embryonic Origin. The Journal of Neuroscience, 30(32), 10655–10666. 10.1523/JNEUROSCI.1281-10.2010

Qi, C., Qian, C., Steijvers, E., Colvin, R. A., & Lee, D. (2025). A pair of dopaminergic neurons DAN-c1 mediate Drosophila larval aversive olfactory learning through D2-like receptors. eLife, 13, RP100890. 10.7554/eLife.100890.3

Richter, V., Rist, A., Kislinger, G., Laumann, M., Schoofs, A., Miroschnikow, A., Pankratz, M. J., Cardona, A., & Thum, A. S. (2025). Morphology and ultrastructure of external sense organs of Drosophila larvae. eLife, 12, RP91155. 10.7554/eLife.91155.3

Riedl, J., & Louis, M. (2012). Behavioral Neuroscience: Crawling Is a No-Brainer for Fruit Fly Larvae. Current Biology, 22(20), R867–R869. 10.1016/j.cub.2012.08.018

Rist, A., & Thum, A. S. (2017). A map of sensilla and neurons in the taste system of *drosophila* larvae. Journal of Comparative Neurology, 525(18), 3865–3889. 10.1002/cne.24308

Rohwedder, A., Pfitzenmaier, J. E., Ramsperger, N., Apostolopoulou, A. A., Widmann, A., & Thum, A. S. (2012). Nutritional Value-Dependent and Nutritional Value-Independent Effects on Drosophila melanogaster Larval Behavior. Chemical Senses, 37(8), 711–721. 10.1093/chemse/bjs055

Rohwedder, A., Wenz, N. L., Stehle, B., Huser, A., Yamagata, N., Zlatic, M., Truman, J. W., Tanimoto, H., Saumweber, T., Gerber, B., & Thum, A. S. (2016). Four Individually Identified Paired Dopamine Neurons Signal Reward in Larval Drosophila. Current Biology, 26(5), 661–669. 10.1016/j.cub.2016.01.012

Russell, C., Wessnitzer, J., Young, J. M., Armstrong, J. D., & Webb, B. (2011). Dietary Salt Levels Affect Salt Preference and Learning in Larval Drosophila. PLoS ONE, 6(6), e20100. 10.1371/journal.pone.0020100

Saito, K., Hiramatsu, S., Watanabe, A., Wu, H., Ichinose, T., Yamagata, N., & Tanimoto, H. (2026). Presynaptic computation of reward intensities through the dual autoreceptor system. Current Biology, 36(9), 2357–2366.e4. 10.1016/j.cub.2026.03.077

Sakai, T., & Kitamoto, T. (2006). Differential roles of two major brain structures, mushroom bodies and central complex, for *Drosophila* male courtship behavior. Journal of Neurobiology, 66(8), 821–834. 10.1002/neu.20262

Saumweber, T., Rohwedder, A., Schleyer, M., Eichler, K., Chen, Y., Aso, Y., Cardona, A., Eschbach, C., Kobler, O., Voigt, A., Durairaja, A., Mancini, N., Zlatic, M., Truman, J. W., Thum, A. S., & Gerber, B. (2018). Functional architecture of reward learning in mushroom body extrinsic neurons of larval Drosophila. Nature Communications, 9(1), 1104. 10.1038/s41467-018-03130-1

Sawin, E. P., Harris, L. R., Campos, A. R., & Sokolowski, M. B. (1994). Sensorimotor transformation from light reception to phototactic behavior inDrosophila larvae (Diptera: Drosophilidae). Journal of Insect Behavior, 7(4), 553–567. 10.1007/BF02025449

Schipanski, A., Yarali, A., Niewalda, T., & Gerber, B. (2008). Behavioral Analyses of Sugar Processing in Choice, Feeding, and Learning in Larval Drosophila. Chemical Senses, 33(6), 563–573. 10.1093/chemse/bjn024

Schleyer, M., Weiglein, A., Thoener, J., Strauch, M., Hartenstein, V., Kantar Weigelt, M., Schuller, S., Saumweber, T., Eichler, K., Rohwedder, A., Merhof, D., Zlatic, M., Thum, A. S., & Gerber, B. (2020). Identification of Dopaminergic Neurons That Can Both Establish Associative Memory and Acutely Terminate Its Behavioral Expression. The Journal of Neuroscience, 40(31), 5990–6006. 10.1523/JNEUROSCI.0290-20.2020

Schroll, C., Riemensperger, T., Bucher, D., Ehmer, J., Völler, T., Erbguth, K., Gerber, B., Hendel, T., Nagel, G., Buchner, E., & Fiala, A. (2006). Light-Induced Activation of Distinct Modulatory Neurons Triggers Appetitive or Aversive Learning in Drosophila Larvae. Current Biology, 16(17), 1741–1747. 10.1016/j.cub.2006.07.023

Schulze, A., Gomez-Marin, A., Rajendran, V. G., Lott, G., Musy, M., Ahammad, P., Deogade, A., Sharpe, J., Riedl, J., Jarriault, D., Trautman, E. T., Werner, C., Venkadesan, M., Druckmann, S., Jayaraman, V., & Louis, M. (2015). Dynamical feature extraction at the sensory periphery guides chemotaxis. eLife, 4, e06694. 10.7554/eLife.06694

Selcho, M., Pauls, D., Huser, A., Stocker, R. F., & Thum, A. S. (2014). Characterization of the octopaminergic and tyraminergic neurons in the central brain of *Drosophila* larvae. Journal of Comparative Neurology, 522(15), 3485–3500. 10.1002/cne.23616

Tastekin, I., Khandelwal, A., Tadres, D., Fessner, N. D., Truman, J. W., Zlatic, M., Cardona, A., & Louis, M. (2018). Sensorimotor pathway controlling stopping behavior during chemotaxis in the Drosophila melanogaster larva. eLife, 7, e38740. 10.7554/eLife.38740

Thum, A. S., & Gerber, B. (2019). Connectomics and function of a memory network: The mushroom body of larval Drosophila. Current Opinion in Neurobiology, 54, 146–154. 10.1016/j.conb.2018.10.007

Villar, M. E., Pavão-Delgado, M., Amigo, M., Jacob, P. F., Merabet, N., Pinot, A., Perry, S. A., Waddell, S., & Perisse, E. (2022). Differential coding of absolute and relative aversive value in the Drosophila brain. Current Biology, 32(21), 4576–4592.e5. 10.1016/j.cub.2022.08.058

Vogt, K. (2020). Towards a functional connectome in *Drosophila*. Journal of Neurogenetics, 34(1), 156–161. 10.1080/01677063.2020.1712598

Vogt, K., Zimmerman, D. M., Schlichting, M., Hernandez-Nunez, L., Qin, S., Malacon, K., Rosbash, M., Pehlevan, C., Cardona, A., & Samuel, A. D. T. (2021). Internal state configures olfactory behavior and early sensory processing in *Drosophila* larvae. Science Advances, 7(1), eabd6900. 10.1126/sciadv.abd6900

Vosshall, L. B., & Stocker, R. F. (2007). Molecular Architecture of Smell and Taste in *Drosophila*. Annual Review of Neuroscience, 30(1), 505–533. 10.1146/annurev.neuro.30.051606.094306

Walter, T., & Couzin, I. D. (2021). TRex, a fast multi-animal tracking system with markerless identification, and 2D estimation of posture and visual fields. eLife, 10, e64000. 10.7554/eLife.64000

Weber, D., Vogt, K., Miroschnikow, A., Pankratz, M. J., & Thum, A. S. (2025). Four individually identified paired dopamine neurons signal taste punishment in larval Drosophila. eLife, 12, RP91387. 10.7554/eLife.91387.4

Widmann, A., Artinger, M., Biesinger, L., Boepple, K., Peters, C., Schlechter, J., Selcho, M., & Thum, A. S. (2016). Genetic Dissection of Aversive Associative Olfactory Learning and Memory in Drosophila Larvae. PLOS Genetics, 12(10), e1006378. 10.1371/journal.pgen.1006378

Winding, M., Pedigo, B. D., Barnes, C. L., Patsolic, H. G., Park, Y., Kazimiers, T., Fushiki, A., Andrade, I. V., Khandelwal, A., Valdes-Aleman, J., Li, F., Randel, N., Barsotti, E., Correia, A., Fetter, R. D., Hartenstein, V., Priebe, C. E., Vogelstein, J. T., Cardona, A., & Zlatic, M. (2023). The connectome of an insect brain. Science, 379(6636), eadd9330. 10.1126/science.add9330

Wolf, S., Dubreuil, A. M., Bertoni, T., Böhm, U. L., Bormuth, V., Candelier, R., Karpenko, S., Hildebrand, D. G. C., Bianco, I. H., Monasson, R., & Debrégeas, G. (2017). Sensorimotor computation underlying phototaxis in zebrafish. Nature Communications, 8(1), 651. 10.1038/s41467-017-00310-3

Wong, J. Y. H., Wan, B. A., Bland, T., Montagnese, M., McLachlan, A. D., O’Kane, C. J., Zhang, S. W., & Masuda-Nakagawa, L. M. (2021). Octopaminergic neurons have multiple targets in *Drosophila* larval mushroom body calyx and can modulate behavioral odor discrimination. Learning & Memory, 28(2), 53–71. 10.1101/lm.052159.120

Wosniack, M. E., Festa, D., Hu, N., Gjorgjieva, J., & Berni, J. (2022). Adaptation of Drosophila larva foraging in response to changes in food resources. eLife, 11, e75826. 10.7554/eLife.75826

Wu, G., Bazer, F. W., Dai, Z., Li, D., Wang, J., & Wu, Z. (2014). Amino Acid Nutrition in Animals: Protein Synthesis and Beyond. Annual Review of Animal Biosciences, 2(1), 387–417. 10.1146/annurev-animal-022513-114113

Zhang, Y. V., Ni, J., & Montell, C. (2013). The Molecular Basis for Attractive Salt-Taste Coding in *Drosophila*. Science, 340(6138), 1334–1338. 10.1126/science.1234133

